# Influence of Alzheimer’s disease related neuropathology on local microenvironment gene expression in the human inferior temporal cortex

**DOI:** 10.1101/2023.04.20.537710

**Authors:** Sang Ho Kwon, Sowmya Parthiban, Madhavi Tippani, Heena R. Divecha, Nicholas J. Eagles, Jashandeep S. Lobana, Stephen R. Williams, Michelle Mak, Rahul A. Bharadwaj, Joel E. Kleinman, Thomas M. Hyde, Stephanie C. Page, Stephanie C. Hicks, Keri Martinowich, Kristen R. Maynard, Leonardo Collado-Torres

**Affiliations:** The Biochemistry, Cellular, and Molecular Biology Graduate Program, Johns Hopkins School of Medicine, Baltimore, MD, USA; Lieber Institute for Brain Development, Johns Hopkins Medical Campus, Baltimore, MD, USA; The Solomon H. Snyder Department of Neuroscience, Johns Hopkins School of Medicine, Baltimore, MD, USA; Department of Biostatistics, Johns Hopkins Bloomberg School of Public Health, Baltimore, MD, USA; 10x Genomics, Pleasanton, CA, USA; Department of Psychiatry and Behavioral Sciences, Johns Hopkins School of Medicine, Baltimore, MD, USA; Department of Neurology, Johns Hopkins School of Medicine, Baltimore, MD, USA; Malone Center for Engineering in Healthcare, Johns Hopkins University, Baltimore, MD, USA; The Kavli Neuroscience Discovery Institute, Johns Hopkins University, Baltimore, MD, USA

**Author notes:** Equal Contribution (Sowmya Parthiban, Madhavi Tippani). Corresponding authors, Stephanie C. Hicks, Keri Martinowich Kristen R. Maynard Leonardo Collado-Torres.

**Keywords:** Spatially-resolved transcriptomics, RNA-protein co-detection, neuropathology, Alzheimer’s disease, amyloid-β (Aβ), phosphorylated tau (pTau)

## Abstract

Neuropathological lesions in the brains of individuals affected with neurodegenerative disorders are hypothesized to trigger molecular and cellular processes that disturb homeostasis of local microenvironments. Here, we applied the 10x Genomics Visium Spatial Proteogenomics (Visium-SPG) platform, which measures spatial gene expression coupled with immunofluorescence protein co-detection, in post-mortem human brain tissue from individuals with late-stage Alzheimer’s disease (AD) to investigate changes in spatial gene expression with respect to amyloid-β (Aβ) and hyperphosphorylated tau (pTau) pathology. We identified Aβ-associated transcriptomic signatures in the human inferior temporal cortex (ITC) during late-stage AD, which we further investigated at cellular resolution with combined immunofluorescence and single molecule fluorescent in situ hybridization (smFISH) co-detection technology. We present a workflow for analysis of Visium-SPG data and demonstrate the power of multi-omic profiling to identify spatially-localized changes in molecular dynamics that are linked to pathology in human brain disease. We provide the scientific community with web-based, interactive resources to access the datasets of the spatially resolved AD-related transcriptomes at https://research.libd.org/Visium_SPG_AD/.

## Introduction

Pathological signaling cascades that alter homeostasis in brain microenvironments occur in various disorders including stroke, infection, tumor, and neurodegenerative diseases, as well as in response to traumatic brain injuries (1–6). The molecular and cellular landscape of the local tissue microenvironment is altered via release of cytokines, chemokines, and other secreted proteins, such as interleukin-1β (IL-1β), C-C motif chemokine ligand 2 (CCL2), and reactive oxygen species (ROS), which triggers downstream signaling cascades via second messenger molecules (7–11). Pathological lesions often exhibit remarkable features such as apoptosis, necrosis, neuroinflammation or changes in cell morphology that can be detected with histological or immunohistochemical staining of post-mortem brain tissue. For example, Alzheimer’s disease (AD) is characterized by two major neuropathologies - extracellular deposition of amyloid-beta (Aβ) peptides that form senile plaques, and intracellular inclusions of hyperphosphorylated tau (pTau) proteins that form various neurofibrillary elements, including tangles (12–14). The progressive accumulation of Aβ and pTau is hypothesized to functionally contribute to cognitive decline and dementia, and to contribute to cell death during disease progression (12,15). However, the pathophysiological mechanisms rendering Aβ and pTau neurotoxic are still not fully understood (16). Identifying how plaques and tangles contribute to molecular and cellular signaling cascades that trigger cell death in the local microenvironment is essential to develop novel approaches for preventing or reversing disease progression in AD.

Several studies have now deployed spatially-resolved transcriptomics (SRT) to study spatial gene expression in mouse models of AD (17,18). However, only humans and some non-human primates naturally develop plaques, tangles and dementia, and hence rodent models cannot fully recapitulate neurodegenerative diseases in humans (19,20). Therefore, investigative studies in human brain tissue from donors diagnosed with AD that carry plaques and tangles are indispensable to study the molecular sequelae associated with Aβ and pTau deposition. To better understand how Aβ deposition and pTau accumulation in the human brain impacts molecular signaling in the local tissue environment in late-stage AD, we utilized spatial profiling coupled with multiplex immunofluorescence using the 10x Genomics Visium Spatial Proteogenomics platform (Visium-SPG) to generate a proteomic-based, spatially-resolved, transcriptome-scale map of the human inferior temporal cortex (ITC), a region which displays reduced cortical thickness in AD (21–23). By combining proteomic and transcriptomic information, Visium-SPG allows for quantification of transcriptional dynamics with respect to spatially-localized pathology. Here, we evaluated the utility of the Visium-SPG platform by testing the hypothesis that local tissue microenvironments in close proximity to AD-related neuropathology have distinct molecular signatures. We first immunostained post-mortem human brain tissue sections from donors diagnosed with AD for Aβ and pTau, and then conducted SRT using the Visium-SPG platform. Integration of proteomic and transcriptomic data allowed us to measure gene expression profiles within local tissue microenvironments that were spatially defined by the presence of pathology. Our multi-omics approach identified genes differentially expressed by proximity to Aβ, which we further validated at cellular resolution using RNAscope smFISH combined with Aβ immunodetection. To make these data easily available to the neuroscience community, we generated user-friendly web resources to further explore the dataset.

## Results

### Experimental design and study overview

To identify local gene expression profiles in the tissue microenvironment with respect to Aβ and pTau deposition, we employed the 10x Genomics Visium-SPG technology in tissue sections from the ITC of 4 human brain donors (**Figure 1A** and **Table S1**). Three donors were clinically diagnosed with AD (Braak V-VI/CERAD Frequent), and 1 donor was an age-matched neurotypical control. Tissue blocks containing the ITC (**Figure S1A**) were collected onto a Visium spatial gene expression slide with 4 capture arrays, where each array contains ∼5,000 expression spots bearing spatially-barcoded oligonucleotide probes (24). For each donor, we performed 2-3 spatial replicates, resulting in a total of 10 samples. Following collection onto capture arrays, tissue sections were stained using monoclonal primary antibodies, 6E10 (25) and AH36 (26,27) to label Aβ and pTau, respectively. Immediately following staining, the entire slide was imaged and then processed for segmentation of Aβ and pTau signals against the confounding noises and artifacts due to lipofuscin and particulates of ribonucleoside vanadyl complex (RVC) ribonuclease inhibitor (**Figure S2** and **Methods**). We confirmed that the staining protocol of Visium-SPG detected Aβ plaques and various forms of neurofibrillary elements, including tangles (**Figure 1B**). Immunostaining and image processing workflows were validated for specificity using images from neurotypical control samples, which largely lacked pathology (**Figure S3**). Following image acquisition, tissue sections underwent permeabilization (**Figure S1B**), followed by on-slide cDNA synthesis, amplification and subsequent library construction. The resulting transcriptomic data from the sequenced libraries was computationally aligned with the immunofluorescence image data to integrate transcriptomic and proteomic information for each tissue sample. We confirmed the spatial orientation of white and gray matter in each tissue section by visualizing marker genes for oligodendrocytes (*MOBP)* and neurons (*SNAP25)* (**Figure 1C**, **Figure S3B**, **Figure S4**, and **Figure S5**). The final image segmentation of Aβ and pTau was used to classify Visium spots into different AD-related neuropathological categories. Following Visium spot classification, we constructed spatial maps of gene expression by analyzing transcriptomic data with respect to AD-related pathology categories (**Figure 1D** and **Figure S6**).

**Fig.1.**
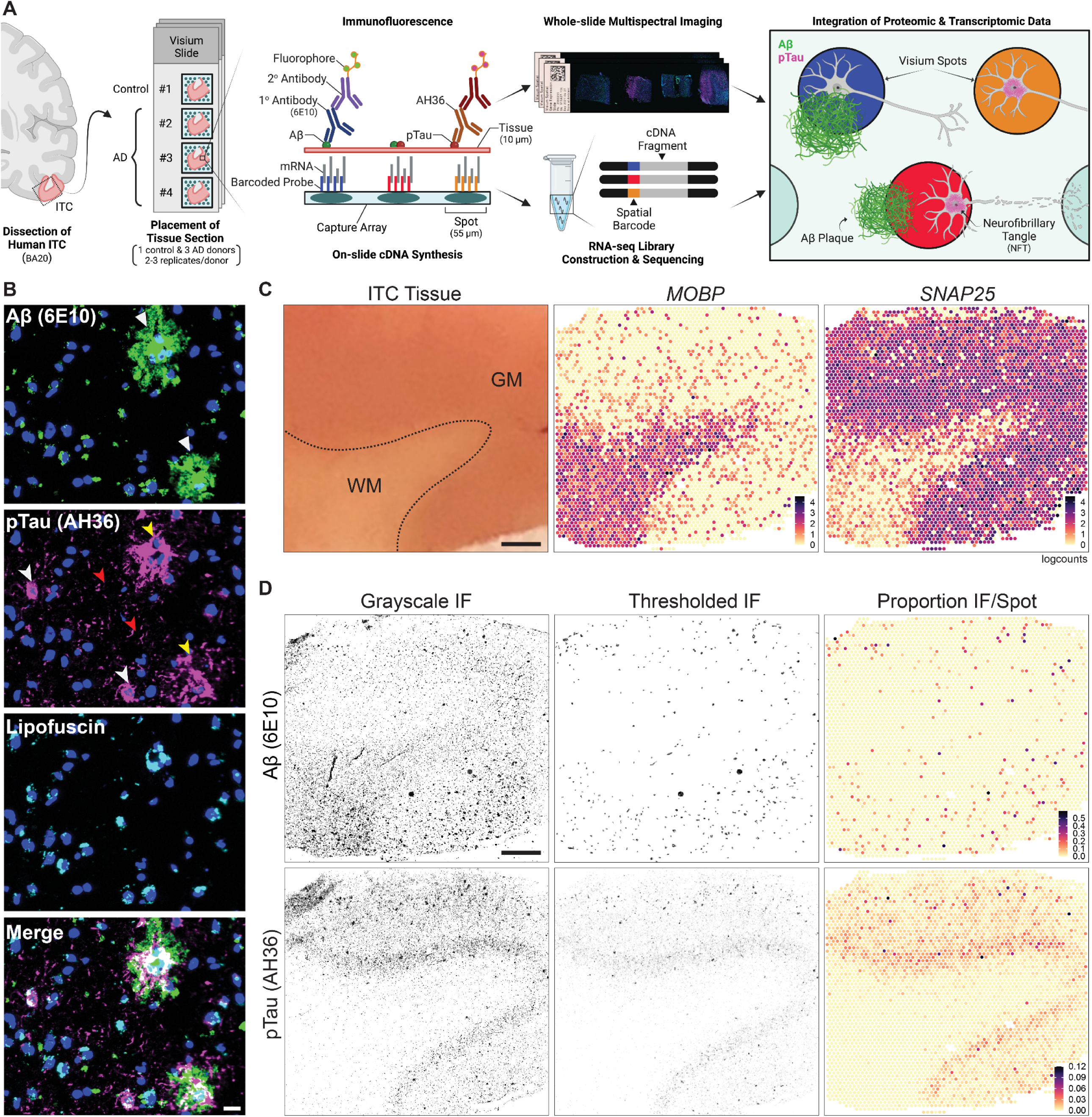
Spatial transcriptomics combined with immunodetection of Aβ and pTau in the human inferior temporal cortex (ITC). (**A**) Schematic of experimental design using Visium Spatial Proteogenomics (Visium-SPG) to investigate the impact of Aβ and pTau aggregates on the local microenvironment transcriptome in the post-mortem human brain. Human ITC blocks were acquired from 3 donors with AD and 1 age-matched neurotypical control. Tissue blocks were cryosectioned at 10μm to obtain 2-3 replicates per donor and sections were collected onto individual capture arrays of a Visium spatial gene expression slide, yielding a total of 3 gene expression experiments. The entire slide (4 tissue sections) was stained and scanned using multispectral imaging methods to detect Aβ and pTau immunofluorescence (IF) signals as well as autofluorescence. Following imaging, tissue sections were permeabilized and subjected to on-slide cDNA synthesis after which libraries were generated and sequenced. Transcriptomic data was aligned with the respective IF image data to generate gene expression maps of the local transcriptome with respect to Aβ plaques and pTau elements, including neurofibrillary tangles. (**B**) High magnification images show Aβ plaques (white triangles) and various neurofibrillary elements such as tangles (white arrowheads), neuropil threads (red arrowheads), and neuritic tau plaques (yellow arrowheads). Lipofuscin (cyan) was identified through spectral unmixing and pixels confounded with this autofluorescent signal were excluded from analysis, scale bar, 20μm. (**C**) ITC tissue block from Br3880 (left) and corresponding spotplots (right) from the Visium data show gene expression of *MOBP* and *SNAP25,* which demarcates the border between gray matter (GM) and white matter (WM), scale bar, 1mm. Color scale indicates spot-level gene expression in logcounts. (**D**) Image processing and quantification of Aβ and pTau per Visium spot. Aβ and pTau signals were thresholded in their single IF channels for segmentation against autofluorescence background, including lipofuscin. Thresholded Aβ and pTau signals were aligned to the gene expression map of the same tissue section from Br3880 and quantified as the proportion of number of pixels per Visium spot, which is visualized in a spotplot, scale bar, 1mm.

### Gene expression profiling of human brain microenvironments with AD-related neuropathology

While imaging of tissue immunostained for Aβ and pTau revealed diverse spatial patterns of AD pathology in the human ITC (**Figure 1B**), classification of these pathologies is complicated due to the existence of multiple variations of Aβ plaques (e.g. diffuse versus neuritic) and neurofibrillary elements (e.g. tangles versus neuropil threads) (28,29). To simplify classifications for subsequent analysis, we first categorized expression spots based on the presence or absence of Aβ or pTau (**Figure 2A**). This resulted in 7 broad subtypes of spots: 1) containing no Aβ and no pTau, 2) containing only Aβ, 3) containing only pTau, 4) containing both Aβ and pTau, 5) directly adjacent to Aβ spots, 6) directly adjacent to pTau spots, and 7) directly adjacent to spots containing both Aβ and pTau. The spot categories from 2 to 4 (Aβ, pTau, both) and from 5 to 7 (next_Aβ, next_pTau, next_both) each represent the core and periphery of the brain microenvironment associated with Aβ, pTau, and the combination of the two. Of note, Aβ plaques and different neurofibrillary forms of pTau varied in shape, number, and size within a donor across the different replicates (**Figure S7A**). Given the heterogeneous appearance and variability in pathological load within individual Visium spots, we focused on Visium spots containing relatively high burdens of neuropathology. We selected spots with robust pathology in the AD samples by thresholding for the number and size of plaques and/or neurofibrillary elements. To do this, we first identified the regions of interest (ROIs) to indicate plaques and tangles as ROI objects. We set a baseline cut-off value in the ROI object counts (number) and pixel-based proportion (size) based on what was observed in the neurotypical control samples. Spots above this threshold were then classified according to binary expression of Aβ and/or pTau (**Figure S7B-C**).

**Fig.2.**
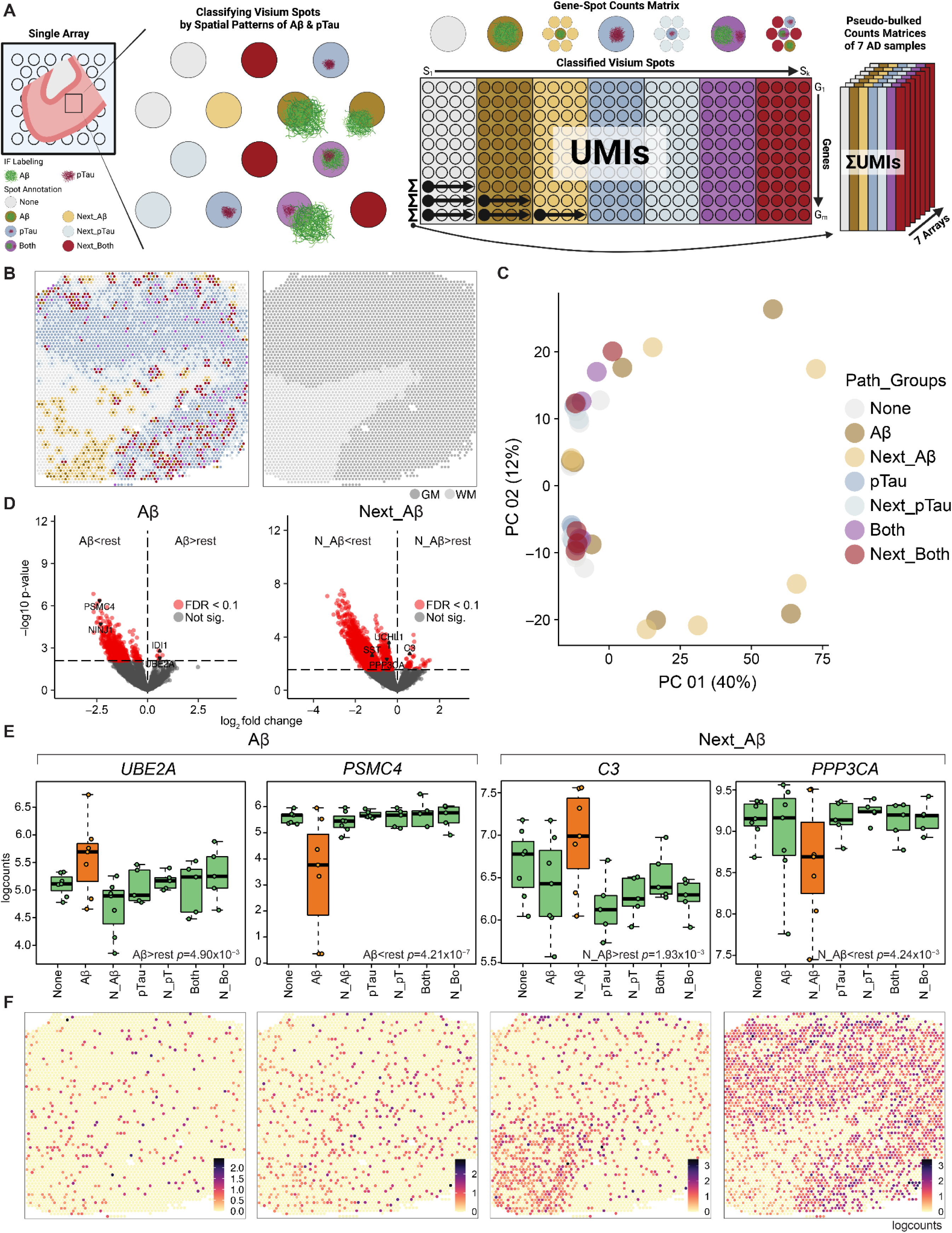
Identification of transcriptomic signatures in local microenvironments harboring AD-related neuropathology. (**A**) Graphical overview of spot-level annotation for AD-related neuropathology (left), and strategy for pseudo-bulking annotated spots (right). Aβ and pTau labeling was quantified in individual spots, which were subsequently annotated accordingly to reconstruct spatial heterogeneity of pathology according to the following hierarchy: spots containing both Aβ and pTau (purple), Aβ pathology (brown), pTau pathology (blue), and then spots adjacent to both Aβ and pTau (red), Aβ pathology (light brown), and pTau pathology (light blue). Spots with no significant Aβ and pTau deposition were labeled white. Annotated spots were then collapsed into the 7 pathological categories in a gene x spot matrix for each tissue sample. Unique molecular identifier (UMI) counts for each gene were pseudo-bulked within an individual pathological category across the 7 replicate arrays from 3 AD donors to yield 49 pathology-enriched gene expression profiles. (**B**) Spotplot for Br3880 showing spot-level annotation of AD-related neuropathology (left) and unsupervised clustering of spots using *BayesSpace* (right). Spots associated with pathology were aggregated within gray matter (GM; dark gray), which was identified using an unsupervised clustering approach, as in **Figure S11A**. Any pathology-associated spots in the white matter (WM; light gray) were not included in downstream analyses. (**C**) PCA was performed on the 41 pseudo-bulked gene expression profiles across the 7 spatial categories of AD-related neuropathology. PC1 separated the Aβ and next_Aβ spots from the other pathological categories. (**D**) Volcano plots depicting differentially expressed genes (DEGs) between Aβ pathology (Aβ) or adjacent Aβ pathology (Next_Aβ) versus the rest of pathological categories. Each dot represents a gene, plotted with its log2 fold change (x-axis) and -log10 *p*-value (y-axis), thus comparing the effect size (fold change) against the statistical evidence for differential expression (*p*-value). The dashed line represents a *p*-value threshold matching FDR<0.1, with DEGs below the line considered not significant (gray, Not sig.) Significant DEGs that passed the FDR threshold were labeled red and classified by their enrichment (>rest) and depletion (<rest) in gene expression according to the ‘enrichment’ model. (**E**) Boxplots of DEGs in Aβ pathology spots (*UBE2A* and *PSMC4)* and adjacent Aβ pathology spots (*C3* and *PPP3CA)* depicted in orange across all samples, compared to all other pathological categories depicted in green. For brevity, ‘Next’ was shortened to N. (**F**) Corresponding spotplots of *UBE2A*, *PSMC4*, *C3*, and *PPP3CA* for Br3880 showing spatial distribution of each DEGs. Color scale indicates spot-level gene expression in logcounts.

To address the possibility of synergistic effects of Aβ and pTau co-expression (30–32) as well as the extracellular effects of Aβ peptides (33,34), we prioritized spots containing both Aβ and pTau (both), followed by spots containing Aβ pathology (Aβ), pTau pathology (pTau), and then spots adjacent to Aβ and pTau (next_both; n_bo), Aβ pathology (next_Aβ; n_Aβ), and pTau pathology (next_pTau; n_pT). Spots with no significant pathology were labeled as ‘none’. This hierarchical classification of the 7 spot categories allowed for inclusion of a sufficient number of spots for the ‘both’ category (**Figure S8** and **Methods**). We then calculated a number of quality control (QC) metrics, identifying low-quality spots due to low library size, low number of expressed genes, or high proportions of mitochondrial reads and only dropped low library size spots on the tissue edges (**Figure S9** and **Methods**). Since AD neuropathology is most abundant in the cortical laminae of the human cortex (35–37), we focused our analysis on spots located in the gray matter (**Figure 2B**). To identify the gray matter using a data-driven approach, we first applied batch correction to the 10 tissue samples using *Harmony* (38) **(Figure S10**), and then employed *BayesSpace* (39), an algorithm for unsupervised spatial clustering. At cluster number *k*=2, we identified spots localizing to the gray and white matter (**Figure S11A**). We verified the identity of *BayesSpace* clusters by spatially registering (40) these spatial domains to manually annotated cortical layers and white matter from our previously published work in the dorsolateral prefrontal cortex (DLPFC) (24) (**Figure S11B**). *BayesSpace* cluster 1 was enriched for genes expressed in the gray matter while *BayesSpace* cluster 2 was enriched for genes expressed in the white matter. As expected, the gray matter-associated *BayesSpace* cluster 1 was enriched with numerous pathology-containing spots associated with Aβ and pTau pathology (**Figure 2B** and **Figure S8)**. Therefore, focusing on the gray matter only, we next pseudo-bulked the unique molecular identifier (UMI) counts for each gene within each individual pathological category across each AD sample. This generated a total of 41 pathology-enriched gene expression profiles across the 7 AD samples (**Figure 2A**) with at least 15 spots per combination, out of the possible 49 gene expression profiles.

Principal component analysis (PCA) revealed that a significant source of biological variation in these 41 pathology-enriched profiles was associated with pathological categories, particularly between Aβ-related categories against the others (**Figure 2C**). To study changes in gene expression associated with these pathological categories, we identified differentially expressed genes (DEGs) between the 7 pathological categories using linear mixed effects modeling. In particular, we used an ‘enrichment’ model (24,40) in which one pathological category was compared to all other categories. Using this strategy, we identified 9 up-regulated and 687 down-regulated DEGs for the Aβ category, and 51 up-regulated and 2,319 down-regulated DEGs for the next_Aβ category, all with a false discovery rate (FDR) of less than 0.1 (**Figure 2D**, **Table S2**, and **Table S3**). No significant DEGs were identified in the remaining 5 categories (**Figure S12A**). Gene expression levels of representative DEGs for the Aβ and next_Aβ category illustrate their degree of enrichment and depletion (**Figure 2E**). The corresponding spatial gene expression patterns are also depicted in the spot plots (**Figure 2F**). We identified two DEGs implicated in the ubiquitin-proteasome pathway, suggesting dysregulated protein degradation around Aβ plaques. Ubiquitin conjugating enzyme E2 A (*UBE2A*) was enriched within the Aβ category compared to all other spot categories (*p*=4.90∗10^-3^), while 26S proteasome AAA-ATPase subunit RPT3 (*PSMC4*) was depleted (*p*=4.21∗10^-7^). We also identified complement component 3 (*C3*) and protein phosphatase 3 catalytic subunit alpha (*PPP3CA*) as differentially expressed within the next_Aβ category. Within the gray matter, *C3* was enriched in the next_Aβ category compared to all other categories (*p*=1.93∗10^-3^), while *PPP3CA* was depleted (*p*=4.24∗10^-3^).

To investigate the gene expression profile of the extended Aβ-associated microenvironment spanning both Aβ and next_Aβ spots, we collapsed the Aβ and next_Aβ spots into a single category and ran differential expression testing compared to the other 5 pathological categories using the same enrichment model (**Table S4**). Based on an exploratory pathway analysis, several DEGs involved in apoptosis, JAK-STAT signaling pathways, inflammation, and cellular response to stress were enriched in the extended Aβ-associated microenvironment. These included genes (**Figure S12B**) encoding the TNF receptor superfamily member 1A (*TNFRSF1A*; *p*=8.18∗10^-4^) (41–43), cytochrome P450 family 1 subfamily B member 1 (*CYP1B1*; *p*=1.37∗10^-3^) (44,45), zinc finger protein 385A (*ZNF385A*; *p*=1.06∗10^-4^) (46), RIO kinase 3 (*RIOK3*; *p*=1.08∗10^-3^) (47), metallothionein 1F (*MT1F*; *p*=5.00∗10^-4^) (48), and *s*ushi domain containing 6 (*SUSD6*; *p*=2.68∗10^-5^) (49), all of which have been recently reported as being associated with AD in human and mouse genomic and transcriptomic studies (50–55). We also identified several depleted DEGs, such as autophagy related 5 (*ATG5*; *p*=9.87∗10^-4^), DnaJ heat shock protein family Hsp40 member A3 (*DNAJA3*; *p*=3.21∗10^-4^), ubiquitin protein ligase E3 component N-recognin 3 (*UBR3*; *p*=1.26∗10^-3^), ubiquitin like modifier activating enzyme 6 (*UBA6*; *p*=7.81∗10^-4^), cyclin dependent kinase 5 (*CDK5*; *p*=2.41∗10^-4^), and SH3 and multiple ankyrin repeat domains 3 (*SHANK3*; *p*=7.26∗10^-5^) (**Figure S12C**). These genes are associated with autophagy (56,57), protein ubiquitination (58), and synaptic plasticity (59,60), which are well-established biological functions associated with AD. Of note, *ZNF385A* (*p*=8.82∗10^-1^ for next_Aβ) and *SUSD6* (*p*=2.94∗10^-2^ for next_Aβ) were not previously identified as statistically significant DEGs in any of the 7 spatially resolved pathological categories when Aβ and next_Aβ spots were not collapsed (**Figure S12D**). In summary, using Visium-SPG, we identified a set of genes differentially expressed in the core (Aβ spots) and peripheral (next_Aβ spots) compartments, as well as in the extended Aβ-associated microenvironment spanning the two compartments. Our analysis suggests the potential involvement of stress responses linked to protein degradation and inflammation, as well as regulation of synaptic plasticity in the Aβ-associated microenvironment.

Next, to add clinical relevance to our integrated spatial gene expression and pathology maps, we performed multi-marker analysis of genomic annotation (MAGMA) (61) to evaluate whether genes associated with genetic risk for AD (62), frontotemporal dementia (FTD) (63), and Parkinson’s disease (PD) (64) genome-wide association studies (GWAS) are linked to DEGs from the Aβ-associated microenvironment. We found that spots classified as next_Aβ, which represent the peripheral compartments of Aβ-associated microenvironments, harbored a substantial genetic risk for AD and PD (**Figure S13A**). Using gene set enrichment analyses, we further evaluated whether different Aβ-associated microenvironments were enriched and/or depleted in 1) the co-expression gene module M109 associated with both cognitive decline and Aβ pathology in AD based on an bulk RNA sequencing (bulk RNA-seq) data from the ROSMAP cohort by Mostafavi et al (65) or 2) the cell type-specific DEGs in the entorhinal cortex between AD donors and controls from a single nucleus RNA sequencing (snRNA-seq) study by Grubman et al (66). We found no statistically significant (*p*<0.05) enrichment with bulk RNA-seq (**Figure S13B**) but we noted that genes differentially expressed in microglia between AD donors and controls were enriched in the periphery of the Aβ-associated microenvironments, suggesting an important role for microglia in pathological processes in the late stage of AD (*p*=0.047) (**Figure S13C**).

### Spatial gene expression of Aβ-associated microenvironments at single cell resolution

The resolution of the Visium platform is limited by the size and spacing of spots across the array. First, the size of the expression spot (55µm diameter) does not allow for cellular or sub-cellular resolution. In the present study, we observed an average of 4.5 nuclei per spot, which is consistent with our previous study of the human DLPFC where we observed an average of 3.3 cell bodies per spot (24). In addition, since spots are spaced in a honeycomb pattern with 100µm distance center-to-center between neighboring spots, transcriptomic data is not captured in the 45µm of inter-spot space (67). To add cellular resolution to our Visium data and further explore the expression of promising DEGs related to AD pathology, we used RNAscope smFISH combined with immunofluorescence (RNA-Protein co-detection workflow; FISH-IF) to define transcript expression in single cells located in the vicinity of Aβ plaques (**Figure 3A**).

**Fig.3.**
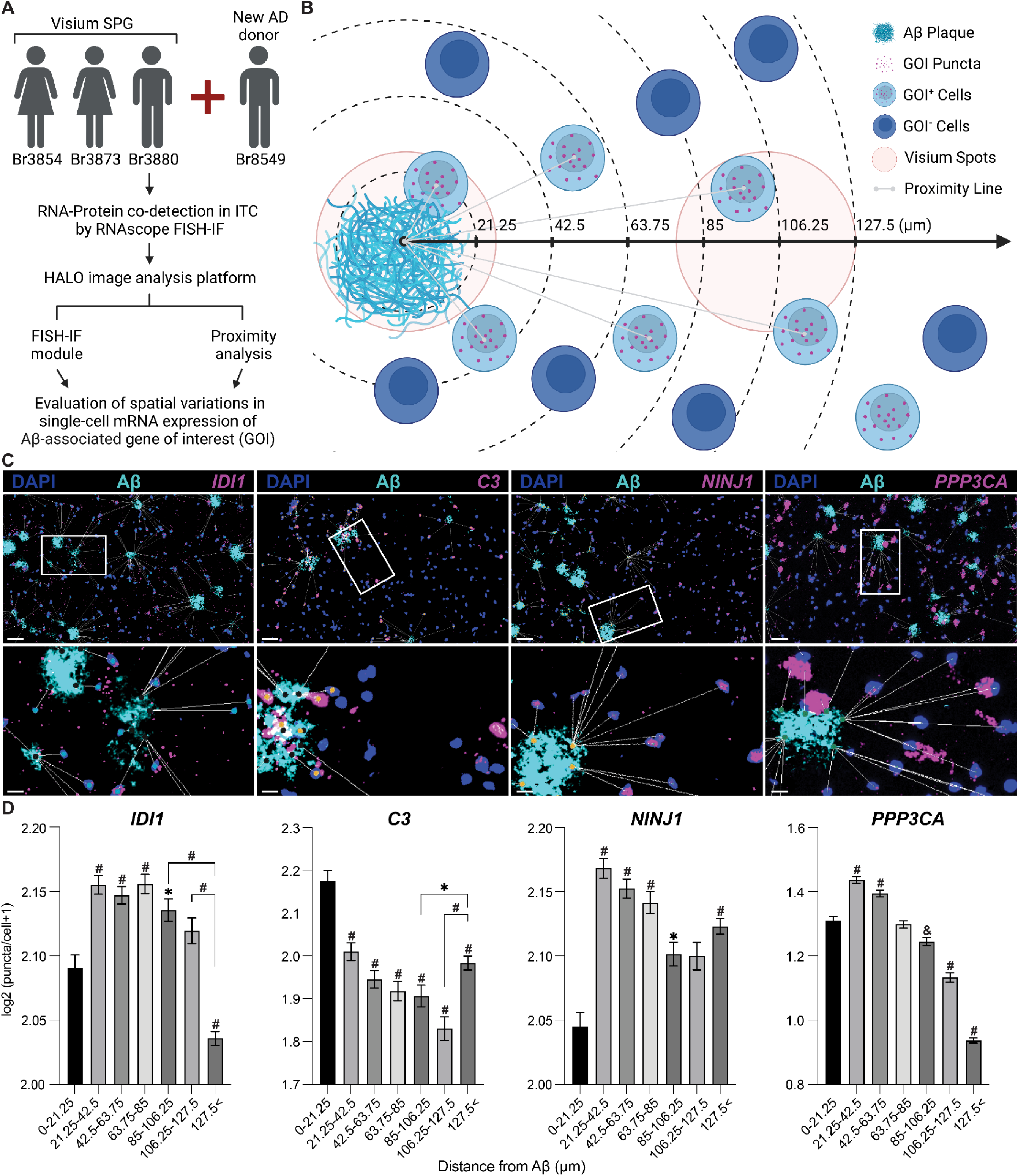
Characterization of Aβ-associated transcriptional signatures at cellular resolution. (**A**) Flowchart of experimental design and data analysis. Human ITC tissues from 3 original AD donors plus additional male donor Br8549 were subjected to multiplexed staining using RNAscope smFISH combined with immunofluorescence (FISH-IF) to detect genes of interest (GOIs) and Aβ plaques. Images were analyzed with HALO image analysis software to assess spatial relationships between Aβ and cells expressing GOI. The FISH-IF module of HALO was used for image segmentation and quantification of Aβ and GOIs. The proximity analysis module was used to determine a distance between Aβ and cells expressing or not expressing GOIs. The outputs of the two modules were integrated to measure the gene expression of GOIs within a predefined proximity of Aβ at cellular resolution. (**B**) Schematic describing proximity analysis. An Aβ-associated microenvironment was demarcated by approximating the Visium spot grid-line system in which the center of a single Visium spot is 127.5µm away from its neighboring spot. This distance was further subdivided into 6 evenly spaced intervals, resulting in a total of 7 bins to finely resolve the spatial gene expression gradients of GOIs. The proximity between Aβ and nearby cells expressing and not expressing GOIs was measured and used to classify into the 7 bins for quantifying the average GOI gene expression. (**C**) RNA-protein co-detection of Aβ and *IDI1*, *C3*, *NINJ1*, *PPP3CA* reveals the spatial distribution patterns of Aβ (cyan) and GOIs (magenta) at lower (Top, scale bar: 50µm) and higher magnifications (Bottom, scale bar: 12.5µm). Proximity lines indicate the distance between Aβ and nearby cells expressing GOIs (max: 127.5µm). (**D**) Bar plots show quantification of gene expression levels for GOIs in **Figure 3C** across 7 consecutive bins representing increased distance from Aβ, as modeled in **Figure 3B**. Gene expression levels were determined with log2 (X+1) transformation where X represents the counts of puncta in a single cell for a given GOI. Data are mean ± SEM. The first bin was compared to all the rest by default for statistical tests (Kruskal-Wallis test, \**p*<0.05, ^&^*p*<0.005, and ^#^*p*<0.0001). The bracket denotes statistical testing between two specified bins. Violin plots are provided in **Figure S16C** describing the cellular distribution and numbers counted for each bin.

In particular, we focused on differentially expressed genes in the Aβ and next_Aβ categories at FDR<0.1. We selected 6 DEGs for follow-up based on their biological significance in AD as well as availability for commercialized RNAscope probes. With Visium-SPG, we identified isopentenyl-diphosphate delta isomerase 1 (*IDI1*; *p*=1.80∗10^-3^ for Aβ>rest) and *C3* as enriched in the vicinity of Aβ plaques. We also identified nerve injury-induced protein 1 (*NINJ1*; *p*=1.87∗10^-5^ for Aβ<rest), *PPP3CA*, ubiquitin C-terminal hydrolase L1 (*UCHL1*; *p*=7.62∗10^-4^ for next_Aβ<rest), and somatostatin (*SST*; *p*=2.68∗10^-3^ for next_Aβ<rest) as depleted in Aβ-adjacent Visium spots (**Figure S14A**). We performed FISH for each selected gene while immunolabeling Aβ plaques in the same tissue sections. We then completed segmentation using HALO software (68–70) to identify Aβ plaques and surrounding cells expressing genes of interest (GOI), which we refer to as GOI expressors (**Figure 3B** and **Figure S15A**). The number of RNAscope puncta was quantified in GOI expressors using the FISH-IF module in HALO (**Figure 3B** and **Figure S15B-C**). After image segmentation and quantification, the proximity between Aβ and GOI expressors was analyzed using the spatial analysis module in HALO (**Figure S15D**). To understand the spatial relationship between Aβ plaques and adjacent cells, including GOI expressors, we approximated the Aβ-associated microenvironment used in Visium analyses by introducing a geometric layout based on the Visium spot grid line system. Specifically, we set a maximum proximity between segmented Aβ plaques and neighboring cells at 127.5µm, which is the radial distance from the center of a Visium spot containing Aβ to the opposite end of an adjacent plaque-free spot (**Figure 3B**). To finely resolve spatial gene expression gradients, we divided this approximated Aβ-associated microenvironment into 6 evenly-spaced, concentric rings for a total of 7 sub-regions in which GOI expressors were classified based on their location to neighboring plaques. This proximity analysis was performed for each tissue section per donor, and the spatially classified cells were pooled across the 4 AD donors. The cumulative number of cells measured in each sub-region varied between 1,615 and 26,789 (with an average median of 14,505) **(Figure S16C**), and the total number of segmented fragments of Aβ plaques (**Figure S15C**) per each tissue section ranged from 1,671 to 5,413 (with a mean of 3,359, a median of 3,659, and a standard deviation of 1,135).

Using this approach, we finely resolved changes in gene expression gradients identified with Visium-SPG within Aβ-associated microenvironment at cellular resolution (**Figure 3C**-**D**). Consistent with Visium data, *IDI1* was lowly expressed in the area closest to Aβ plaques (0-21.25µm), but was significantly increased in the region extending to 106.25µm, at which point expression decreased, reaching its lowest expression levels in the regions farthest from plaques (>127.5µm). *C3* expression peaked in regions directly adjacent to plaques, followed by a downward trajectory and then a second peak beyond the defined local microenvironment at 127.5µm (**Figure 3D** and **Figure S16D**). The lowest levels of *NINJ1* were observed close to plaques, and *PPP3CA* was slightly enriched in the inner microenvironment (21.25-63.75µm), but its progressive decrement was detected on the outskirts (>85µm). Similar to this declining trend, *UCHL1* and *SST* both exhibited statistically significant decreases in gene expression levels in the distal region of the microenvironment when the total cell population was counted as a normalization factor (**Figure S16A**). In summary, we defined spatial gene expression patterns of the Aβ-associated genes identified with Visium-SPG at cellular resolution using RNAscope FISH-IF to add further context to how their gene expression levels become spatially variable with respect to proximity to Aβ pathology.

## Discussion

Application of SRT to the study of human brain disorders (71,72) has rapidly increased due to the technology’s innovative ability to preserve spatial-anatomical context while assessing gene expression. Here, we demonstrate, to our knowledge, the first use case of 10x Genomics Visium-SPG in AD, which incorporates protein co-detection with SRT, in human post-mortem brain tissue to investigate the influence of AD-related neuropathology on local gene expression. The multi-omic approach of immunofluorescent (IF) staining followed by SRT in the Visium-SPG assay allows for simultaneous detection of mRNA with specific proteins of interest. This integrative approach can resolve local transcriptomic profiles of pathology-associated proteins to comprehensively assess the molecular impact of neuropathology on spatially-defined microenvironments. Simultaneous detection of mRNA and proteins on the same tissue section reduces the computational burden associated with aligning spatially adjacent tissue sections and their latent structural variations in tissue architecture (18,73–75).

Using Visium-SPG, we generated transcriptome-scale tissue atlases of human ITC harboring AD-related neuropathology. The ITC is a cortical structure that succumbs to progressive accumulation of Aβ and pTau in AD (76,77). We focused on late-stage AD (Braak V-VI/CERAD Frequent) to define the biological consequences of severe pathology. Consistent with prior clinicopathological examinations, our IF staining revealed heavily intertwined Aβ and pTau pathology, which spanned various cellular and subcellular compartments. Diffuse and neuritic plaques were observed throughout the tissue sections, along with a variety of neurofibrillary elements including neuropil threads and tangles. To investigate dynamic transcriptional patterns associated with Aβ and pTau depositions, we computationally registered the IF image data with SRT data from the same tissue section and aligned Aβ and pTau protein signals with spatially resolved transcriptomics. AD neuropathology within Visium expression spots was classified using a binary categorization based on the absence or presence of protein signal. We subsequently classified spots according to a pathology hierarchy that prioritized the ‘both’ pathology as we were most interested in potential synergies due to co-localization or proximity between Aβ and pTau. Alternative hierarchies prioritizing either Aβ or pTau did not result in the identification of many ‘both’ spots due to the categorization into Aβ or pTau only. Pathological maps were generated by measuring the number of ROI objects and proportion of pixels within the ROIs per spot to estimate local pathological burden. We used pseudo-bulking and differential gene expression analyses to investigate molecular changes in microenvironments with a substantial pathological burden of Aβ, pTau or both, as well as their neighboring microenvironments (next_Aβ, next_pTau, and next_both), and microenvironments without significant pathology (none). By integrating spatial transcriptomic data with pathology maps derived from immunofluorescence, we identified differentially expressed genes (DEGs) between the core (Aβ) or periphery (next_Aβ) of the Aβ-associated microenvironments against the other six pathological categories. These DEGs included *UBE2A*, *PSMC4*, *C3*, and *PPP3CA*. Additionally, we performed a sensitivity analysis by combining Aβ and next_Aβ spots into a single category to describe the extended Aβ microenvironment (Aβ_env) that allowed us to identify new Aβ-associated DEGs, such as *ZNF385A* and *SUSD6*. These genes were not initially identified when Aβ and next_Aβ spots were analyzed separately, likely due to their similar gene expression patterns between Aβ and next_Aβ spots. The DEGs we identified suggest a handful of biological processes and signaling networks that may contribute to the neurodegenerative and proinflammatory conditions in the vicinity of Aβ plaques. Among them were the cell survival/death signaling pathways (e.g., *CYP1B1*, *CDK5*, *TNFRSF1A*, *ZNF385A*, *BCL2*, *BID*, and *AIFM1*), the complement cascade system (e.g., *C3*, *C1QA*, and *C1QBP*) as well as the ubiquitin proteasome system (e.g., *UBE2/4* family, *UBA3B*, *UBA6*, *UCHL1*, *PSMA1*/*3*/*5*/*6*, *PSMC4*/*5*, and *PSMD4*/*6*/*7*/*13*), all of which are commonly dysregulated in neurodegenerative diseases, including AD (52,78–90). Importantly, some of the identified biological processes, especially the complement system, were also found in a SRT study recently published from brain tissue in a mouse model of AD (18).

To investigate the pathobiological implications of spatially-resolved Aβ-associated DEGs in neurodegeneration and dementia, we performed MAGMA and found that the peripheral microenvironments of Aβ plaques (next_Aβ spots) harbored significant genetic risk for AD. We further examined the expression of candidate genes associated with genetic risk for AD in our different Aβ microenvironments. Surprisingly, genes associated with increased risk for early- and late-onset AD did not exhibit differential expression in any of the 7 pathology-associated microenvironments in this late-stage AD tissue, with the exception of ADAM10 (91), a metalloprotease whose gene expression was reduced in the core region of Aβ-associated microenvironments (**Figure S14B**). This result could be due to their ubiquitous (e.g., *APOE*, *CLU*, *PLD3*, and *APP*) or sparse (e.g.,*TREM2* and *PSEN2*) gene expression patterns, which limit their detection of significant spatial variation in AD. Intriguingly, no enrichment of genes associated with frontotemporal dementia (FTD) was found in the core Aβ microenvironments. This observation appears to be consistent with the notion that FTD is devoid of cerebral amyloidosis (92,93), supporting the specificity of Aβ pathology in late-stage AD. Additionally, we observed a strong association between the peripheral compartment of Aβ microenvironment and Parkinson’s disease (PD), which may suggest a potentially shared genetic risk between AD and PD in this local brain microenvironment. Our subsequent clinical gene set enrichment analyses using genes identified in bulk RNA-seq (65) and single-nucleus RNA-seq (snRNA-seq) studies (66) confirmed the relevance of our dataset to AD while providing new technical insights into leveraging transcriptomic studies for gene set enrichment analyses. Notably, while genes identified from co-expression modules derived from bulk RNA-seq analysis using *SpeakEasy* (65,94), which is similar to *WGCNA* (95), did not reveal any enrichments, genes differentially expressed between cell type clusters derived from snRNA-seq analysis revealed spatial enrichment patterns of microglia in the peripheral Aβ-associated microenvironments. This example highlights the utility of leveraging integrated single cell/nucleus and spatial transcriptomic approaches to understand the complexity of human brain transcriptomes underlying AD. Indeed, the application of emerging computational analyses for large-scale SRT studies, including spot-level deconvolution (96–98) and cell-to-cell communication (99), could help to better characterize the multicellular composition and crosstalk within the Aβ plaque microenvironments in AD.

To further explore the DEGs associated with spatially-resolved Aβ microenvironments, we conducted RNAscope FISH-IF for a selected subset of genes identified with Visium-SPG. The cholesterol metabolism gene *IDI1* and complement gene *C3* were broadly enriched in the vicinity of Aβ-associated microenvironments, whereas *NINJ1* was significantly depleted in the immediate vicinity of Aβ plaques. *PPP3CA*, *UCHL1*, and *SST* were enriched within Aβ-associated microenvironments, but their enrichment gradually decreased with distance from the Aβ-associated microenvironment. These differential gene expression patterns and topography appeared to inform molecular and cellular events (100) occurring in the proximity of Aβ plaques during the late stages of AD. For example, cholesterol synthesis is altered in both neurons and glial cells in AD and is linked to Aβ accumulation (101–107), which may support our observation of the increased *IDI1* gene expression proximal to Aβ deposition. Elevated *C3* may signify enriched glial cell populations in the vicinity of Aβ and the inflammatory crosstalk between microglia and astrocytes (108,109). Indeed, preliminary findings show the presence of *C3^+^* microglia (TMEM119) and *C3*^+^ astrocytes (GFAP) near Aβ plaques (**Figure S16D**), but more work will be needed in the future to determine the extent to which this occurs at plaques across the cortex. In terms of the spatially depleted genes, *NINJ1* was initially identified as a nerve regeneration factor in the peripheral nervous system, but in the central nervous system, it is better known as a mediator of immune cell migration and non-programmed cell death such as necroptosis and pyroptosis by directing membrane rupture to release proinflammatory factors for elimination of pathogenic molecules (110). The decrease in *NINJ1* expression near Aβ can be interpreted in two ways: as a result of lytic cell death of immune cells or as a result of silenced pro-inflammatory reactions that promote the formation and/or spread of Aβ plaques (111,112). Further studies are required to understand how inflammatory factors like *C3* and *NINJ1* coordinate immune reactions in the vicinity of Aβ plaques. The calmodulin-dependent serine/threonine protein phosphatase *PPP3CA* (calcineurin), the deubiquitinating enzyme *UCHL1*, and the neuropeptide *SST* are all implicated in AD (85,113–115). Aβ plaques appear to be enriched near cells expressing these DEGs, but their density decreases with distance from these cells. Given that these three genes are dominantly expressed in neuronal populations in the human cortex (116), it will be informative in the future to investigate the mechanisms underlying the spatial relationship between the neuronal types and Aβ in terms of the plaque formation and transmission. Taken together, RNAscope-based analyses delineated local variations in gene expression around Aβ at cellular resolution. While the majority of tested genes showed good correspondence between Visium-SPG and RNAscope FISH-IF, there were examples where Visium-SPG patterns did not match spatial gene expression patterns with RNAscope FISH-IF. For example, in the Visium-SPG data, *C3* was highly enriched in the peripheral regions of Aβ plaque microenvironments, while with RNAscope FISH-IF, the peak was identified within the core microenvironments. Discrepancies between the two platforms are likely due to the difference in resolution, and hence using orthogonal techniques for further characterizing DEGs identified with Visium-SPG may provide important additional information.

While this study provides the groundwork for identifying changes in gene expression associated with spatially-resolved AD neuropathology, it has several important limitations that can be addressed in future research. First, our analysis focused on the gray matter. We noted a sparse distribution of Aβ (28) and/or pTau (117) in the white matter region near the deeper cortical layers. In particular, these Aβ-associated Visium spots in the white matter expressed several endothelial markers including *CLDN5* (Claudin 5) *and MCAM* (CD146). Cerebral amyloid plaques are rarely found in white matter (118), but our finding suggests that the Aβ deposits may pertain to vascular Aβ localized in capillaries and blood vessels (119–122). It would be interesting to investigate these subcortical Aβ-associated vascular microenvironments, given their putative contribution in white matter hyperintensity and cerebrovascular dysfunction in AD (123–128). Second, while our categorical pathology classification approach allowed us to identify DEGs associated with different pathological microenvironments, we acknowledge it may not address the full spatial heterogeneity of AD pathology, and in the future, more complex modeling approaches could be implemented. With more fine grained measurements one could identify DEGs whose spatial gene expression patterns are sensitive to the pathological gradients of Aβ and pTau. Third, limitations in the sample size precluded assessing differences across the spectrum of AD severity. Since the primary objective was to investigate microenvironments bearing AD-related neuropathology compared to those without, we used a within-subjects design and did not directly compare gene expression between AD donors and a neurotypical control. Nevertheless, our study provides a computational framework for designing future studies in larger cohorts that scale sample size to increase power to answer other biological questions. New innovations in next-generation sequencing, imaging, and computational platforms will also enable follow-up work that has the ability to profile the clinical heterogeneity of AD (129) across other related neurodegenerative diseases and dementias (130,131) as well as across cortical layers (132,133), subcortical structures (28,66,134,135), and developmental stages (136). Lastly, Visium-SPG is currently limited to surveying only a few proteins of interest, but new technologies that use DNA-barcoding approaches for protein detection (137–139) are now emerging. These enhanced platforms will improve the ability to use multiplexed antibodies to simultaneously track different variants of pathogenic Aβ and pTau (140,141). Emerging SRT platforms using complementary in situ sequencing approaches also have enhanced spatial resolution (142–144), which can finely resolve transcriptomic information at cellular and subcellular levels. Such features will be highly advantageous for detecting subtle differences in transcriptomic profiles of Aβ and pTau elements that are dispersed across neuronal and synaptic structures.

## Conclusion

In summary, we used Visium-SPG to identify local transcriptional signatures in spatially-defined Aβ-associated microenvironments in the inferior temporal cortex (ITC) of donors with late-stage AD. We applied RNAscope FISH-IF to further explore spatial gene expression patterns of Visium-identified genes associated with Aβ at cellular resolution. As a proof of concept study, our work lays the groundwork for deploying multi-omic approaches using both proteomics and transcriptomics to understand spatial heterogeneity in human brain diseases. While future studies are needed to produce mechanistic insights, the ongoing collective efforts to deepen single -omics with multi -omics data of different modalities, for example, with genomics, epigenomics, and metabolomics (145–147) are anticipated to contribute to information that sheds light on the complex biological processes and multifactorial nature of AD. To support ongoing effort in the neuroscience community to better understand molecular changes associated with AD, our work is distributed as a resource composed of freely accessible interactive websites for exploring the processed data in analysis-readyR/Bioconductor objects as well as raw data and reproducible code for the computational analyses.

## Methods

### Post-mortem human brain tissue samples

Brain donations in the Lieber Institute for Brain Development (LIBD) Human Brain Repository were collected from the County of Santa Clara Medical Examiner-Coroner Office in San Jose, CA and from the Western Michigan University Homer Stryker MD School of Medicine, Department of Pathology in Kalamazoo, MI, both under WCG IRB protocol #20111080. Four donors with Alzheimer’s disease (AD) ranged in age from 66 to 91 and were matched for age and demographics (European ancestry) with 1 neurotypical donor (**Table S1**). Clinical characterization, diagnoses, and macro- and microscopic neuropathological examinations were performed on all samples using a standardized paradigm. A 15-item series of questions assessing changes in memory, cognition, movement, caregiving, and daily functioning were obtained for all donors, as part of the LIBD Telephone Autopsy Questionnaire. Details of tissue acquisition, handling, processing, dissection, clinical characterization, diagnoses, neuropathological examinations, RNA extraction and quality control measures have been described previously (148,149). Alzheimer’s neuropathology was measured and rated with the Braak staging schema and CERAD scoring system (150) to evaluate tau neurofibrillary tangle burden and senile plaque burden, respectively. Scales range from Braak stage 0 to Braak stage VI, and CERAD is none-sparse-moderate-frequent.

### Visium-SPG immunofluorescence (IF) staining and image acquisition

Inferior temporal cortex (ITC, Brodmann area 20) from 3 AD subjects and 1 age-matched control was microdissected using a hand-held dental drill. Frozen samples were embedded in OCT (TissueTek Sakura, Cat# 4583) and cryosectioned at 10 micron thickness (Leica, CM3050s). Two to three non-adjacent tissue sections from the same ITC block were mounted on a Visium Spatial Gene Expression Slide (10x Genomics, Cat# 2000233) for each donor, with the exception of the neurotypical donor Br3874, for which two different brain blocks were used to address their different larminar structures. IF staining was performed according to the manufacturer’s instructions (10x Genomics, CG000312 Rev B) with no modifications. To maximize preservation of RNA integrity for the downstream Visium-SPG assay, the IF staining procedure was optimized to accommodate methanol fixation and short incubation periods. Briefly, tissue sections on a Visium slide were fixed for 30 minutes in pre-chilled methanol, treated with blocking buffer containing BSA (Miltenyi Biotec, Cat# 130-091-376) and the ribonuclease inhibitor, ribonucleoside-vanadyl complex RVC (New England Biolabs, Cat# S1402S), and incubated for 30 minutes at room temperature with primary antibodies against Aβ, pTau, GFAP, and MAP2 [mouse anti-β-amyloid antibody, clone 6E10 (Biolegend, Cat# 803001, 1:50), rabbit anti-Tau antibody pSer202/pThr205, clone AH36 (StressMarq, Cat# SMC601, 1:100), rat anti-glial fibrillary acidic protein (GFAP) antibody (Thermofisher, Cat# 13-0300, 1:70), and chicken anti-microtubule-associated protein 2 (MAP2) antibody (Abcam, Cat# ab92434, 1:70)]. Following a series of 5 washes, appropriate host species secondary antibodies conjugated to Alexa dyes were applied for 30 minutes at room temperature [donkey anti-mouse IgG conjugated to Alexa 488 (Thermofisher, Cat# A-21202, 1:200) donkey anti-rabbit IgG conjugated to Alexa 555 (Thermofisher, Cat# A-31572, 1:350), donkey anti-rat IgG conjugated to Alexa 594 (Thermofisher, Cat# A-21209, 1:400), and goat anti-chicken IgY conjugated to Alexa 633 (Thermofisher, Cat# A-21103, 1:300)]. DAPI (Thermofisher, Cat# D1306, 1:3000, 1.67 μg/ml for final concentration) was applied for nuclear counterstaining at the same time with the secondary antibodies. After a series of 5 washes and 20 immersions in 3x saline-sodium citrate (SSC) buffer diluted from 20x concentrate (Quality Biological, Cat# 351-003-131), the slide was coverslipped with 85% glycerol supplemented with RiboLock RNase inhibitors (Thermofisher, Cat# EO0382, 2 U/μl for a final concentration).

Slides were imaged on a Vectra Polaris (Akoya Biosciences) slide scanner with a MOTiF™ technology at 20x magnification using multispectral imaging and whole slide scanning. The exposure time per channel was as follows: 1.65-1.80 msec for DAPI; 380-400 msec for Opal 520; 43-65 msec for Opal 570; 150-178 msec for Opal 620; 615-643 msec for Opal 690; 100 msec for Autofluorescence (AF). There are several technical challenges to note regarding the staining, imaging, and analysis steps. First, chemical-based quenchers such as Trueblack (Biotium) were not used to overcome the lipofuscin autofluorescence in post-mortem human tissue due to their potential incompatibility with the Visium platform. Secondly, the dark green precipitates of the RVC ribonuclease inhibitor introduced particulate matter that added background fluorescence. This was an unknown issue in early releases of the Visium-SPG protocol, but it has been addressed in the recent manuals from 10x Genomics (10x Genomics, CG000312 Rev D) and resolved by heating the compound for improved suspension. Lastly, the current specifications of the whole-slide scanning system, though offering impressive speed and performance for multi-spectral imaging to accommodate the time-sensitive Visium-SPG workflow, are limited in its capacity for automatic signal intensity adjustment. This limitation restricted our ability to minimize local saturation of signals across several fields of view while imaging the entire tissue-wide neuropathology. Consequently, the pixel intensities were not considered as a reliable resource to estimate a pathological burden.

### Visium-SPG cDNA library generation and sequencing

Immediately after slide imaging, each tissue section was permeabilized and processed for cDNA synthesis, amplification, and library construction according to the Visium Spatial Gene Expression User Guide (10x Genomics, Cat# CG000239 Rev B). Prior to the Visium Spatial Transcriptomics assay, Visium Spatial Tissue Optimization experiments were performed to optimize permeabilization conditions using Visium Tissue Optimization Slides (10x Genomics, Cat# 3000394) with reference to Visium Spatial Tissue Optimization User Guide (10x Genomics, Cat# CG000238 Rev A). 12 minutes was determined to be the optimal permeabilization time for all samples (**Figure S1B**). The resulting whole transcriptome Visium libraries were sequenced on Illumina NovaSeq 6000 with S4 200 cycle kit using a loading concentration of 300 pM (read 1: 28 cycles, i7 index: 10 cycles, i5 index: 10, read 2: 90 cycles).

### IF image processing and segmentation for Aβ and pTau

We utilized multispectral imaging and spectral unmixing methods to address background staining from autofluorescent molecules such as lipofuscin, which is common in post-mortem human brain tissue (151,152), as well as from interference effects of undissolved precipitates in the RVC, a ribonuclease inhibitor used in the staining protocol. Specifically, we used a similar strategy to our previous study (153) that allowed us to identify and then mask lipofuscin and RVC-related particulate matter from all fluorescence channels (**Figure S2**). Thresholding background interference with this image processing step allowed us to better distinguish Aβ and pTau aggregates. Our workflow was as follows: first, QPTIFF image files were generated from a Vectra Polaris (Akoya Biosciences) slide scanner. An annotation was drawn to outline each entire slide in a given QPTIFF using Phenochart whole slide viewer (Akoya Biosciences). Spectral fingerprints, based on single positive samples, were generated for DAPI, Alexa 488, Alexa 555, Alexa 594, and Alexa 633, then selected to construct an unmixing algorithm. The unmixing algorithm was applied to the annotated region to decompose multi-spectral profiles into spectrally unmixed multi-channel TIFF tiles using inForm automated image analysis software (Akoya Biosciences). Autofluorescence, including lipofuscin, was segregated to an “AF” channel. Using the `InFormStitch()` function from *VistoSeg* (154), individual image tiles were stitched together into a full slide image using the coordinates of each tile that are saved in the filename. This stitched TIFF was split into multi-channel TIFF images containing an individual capture area using the `splitSlide_IF()` function from *VistoSeg*. GFAP and MAP2 staining data was initially planned as part of the study to investigate the local impact of AD on astrocytes and neuronal processes. However, due to the heavily branched and intermingled morphology of these cells, and the confounding particulate matter from RVC, it proved exceedingly challenging to accurately segment their morphological features. While the raw image data is available and shared to the scientific community, analysis of these image channels for quantification of GFAP and MAP2 signals has been excluded from the present study. Therefore, we focused on pathology-containing Aβ and pTau image channels for quantification. We used *dotdotdot* (153) to segment the channel containing DAPI. Due to the particulate matter background staining from RVC, we observed aberrant punctate staining common to multiple channels including Alexa 488 (Aβ), Alexa 555 (pTau), Alexa 594 (GFAP) and Alexa 633 (MAP2), in addition to the AF channel. To take a conservative approach, we masked the confounding fluorescent signals in Alexa 488 and Alexa 555 channels, using those from Alexa 633, Alexa 594, and AF channels (**Figure S2**). We then used individual thresholds based on the intensity histograms of Aβ and pTau to segment regions of interest (ROIs) from their respective fluorescent channels. For example, in tissue samples from Br3880, ROIs smaller than 1500 pixels were dropped from the Aβ channel due to the inability to distinguish small diffuse Aβ plaques from background lipofuscin and particulate RVC. Then we applied a shape filter to the size-thresholded Aβ ROIs, to retain ROIs mostly circular in shape. We then calculated both the number of ROIs per Visium spot (NABeta and NPTau, the N prefix is for number) and the proportion of the Visium spot covered by segmented signal (PABeta and PPTau, the P prefix is for proportion). In the neurotypical control samples, the same image thresholding approach was applied to address nonspecific background staining, and the final segmentations from samples from a neurotypical control were used to set thresholds in the AD samples.

### Visium raw data processing and quality control (QC)

FASTQ and image data were pre-processed with the 10x *SpaceRanger* pipeline (version 1.3.0 for Visium-SPG) (155). Outputs of *SpaceRanger*, including the spotcounts matrix, were stored in a *SpatialExperiment* v1.5.3 (156) object (Bioconductor version 3.13) for downstream analysis. Data were filtered to remove genes that were not detected and spots with zero counts. We used *scran* v1.23.1 (Bioconductor version 3.13) (157) to calculate QC metrics, including mitochondrial expression, low library size, and low gene count. Many of the low-quality spots resided in the white matter and/or at the outermost edges near the leptomeninges (**Figure S9A**). Due to their potential pathobiological significance in cortical laminar organization and AD (158–160), we retained these spots in the downstream analyses. However, we excluded 152 edge spots with low library sizes, which were located on the edges of tissue sections. We also manually eliminated 20 Aβ-positive spots that were falsely annotated due to glare artifacts in the Aβ channel (**Figure S9B**). This preprocessing step for spot-level QC filtered out 172 spots, yielding a total of 38,115 spots across the 10 samples for downstream analyses. We performed dimensionality reduction using the *scater* v1.23.1 package (161) to calculate the top 50 PCs followed by using the *Harmony* package v0.1.0 (38) to perform batch-correction across Visium capture areas.

### Unsupervised clustering by *BayesSpace*

To study AD-related neuropathology from gray matter, we applied the spatially-aware clustering method *BayesSpace* v1.5.1 (39) to identify anatomical regions in the Visium data that best corresponded to gray matter spanning cortical layers. *BayesSpace* is a Bayesian statistical model that uses the spatial arrangement of Visium spots to borrow information from each spot’s neighborhood in order to assign clusters, thereby producing smoother spatial clusters.

To use *BayesSpace* on all samples at once, we arranged all samples into the same plane as recommended for joint clustering analysis https://edward130603.github.io/BayesSpace/articles/joint_clustering.html with a row offset of 100 per sample. We used `spatialCluster()` with default parameters for `q` = 2 up to 28 applied on the 50 batch-corrected dimensions from *Harmony* (38). Note that *BayesSpace* is not guaranteed to result in *k* equal to *q* as it can merge smaller initial clusters (`init.method = "mclust"` by default) into a single spatial cluster. For example, `spatialCluster(q = 28)` resulted in 24 clusters.

### Spatial registration of *BayesSpace* clusters to identify a data-driven gray matter

On the batch corrected *SpatialExperiment* object, spots were pseudo-bulked by their *BayesSpace* cluster for q = 2 through 28 and sample id (Visium capture area), using the `aggregateAcrossCells()` function from *scuttle* v1.5.0 (161). Read counts were normalized to log counts using the `calcNormFactors()` function from *edgeR* v3.37.0 (162). `filterByExpr()` from *edgeR* with default parameters was then used to filter genes that were lowly expressed.

Adjusting for age, sex, and diagnosis, differential gene expression across clusters was then performed using the enrichment model as described previously (24). From *spatialLIBD* v1.7.13 (40), `layer_stat_cor()` was used to compute the correlation between the enrichment results of each BayesSpace cluster for each k and the enrichment results available in Supplementary Table 4 from Maynard, Collado-Torres et al (24). The `fetch_data()` function from *spatialLIBD* (40) was used to retrieve this reference dataset. These correlation results were visualized as a heatmap using the `layer_stat_cor_plot()` function.

### Spot-level identification and annotation of heavy-burden AD-related neuropathology

To identify Visium spots with a heavy pathological burden of Aβ and/or pTau for downstream analyses, the pathological burden was measured and thresholded at the spot level, using Aβ and pTau signals, which were quantified by the number (N) of their ROI objects and proportion (P) of pixels covering the ROIs within a spot, respectively. The 99.9th percentile values of NAbeta and PAbeta in samples V10A27004_A1_Br3874 and V10T31036_A1_Br3874, which are both control tissue samples, were identified as Aβ thresholds. As pTau thresholds, the maximum values of NpTau and PpTau in the control samples were identified. Thus, spots with a higher number of Aβ ROI objects (NAbeta) than 1 or a greater proportion of Aβ pixel spot coverage (PAbeta) than 0.108 were labeled to have significant Aβ pathology. Spots with a higher number of pTau ROI objects (NpTau) than 8 or a greater proportion of pTau pixel spot coverage (PpTau) than 0.0143 were identified as having significant pTau pathology. The `.find_neighbors()` internal function in *BayesSpace* v1.6.0 was used to identify neighboring spots to each pathology spot and were further characterized based on whether they were only adjacent to an 1) Aβ spot, 2) pTau spot or 3) ‘both’ spot.

### Differential gene expression analysis and modeling on gray-matter spots

Spots from *BayesSpace* `q = 2` corresponding to cluster 2, which are the white matter spots, were first dropped as well as those from the control donor Br3874. This resulted in 3,874 spots for none; 1,072 for Aβ; 2,284 for next_Aβ; 8,810 for pTau; 3,217 for next_pTau; 727 for both; and 1,102 for next_both. Then `registration_pseudobulk(min_ncells = 15)` from *spatialLIBD* v1.9.18 (40) was used to pseudo-bulk cells of the same sample ID and pathological category resulting in 41 out of the possible 49 (7 samples x 7 pathological categories) pseudo-bulked samples passing the `min_ncells = 15` filter. Next, `filterByExprs()`, `cpm()`, and `calculateNormFactors()` from *edgeR* (162,163) were used to filter out lowly expressed genes and log-normalize counts resulting in 8,443 genes passing these filters. We then identified differentially expressed genes using three models, as in our previous work (24), using new *spatialLIBD* (40) functions with the `registration_*()` prefix. Given the limited variability across donors, we did not adjust for any covariates. For example, sex is perfectly confounded with sample Br3880 as the other two AD cases are females. First, the ANOVA model used `registration_stats_anova()` from *spatialLIBD* and powered by *limma* (164) to test for differences in mean expression across the 7 pathological categories and found no DEGs (FDR<0.05). Second, the enrichment model used `registration_stats_enrichment()` from *spatialLIBD* to test for differences in expression between one pathological categories versus the remaining six and identified unique 1,840 genes that are DEGs (FDR<0.05) for at least one pathological category. These genes can be either enriched (t>0) or depleted (t<0). See **Table S2** for the Aβ results and **Table S3** for the next_Aβ results. Third, the pairwise model used `registration_stats_pairwise()` from *spatialLIBD* to test for DEGs between each pair of pathological categories (21 pairs total) resulting in 98 unique genes having significant differences (FDR<0.05) in at least one pair. See https://github.com/LieberInstitute/Visium_SPG_AD/blob/master/code/05_deploy_app_wholegenome/Visium_SPG_AD_wholegenome_model_results_FDR5perc.csv for the full results. *EnhancedVolcano* v1.16.0 (165) was used to make the volcano plots for the enrichment model results comparing the log2 fold change against the *p*-values, with a FDR<0.1 cutoff line.

### Differential gene expression analysis and modeling on gray-matter spots for the Aβ microenvironment

Similar to the previous analysis, we re-categorized the Aβ and next_Aβ spots as “Aβ_env”. We then performed the same type of analysis using functions from *spatialLIBD* (40). With this analysis only one gene was significant (FDR<0.5) for the *anova* model: *TMEM163.* There were 287 DEGs for the *enrichment* model and 15 unique DEGs for the *pairwise* model (FDR<0.5). See **Table S4** for the Aβ_env results or https://github.com/LieberInstitute/Visium_SPG_AD/blob/master/code/18_deploy_app_wholegenome_Abeta_microenv/Visium_SPG_ <u>AD_wholegenome_model_results_FDR5perc.csv</u> for the full results. 275 out of the 287 *enrichment* DEGs corresponded to the Aβ_env, with 265 of them depleted (t<0) and 10 of them enriched (t>0). We used `compareCluster(pvalueCutoff = 0.3, qvalueCutoff = 0.3)` with `enrichGO()` and `enrichKEGG()` from *clusterProfiler* v4.6.2 (166) with the Aβ_env enrichment DEGs. While GO terms and KEGG pathways were not significantly enriched (FDR<0.5), enriched (t>0) Aβ_env genes *CYP1B1*, *TNFRSF1A, ZNF385A*, and *RIOK3* were most frequently associated with GO terms.

### Statistics and reproducibility

Statistical tests were performed with R version 4.2 with Bioconductor versions 3.13, 3.14, 3.15, and 3.16.0. In addition to software versions listed in the Methods, R session information was recorded with `sessioninfo::session_info()` for many specific analyses on log files or scripts themselves and is available on GitHub and Zenodo (Data and materials availability).

### MAGMA

Version (v1.10) of Multi-marker Analysis of GenoMic Annotation (MAGMA) (61) was used to measure genetic risk association of spots containing Aβ plaques and neighbored Aβ plaques, representing the core (Aβ) and periphery (next_Aβ) of Aβ-associated microenvironments, with Alzheimer’s disease (AD: Jansen et al. (62)); Frontal Temporal Disorder (Ferrari et al. (63)); and Parkinson’s Disease (PD: Nalls et al. (64)). Genes identified as either significantly enriched or depleted based on the Benjamini & Hochberg procedure with a FDR<0.1 (167) were used to create the marker gene sets for each pathology type. We used MAGMA to first annotate SNPs to genes, with window sizes of 10kb to +35kb for each gene. The 1000 Genomes EUR reference panel was used for this step (168). Next, gene-level analyses were performed using the summary statistics provided from the above-mentioned GWAS studies. In the MAGMA software, we used the `snp-wise = mean` model to test whether any of the genes were enriched for genetic risk for each of the GWAS diseases. Finally, competitive gene set analysis was performed with the marker gene sets for Aβ and n_Aβ categories to test whether they are related to the GWAS diseases (Figure S13A).

### Gene set enrichment analysis

Grubman et al (66) reported a list of differentially expressed genes that overlaps between their findings and those of Mathys et al (136) for each cell type from AD samples. For genes with a reported FDR<0.1, we performed an enrichment analysis using Aβ and n_Aβ categories with *spatialLIBD* v1.11.12 gene_set_enrichment()(40) (Figure S13C). We performed the same enrichment analysis on differentially expressed genes for AD on bulk RNA-seq reported by Mostafavi et al (65) (Figure S13B).

### RNAscope smFISH combined with immunofluorescence (FISH-IF) for RNA-protein co-detection

An independent brain donor diagnosed with AD was added to the cohort for this experiment (**Figure 3A** and **Table S1**), and 1 tissue slice from ITC for each of the 4 AD donors was prepared and assayed with the same monoclonal Aβ antibody (6E10) and commercially available RNAscope probes for the genes of interests. In detail, frozen tissue blocks from each of the 4 AD subjects were sectioned at 10 micron thickness (Leica, CM3050s) and stored at −80°C. In Situ Hybridization (ISH) assays were performed with RNAscope technology utilizing the RNAscope Fluorescent Multiplex Kit v.2, 4-plex Ancillary Kit, and RNA-Protein co-detection ancillary kit (Advanced Cell Diagnostics ACD, Cat# 323100, 323120, and 323180) in reference to the manufacturer’s instructions: Multiplex Fluorescent Reagent Kit v2 User Manual (ACD, 323100-USM) and RNAscope^®^ Multiplex Fluorescent v2 Assay combined with Immunofluorescence - Integrated Co-Detection Workflow (ACD, MK 51-150 Rev A, Appendix C). Tissue sections were fixed with pre-chilled 10% neutral buffered formalin solution (Sigma-Aldrich, Cat# HT501128) for 15 minutes at 4°C, dehydrated in increasing concentrations of ethanol (50%, 70%, 100%, and 100%), and pretreated with hydrogen peroxide for 10 minutes at room temperature (RT). Sections were incubated with the mouse anti-β-amyloid antibody, clone 6E10 (Biolegend, Cat# 803001, 1:50) overnight at 4°C. After primary antibody incubation, sections were fixed in 10% neutral buffered formalin solution for 30 minutes at RT and treated with protease IV for 30 minutes at RT. The following RNAscope probes were used for in ISH reaction: *PPP3CA*, *UCHL1*, *SST*, *C3*, *IDI1*, and *NINJ1 (*ACD, Cat# 477311, 594281-C2, 310591-C3, 485641, 430701-C2, and 540051-C3, respectively, 1x dilution for *PPP3CA* and *C3*, and 1:50 (50x) for *UCHL1*, *SST*, *IDI1*, and *NINJ1*). The 6 probes were grouped into 2 combinations and multiplexed to co-label 1) *PPP3CA*, *UCHL1*, and *SST* and 2) *C3*, *IDI1*, and *NINJ1* within each tissue sample. After probe labeling, sections were subjected to amplification steps (AMP1–3) and conjugation to fluorescent Opal dyes 570, 620, and 690 (Akoya Biosciences, Cat# FP1488001KT, FP1495001KT, and FP1497001KT, respectively, 1:600-1:2500). At the end of RNAscope procedures, sections were treated with donkey anti-mouse IgG conjugated to Alexa 488 (Thermofisher, Cat# A-21202, 1:400) for 40 minutes at RT and stained with the DAPI solution provided in the RNAscope multiplex kit for counter nuclear staining. The slides were then coverslipped with Fluoromount G (Southern Biotechnology, Cat# 0100-01) and imaged on Vectra Polaris with a MOTiF™ scanning mode. The resulting QPTIFF images were annotated, spectrally unmixed, and fused to recreate an entire slide image using Phenochart (Akoya Biosciences), inForm (Akoya Biosciences), and HALO^®^ image analysis platform (Indica labs), respectively.

### RNAscope quantification and spatial proximity analysis using HALO

Prior to segmentation, each tissue section was manually annotated to define a gray matter region based on enrichment of Aβ plaques and to exclude problematic areas such as smeared signals and tissue tears found on the edges. The FISH-IF module and proximity analysis module (version 2.1.5) in HALO were utilized to assess the gene expression changes of genes of interest (GOIs) in the proximity of nearby Aβ plaques with reference to the manufacturer’s guidelines: HALO 3.3 FISH-IF Step-by-Step guide (Indica labs, Version 2.1.4 July 2021) and Digital Quantitative RNAscope Image Analysis Guide (Indica labs). For segmentation of extracellular Aβ plaques, the FISH-IF module was optimized to treat Aβ as a nuclear and cellular marker. This process required a paralleled FISH-IF segmentation to analyze cells expressing and not expressing GOIs, independent of the Aβ segmentation (**Figure S15C**). For a given tissue section, therefore, two FISH-IF datasets were generated, one for Aβ and another for GOIs, which we combined to map the segmented objects back to the identical tissue sections (**Figure S15A**). Segmentation was optimized for each object, including Aβ and RNAscope signals for *PPP3CA, UCHL1, SST, C3, IDI1,* and *NINJ1*, using user-defined size and intensity thresholds. A thorough visual inspection was performed to ensure that the segmentation outputs matched the input raw images in terms of object size and shape. The same thresholds were applied to a given GOI across all tissue samples, regardless of the donors. The size thresholds for nucleus and cytoplasm as well as those for Aβ plaques remained constant. Only the copy intensity parameter was adjusted by individual tissue sections to address staining variations between samples, using their average cell intensity of RNAscope signals.

The two datasets of Aβ and GOIs from each tissue section were subjected to downstream proximity analysis to measure the distance between the segmented Aβ and cells expressing and not expressing GOIs, namely GOI- and non-expressors. Based on their proximity to the nearby Aβ, the GOI- and non-expressors were classified into the following 7 intervals: 1) 0-21.25µm, 2) 21.25-42.5µm, 3) 42.5-63.75µm, 4) 63.75-85µm, 5) 85-106.25µm, 6) 106.25-127.5µm, and 7) >127.5µm. The spatially categorized cells from the proximity analysis were then aligned with the complementary FISH-IF dataset for the GOI expressors and non-expressors and assigned their corresponding RNAscope puncta counts, using their unique cell identifier (ID) numbers. The puncta counts were then summed up to compute the average single cell gene expression in each interval. Normalization was implemented with the total number of GOI expressors per interval. For the spatially down-regulated and/or depleted genes identified from the Visium assays, we normalized their gene expression by the total number of cells existing in each sub-region. We summed the GOI expressors and non-expressors and used the value as a denominator to account for the reduction in gene expression driven by not only downregulation of GOIs within the same cell, but also the complete absence of GOI expressors in a defined sub-area.

The latter was more pronounced with genes whose expression was sparsely distributed, such as *SST*. This normalization approach aided in the interpretation of the spatial gradients of several GOIs with statistical significance (**Figure S16A-B)**. A log2 (X+1) transformation was implemented, where X is the raw puncta count, to account for the skewed cellular distribution as well as the non-expressors with zero puncta (**Figure S16C**). Quantification plots and statistical testings for the non-parametric Kruskal-Wallis test were made with Prism (version 9).

### Data and materials availability

The source data are also publicly available from the Globus endpoint ‘jhpce#Visium_SPG_AD’ that is also listed at http://research.libd.org/globus. The raw data provided through Globus includes all the FASTQ files and raw image files. The FISH-IF and proximity data, as well as the segmentation settings, were exported as text files and are available on the raw data directory named AD_RsFISHIF_RawData. Processed data are publicly available from the Bioconductor package *spatialLIBD* version 1.11.12 (40) through the `fetch_data()` function. All source code is publicly available through GitHub and permanently archived through Zenodo at https://github.com/LieberInstitute/Visium_SPG_AD (169). Interactive websites are powered by *spatialLIBD* (40), *iSEE* (170), and *samui* (171).

## Supporting information

Supplementary Figures at 300 ppi

## Acknowledgements

We acknowledge Louise A. Huuki-Myers, Boyi Guo, Matthew N. Tran, Svitlana Bach, and Ryan A. Miller for their insightful assistance with statistical testing, MAGMA analyses, data interpretation, and data transfers respectively. They were all employed by the Lieber Institute for Brain Development (LIBD) except for Boyi Guo from the Johns Hopkins Bloomberg School of Public Health, Department of Biostatistics. We acknowledge Elizabeth Engle and the Johns Hopkins Tumor Microenvironment Lab core facility for assistance with the Vectra Polaris slide scanner. We would like to thank the Joint High Performance Computing Exchange (JHPCE) for providing computing resources for these analyses. We thank Amy Deep-Soboslay and James Tooke from LIBD for curation of brain samples and assistance with coordinating dissections. We also thank the office of the County of Santa Clara Medical Examiner-Coroner Office in San Jose, CA and the Western Michigan University Homer Stryker MD School of Medicine, Department of Pathology in Kalamazoo for making post-mortem human tissue donations possible to advance these studies. We are indebted to the generosity of families of brain donors for supporting this research. Finally, we thank the families of Connie and Stephen Lieber and Milton and Tamar Maltz for their generous support of this work. Schematic illustrations were generated using Biorender.

## Author contributions

Conceptualization: KM, KRM

Methodology: KM, KRM, LCT, SHK, MT, SP

Software: LCT, MT, SP, HRD, NJE

Validation: SHK, JSL

Formal analysis: SHK, MT, LCT, SP

Investigation: SHK, MT, LCT, SP, JSL

Resources: TMH, JEK, RAB, SRW, MM

Data curation: SHK, LCT, SP, MT

Writing-original draft: SHK, MT, SP, LCT

Writing-review and editing: KM, KRM, SCP, SCH, LCT

Visualization: SHK, LCT, SP

Supervision: KM, KRM, LCT, SCP, SCH

Project administration: KM, KRM Funding acquisition: KM, KRM

## Tables

**Supplementary Table S1. Sample donor demographics.**

This table summarizes the donor demographic information, clinical diagnosis, tissue sample details, and study design for the spatially-resolved transcriptomics study based on Visium-SPG and RNAscope FISH-IF assays. The information provided includes the demographic and clinical details of the donors selected for the study, the types of assays performed, the number of Visium tissue replicates, and tissue sample identifiers, including slide serial numbers and Visium capture areas.

**Supplementary Table S2. DEG statistics for the enrichment model for the Aβ category across the 7 pathological categories.**

This supplementary table presents the results of differential expression (DE) testing for Aβ spots (i.e., the ‘Aβ’ category), based on the enrichment model, using the pseudo-bulked Visium-SPG data. The table includes columns for Ensembl gene ID, gene name (symbol), log2 fold change (logFC), enrichment model *t*-statistic, two sided *p*-value, and FDR-adjusted *p*-value. This table can also be re-downloaded from https://libd.shinyapps.io/Visium_SPG_AD_wholegenome/ under the tabs labeled “layer-level data” and “model box plots”.

**Supplementary Table S3. DEG statistics for the enrichment model for the next_Aβ category across the 7 pathological categories.**

This supplementary table presents the results of differential expression (DE) testing for the Aβ-adjacent spots (i.e., the ‘next_Aβ’/‘n_Aβ’ category), based on the enrichment model, using the pseudo-bulked Visium-SPG data. The table includes columns for Ensembl gene ID, gene name (symbol), log2 fold change (logFC), enrichment model *t*-statistic, two sided *p*-value, and FDR-adjusted *p*-value. This table can also be re-downloaded from https://libd.shinyapps.io/Visium_SPG_AD_wholegenome/ under the tabs labeled “layer-level data” and “model box plots”.

**Supplementary Table S4. DEG statistics for the enrichment model for the Aβ_env category across the 6 pathological categories, when Aβ and next_Aβ were combined into Aβ_env.**

This supplementary table presents the results of differential expression (DE) testing for the combined Aβ and Aβ-adjacent spots (i.e., the ‘Aβ microenvironment’/‘Aβ_env’ category), based on the enrichment model, using the pseudo-bulked Visium-SPG data. The table includes columns for Ensembl gene ID, gene name (symbol), log2 fold change (logFC), enrichment model *t*-statistic, two sided *p*-value, and FDR-adjusted *p*-value. This table can also be re-downloaded from https://libd.shinyapps.io/Visium_SPG_AD_wholegenome_Abeta_microenv/ under the tabs labeled “layer-level data” and “model box plots”.

## Supplemental information

**Figure S1.**
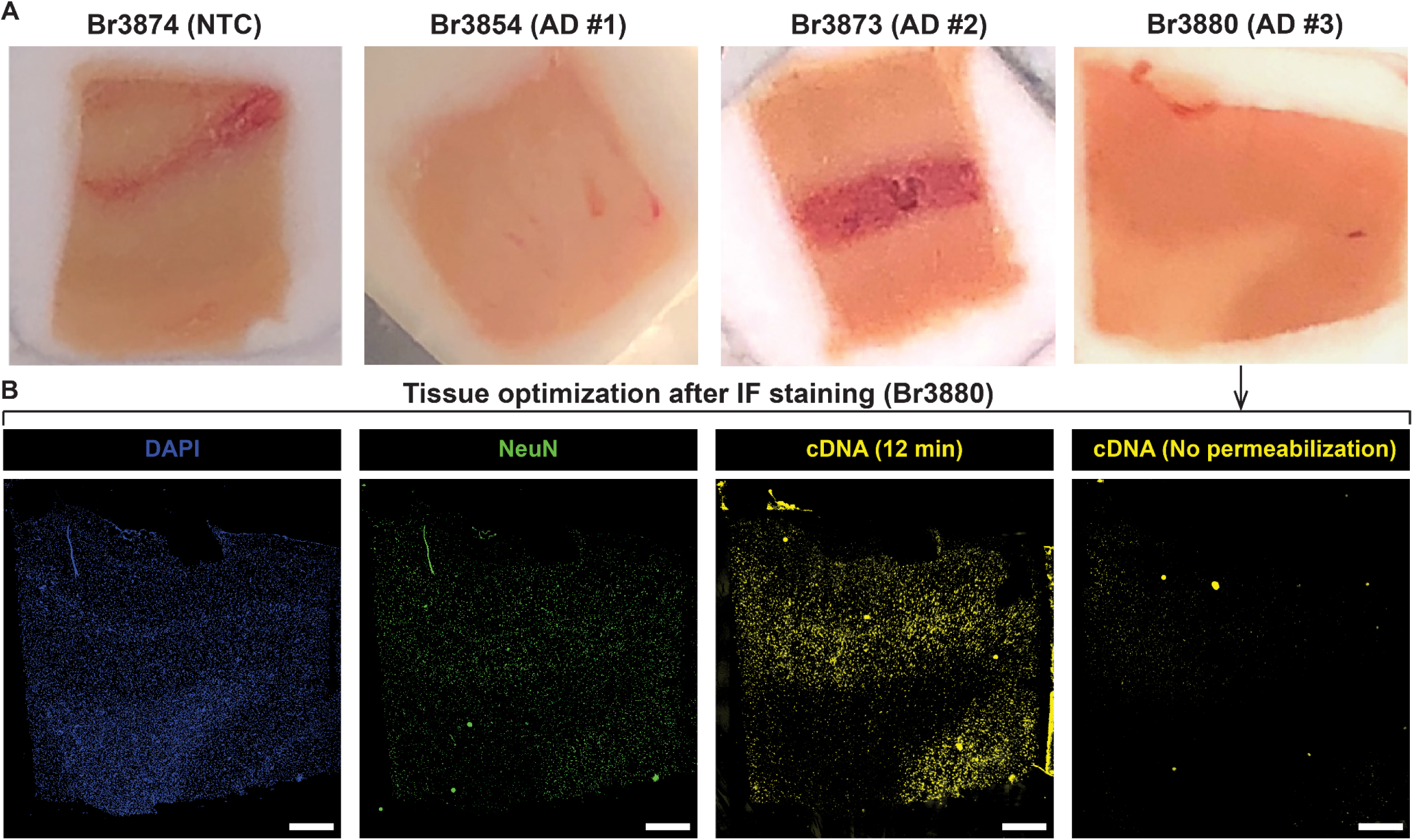
Tissue permeabilization optimization for ITC blocks used for Visium-SPG experiments. (**A**) Representative images of inferior temporal cortex (ITC) tissue blocks obtained from 1 neurotypical control (NTC) and 3 donors diagnosed with AD that were used for Visium-SPG experiments. (**B**) Representative image from Br3880 of tissue permeabilization optimization experiment. Tissue optimization experiments were conducted to determine the optimal permeabilization time for sufficient cDNA synthesis following immunofluorescence staining. Cryosections from the tissue blocks were immunostained for NeuN and counterstained with DAPI to label the nucleus. After whole-slide imaging, the same cryosections were permeabilized for different time points and subjected to cDNA synthesis with fluorescently labeled nucleotides resulting in a fluorescent cDNA footprint (yellow). The optimal permeabilization time was determined to be 12 minutes, a timepoint at which robust cDNA synthesis was consistently observed across all samples and donors. A non-permeabilized tissue section lacking successful cDNA synthesis shows the baseline fluorescence level. Scale bar, 1mm.

**Figure S2.**
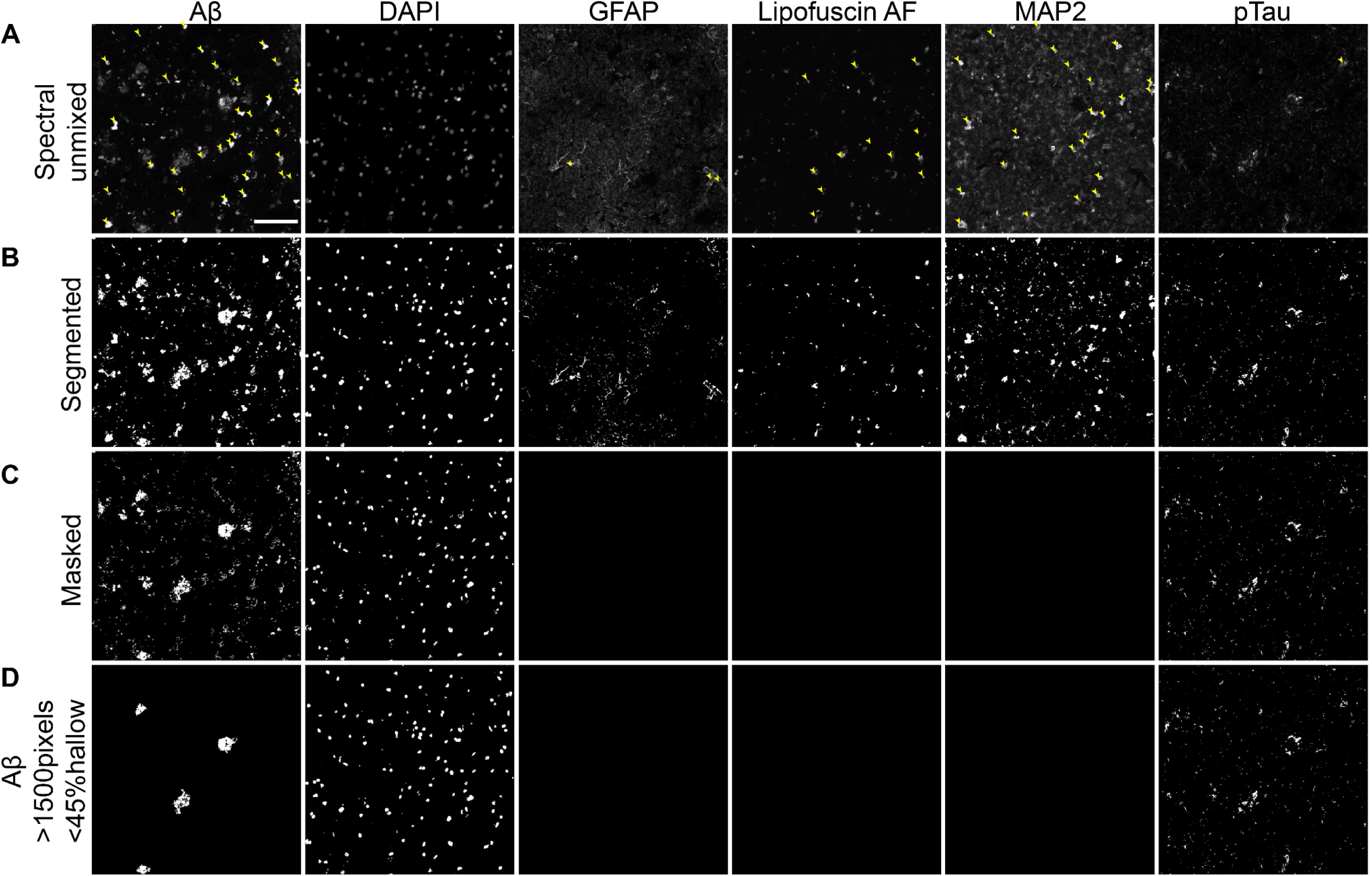
Image processing workflow to threshold background fluorescence and staining artifacts. High-magnification grayscale images of multichannel IF staining of the gray matter from AD donor Br3880 illustrating the sequence of image processing steps applied to all 6 fluorescence channels. (**A**) After spectral unmixing, residual lipofuscin and particulate matter of ribonucleoside vanadyl complex (RVC), a ribonuclease inhibitor, were observed as background noise and confounding staining artifacts (yellow arrowheads) across multiple channels of the processed IF image including Aβ, GFAP, lipofuscin autofluorescence (AF), MAP2, and pTau. (**B**) All channels were segmented by intensity-based thresholding for each individual channel. (**C**) The segmented objects in the GFAP, AF, and MAP2 channels were used to mask and remove noise and artifacts from all channels. (**D**) The Aβ channel was further thresholded to isolate Aβ plaques using their distinct morphology (4th row, Aβ) based on size (>1500 pixels) and shape (<50% hollow). Detailed descriptions of staining artifacts and image processing workflow were provided in Methods. Scale bar, 100µm.

**Figure S3.**
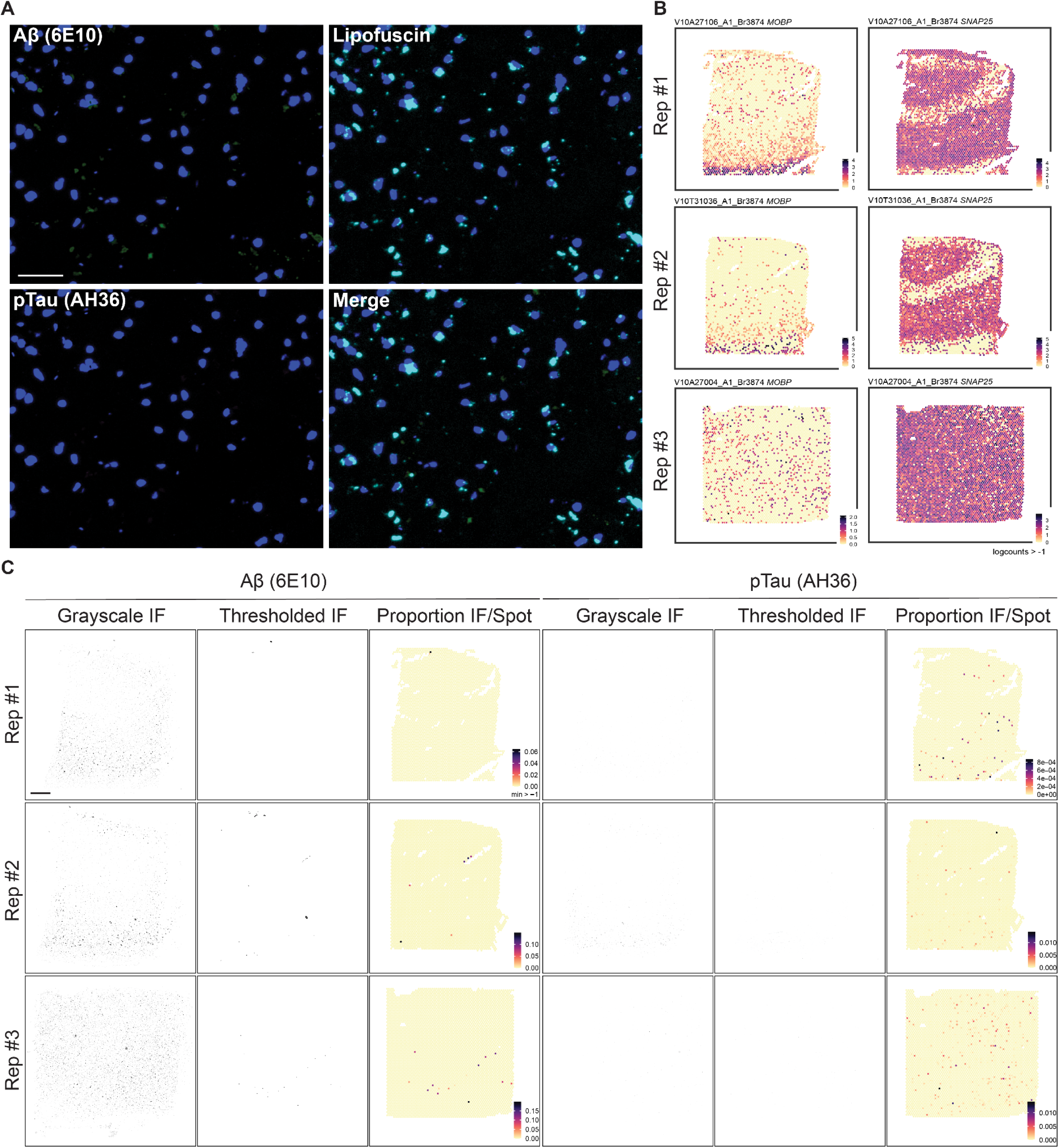
Validation of immunostaining (IF) and image processing workflow using neurotypical control samples. (**A**) Representative high magnification images of Aβ (6E10), pTau (AH36), and lipofuscin fluorescence in control tissue sections showing little to no AD-related neuropathology in the control donor. Scale bar, 50µm. (**B**) Spotplots of *SNAP25* and *MOBP* across 3 replicate tissue sections from neurotypical control Br3874 showing spatial orientation of gray and white matter, respectively. Replicate 3 (Rep3) was obtained from a different tissue block from the same donor, showing a different tissue-wide morphology. The unclear separation between gray and white matter was noted in this replicate but still included in the downstream analyses for *Harmony*-based batch correction as in **Figure S10**. (**C**) IF image thresholding of Aβ and pTau pathology in control tissue sections and integration with the complementary SRT data. Raw grayscale IF images of Aβ and pTau signals were segmented and thresholded to integrate with complementary spatially resolved spot-level transcriptomes. Thresholded IF images show segmented and thresholded IF signals by size and shape and thus removal of the autofluorescent signals and noise due to lipofuscin and particulate matter of ribonucleoside vanadyl complex (RVC), a ribonuclease inhibitor used in the staining protocol. 6E10 and AH36 refer to clonal names of the monoclonal antibodies raised against Aβ and pTau, respectively. As expected for this control donor, we observed little to no pathology signal on the thresholded IF. Color scale indicates the proportion of Aβ and pTau pixels per Visium spot. Scale bar, 1mm.

**Figure S4.**
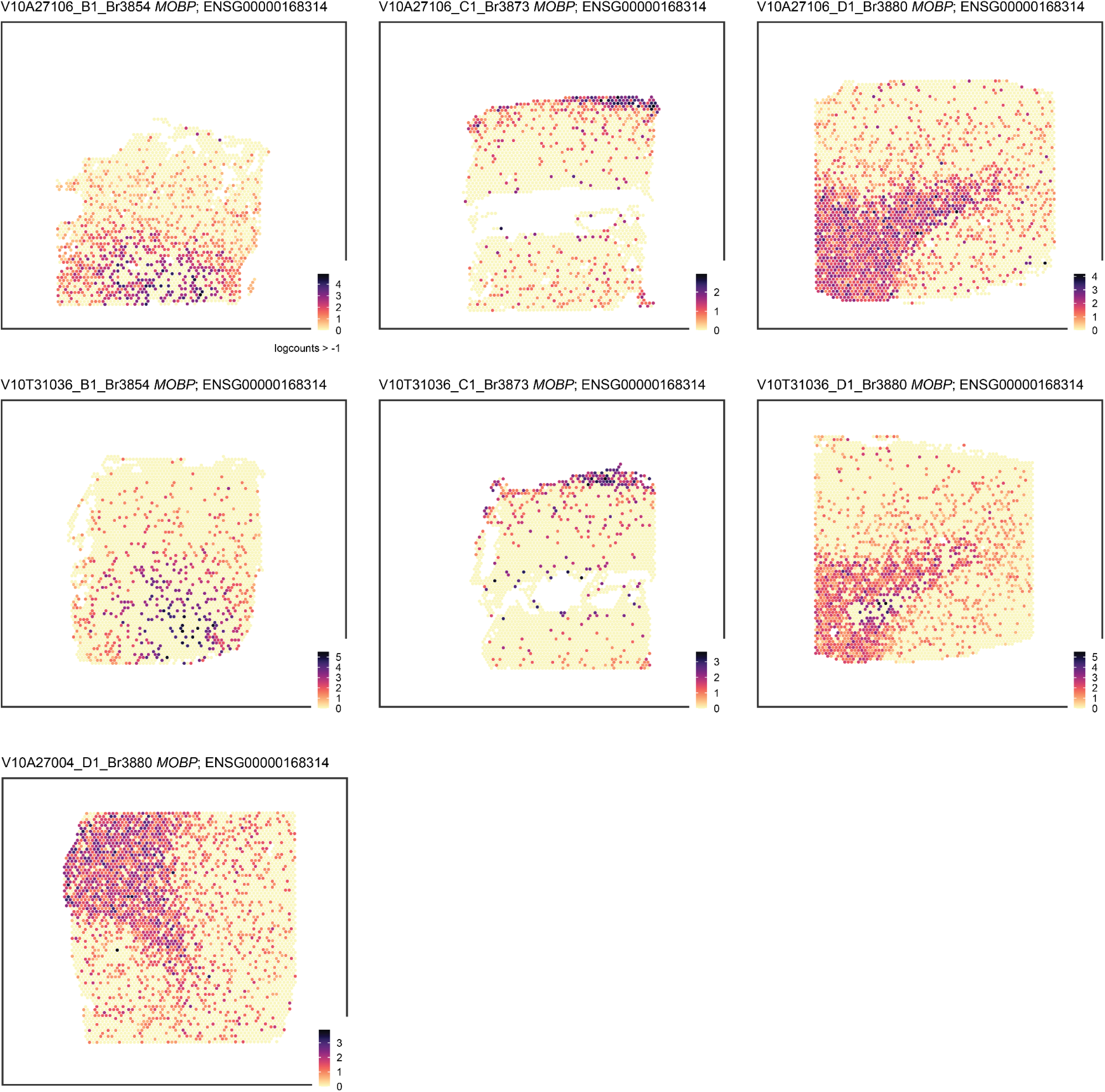
Spotplots of *MOBP* across 7 AD tissue sections from 3 AD donors. Spot-level gene expression of *MOBP* (myelin-associated oligodendrocyte basic protein) confirms the location of the white matter in each tissue section. Color scale indicates spot-level gene expression in logcounts.

**Figure S5.**
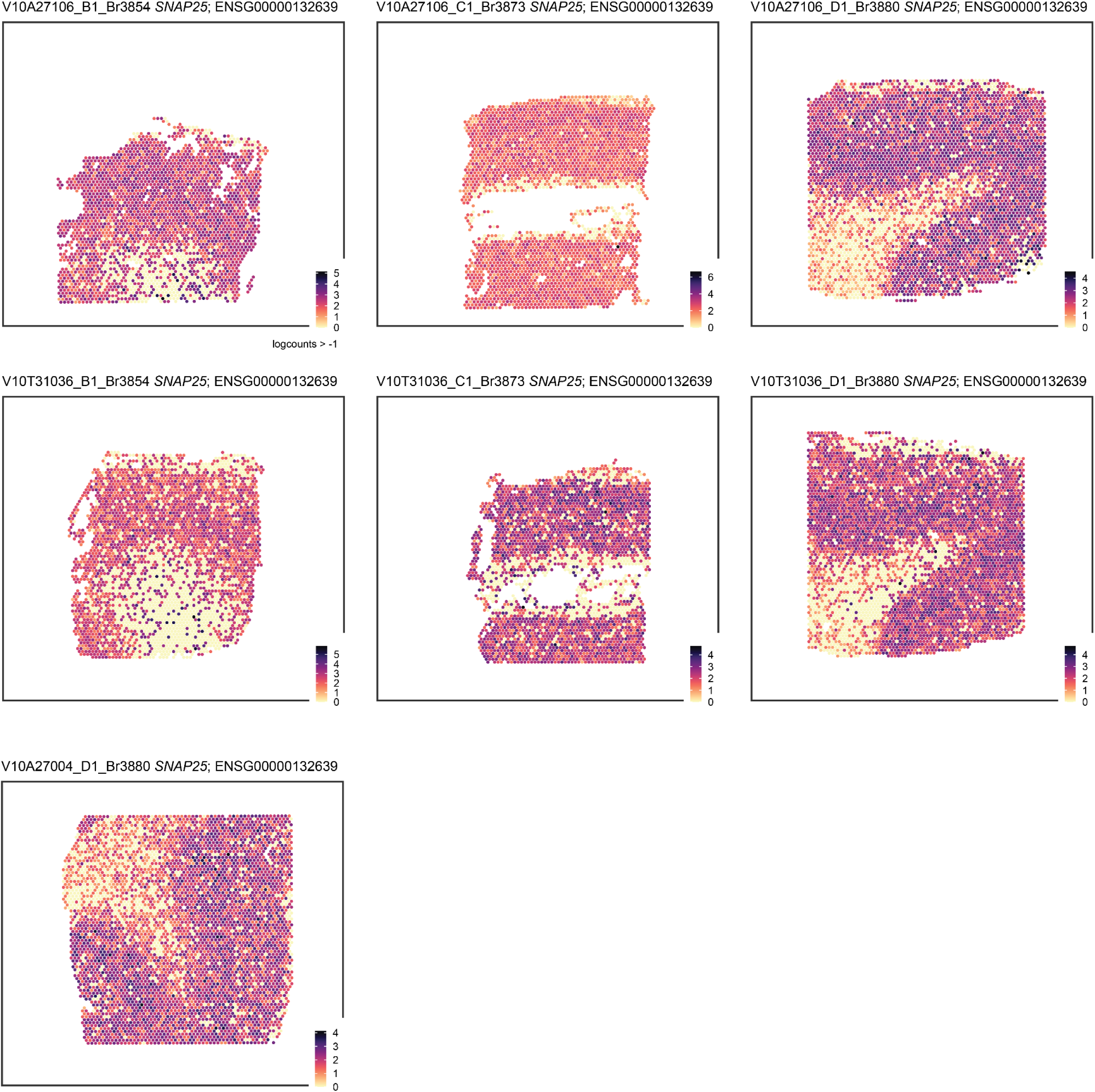
Spotplots of *SNAP25* across 7 AD tissue sections from 3 AD donors. Spot-level gene expression of the neuronally-enriched gene *SNAP25* (synaptosomal-associated protein 25) confirms the location of the gray matter in each tissue section. Color scale indicates spot-level gene expression in logcounts.

**Figure S6.**
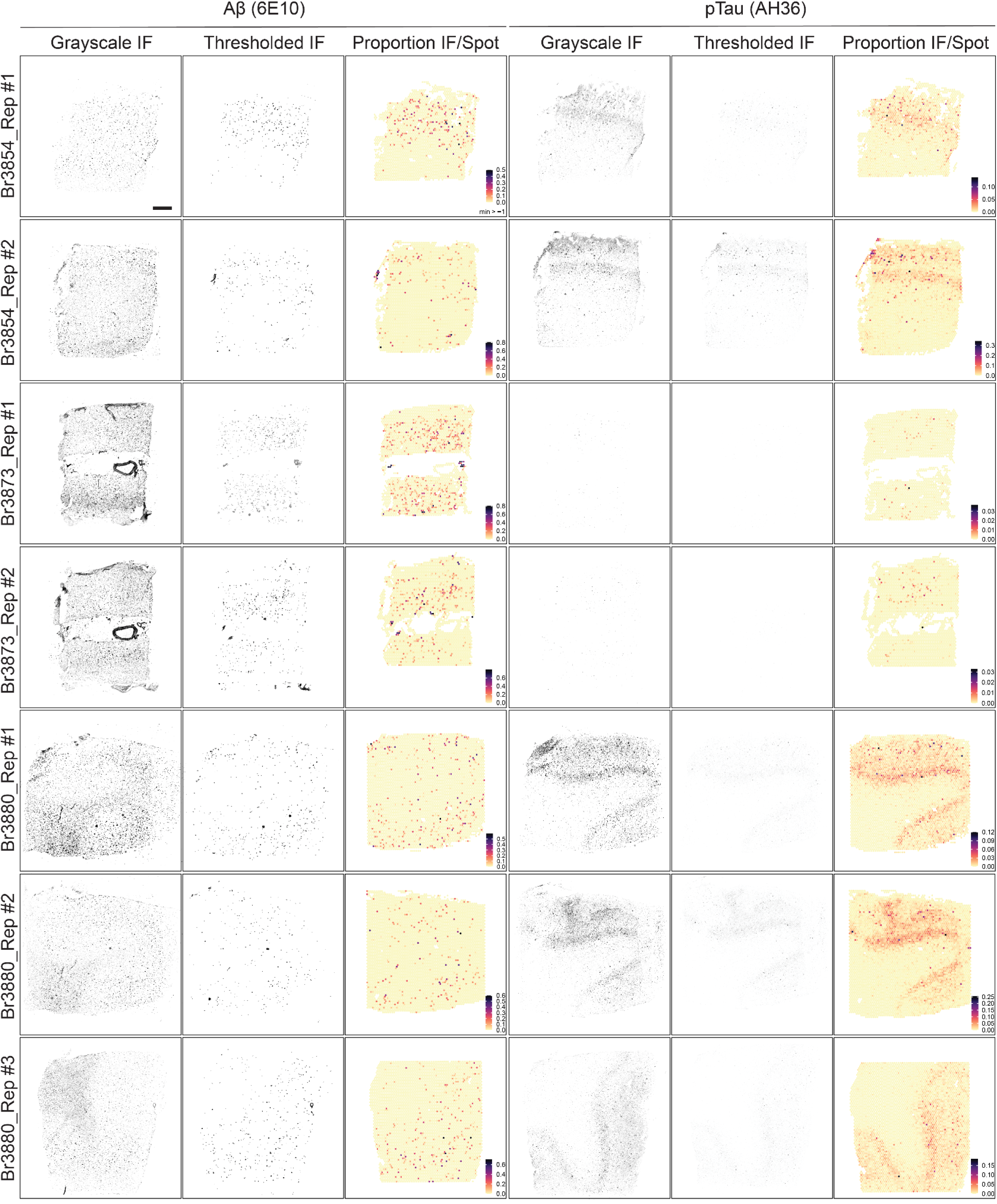
Immunofluorescence image thresholding and integration with spatially-resolved transcriptomics data. Raw grayscale immunofluorescent (IF) images of Aβ and pTau signals were segmented and thresholded by size and shape to remove autofluorescent signals and noises due to lipofuscin and ribonucleoside vanadyl complex (RVC), a ribonuclease inhibitor used in the staining protocol. 6E10 and AH36 refer to clonal names of the monoclonal antibodies raised against Aβ and pTau, respectively. Thresholded segmentations of image data were then integrated with complementary spatially resolved spot-level transcriptomes (i.e., Visium spots) to construct transcriptome-scale spatial maps of AD-related neuropathology in human ITC tissue sections across 3 AD donors. Color scale indicates the proportion of Aβ and pTau pixels per Visium spot. Scale bar, 1mm.

**Figure S7.**
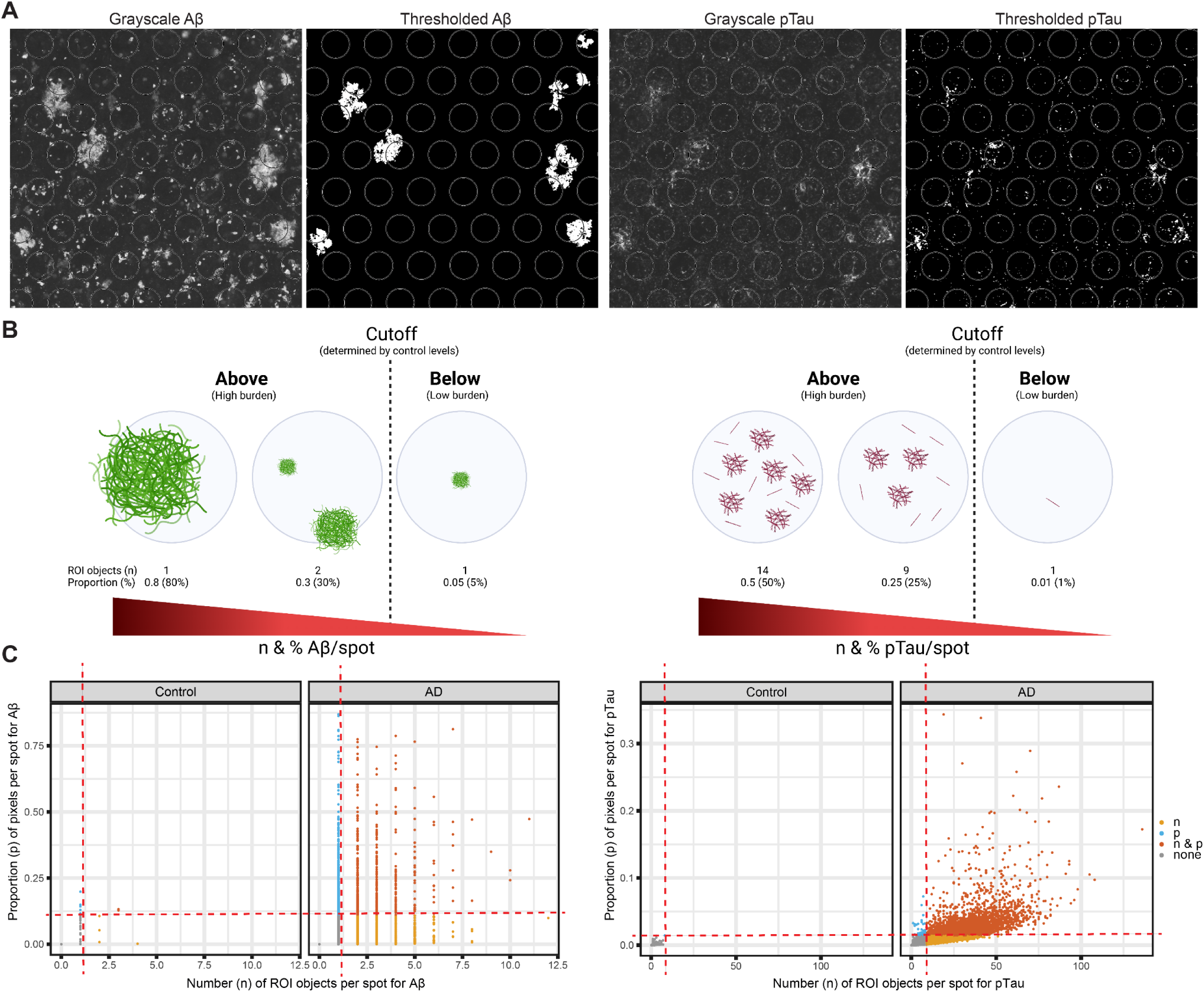
Identification of high-burden Aβ and pTau spots included in downstream analyses. (**A**) Representative raw grayscale IF and segmented/thresholded images of Aβ (left) and pTau (right) mapped to spot-level transcriptomes on the Visium platform. Diameter of Visium spot, 55µm. (**B**) Schematic cartoon illustrating how AD-related neuropathology was thresholded at the spot level. For individual Aβ (left) and pTau (right) pixels mapped to Visium spots, the number (n) of their region-of-interest (ROI) objects and proportion (p) of their pixels within ROIs were counted per spot. Cut-offs for each pathology were determined based on the values calculated from control tissue sections, which largely lacked pathology and were therefore representative of background signals. Visium spots containing the high pathological burden passing the defined threshold in both metrics (number of objects and proportion of pixels) were included in downstream analyses. (**C**) Distribution of Visium spots bearing varying degrees of AD-related pathology. Visium spots from 3 control (1 neurotypical donor) and 7 AD tissue sections (3 AD donors) were classified separately for Aβ (left) and pTau (right) using two different metrics: the number (n) of ROI objects and proportion (p) of pixels. Two thresholds (red dashed lines) were set with respect to the baseline levels observed in samples V10A27004_A1_Br3874 and V10T31036_A1_Br3874 from the neurotypical control donor. For Aβ, the cut-off values were 1 for (n) and 0.108 for (p); for pTau, 8 for (n) and 0.0143 for (p). Visium spots passing both (orange) or either of the thresholds (yellow and blue) were selected as heavy-burden spots for downstream analyses. Visium spots containing pathology below the thresholds were labeled gray indicating no significant pathology (none).

**Figure S8.**
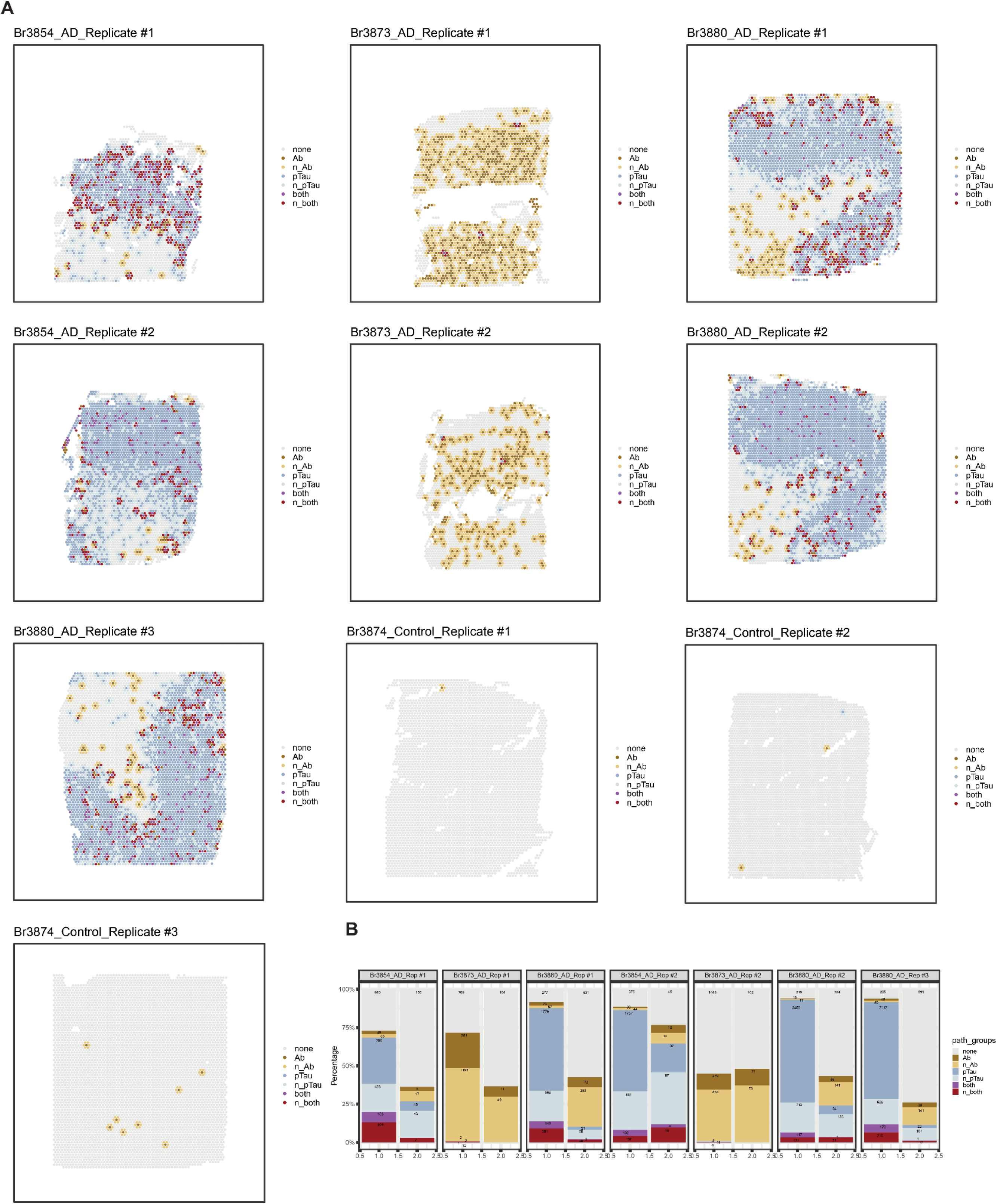
Classification of spots into different AD-related neuropathological categories. (**A**) Spot-level annotations of AD-related neuropathology across 10 tissue sections. The legend on the right indicates 7 spatial categories of AD-related neuropathology: none for no significant pathology; Ab for Aβ pathology; n_Ab for next to Aβ-containing spots, pTau for pTau pathology; n_pTau for next to pTau-containing spots; both for combined Aβ and pTau pathology; n_both for next to both. (**B**) Composition of annotated Visium spots constituting various spatial patterns of AD-related neuropathology for each AD tissue section across the 7 AD tissue sections used in the downstream analyses. The first column in each sample represents spots residing in the gray matter cluster and the second, the white matter cluster as identified by unsupervised clustering with *BayesSpace*, shown in **Figure S11**.

**Figure S9.**
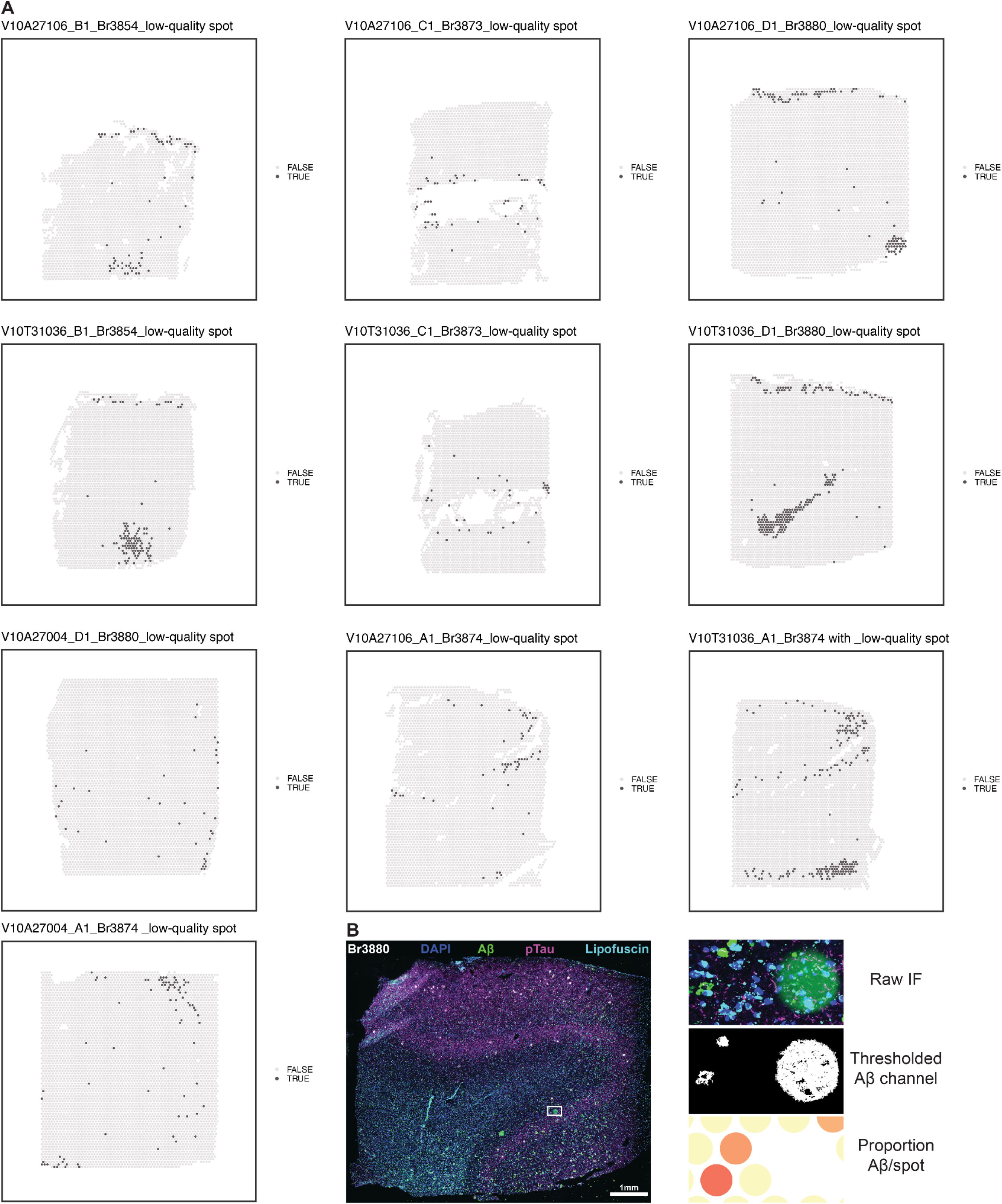
Spot-level quality control of Visium SRT data. (**A**) Visium spots with low library sizes, low gene expression features, or high mitochondria reads were identified by *scran* for all 10 samples and labeled in dark gray as TRUE. These spots in the 7 AD samples were retained in the downstream analyses due to their biological implication in the laminar organization of AD. (**B**) Manual removal of Visium spots with glare artifacts in the Aβ channel. The raw IF image of a tissue section from Br3880 (left) contains imaging-related glare artifacts in the green channel for Aβ detection (white rectangle). The artifacts were visually identifiable by shape, size, and translucence (right top, raw IF), but remained after image thresholding (right middle, thresholded Aβ channel). Hence, impacted Visium spots were manually identified and excluded using their coordinates on the Visium platform (right bottom, proportion Aβ/spot).

**Figure S10.**
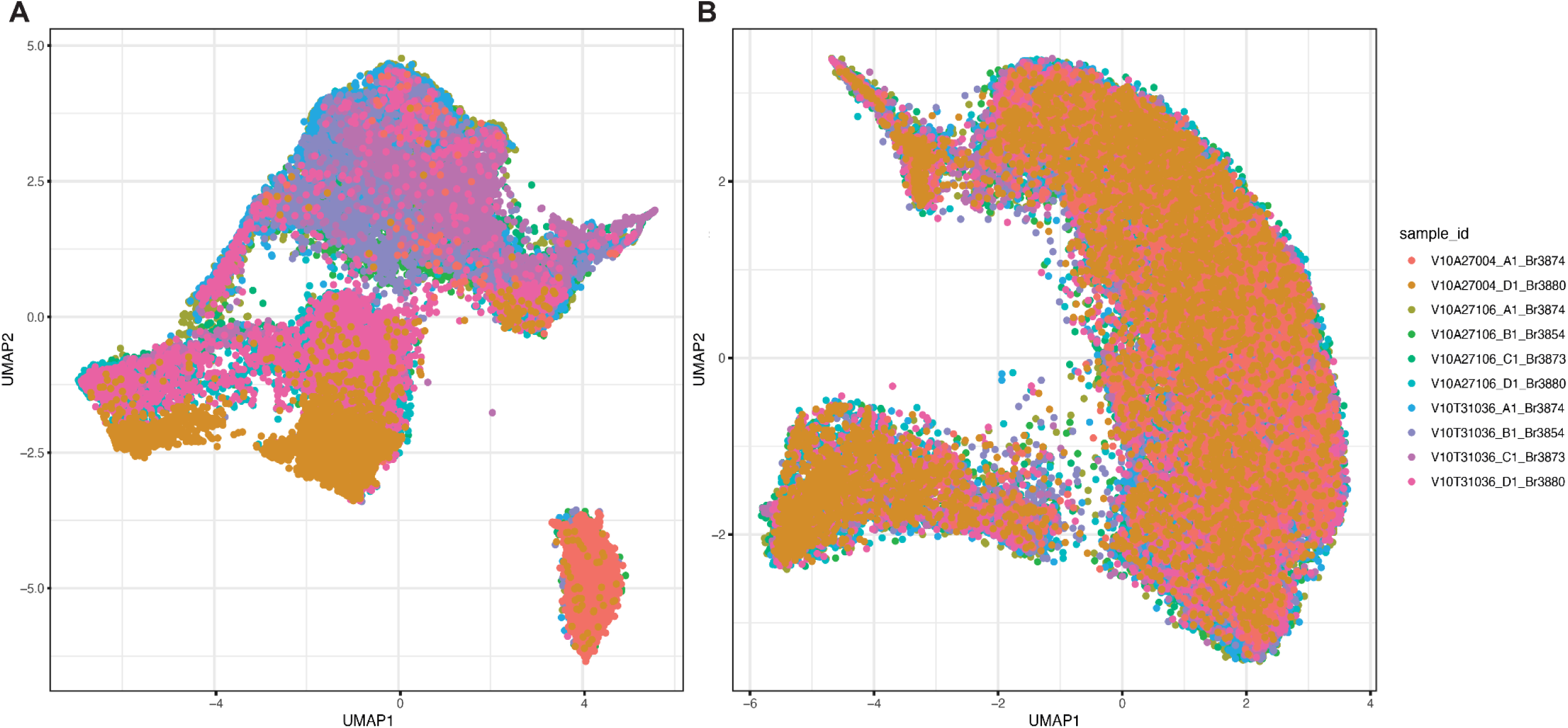
***Harmony*-based batch correction of Visium SRT data.** (**A**) UMAP representation of Visium spots aggregated across 10 tissue sections illustrates the effects of donor and technical variability, colored by sample IDs. (**B**) UMAP on *Harmony*-corrected PCs shows that donor effects and technical variations across multiple Visium-SPG experiments have been adjusted for.

**Figure S11.**
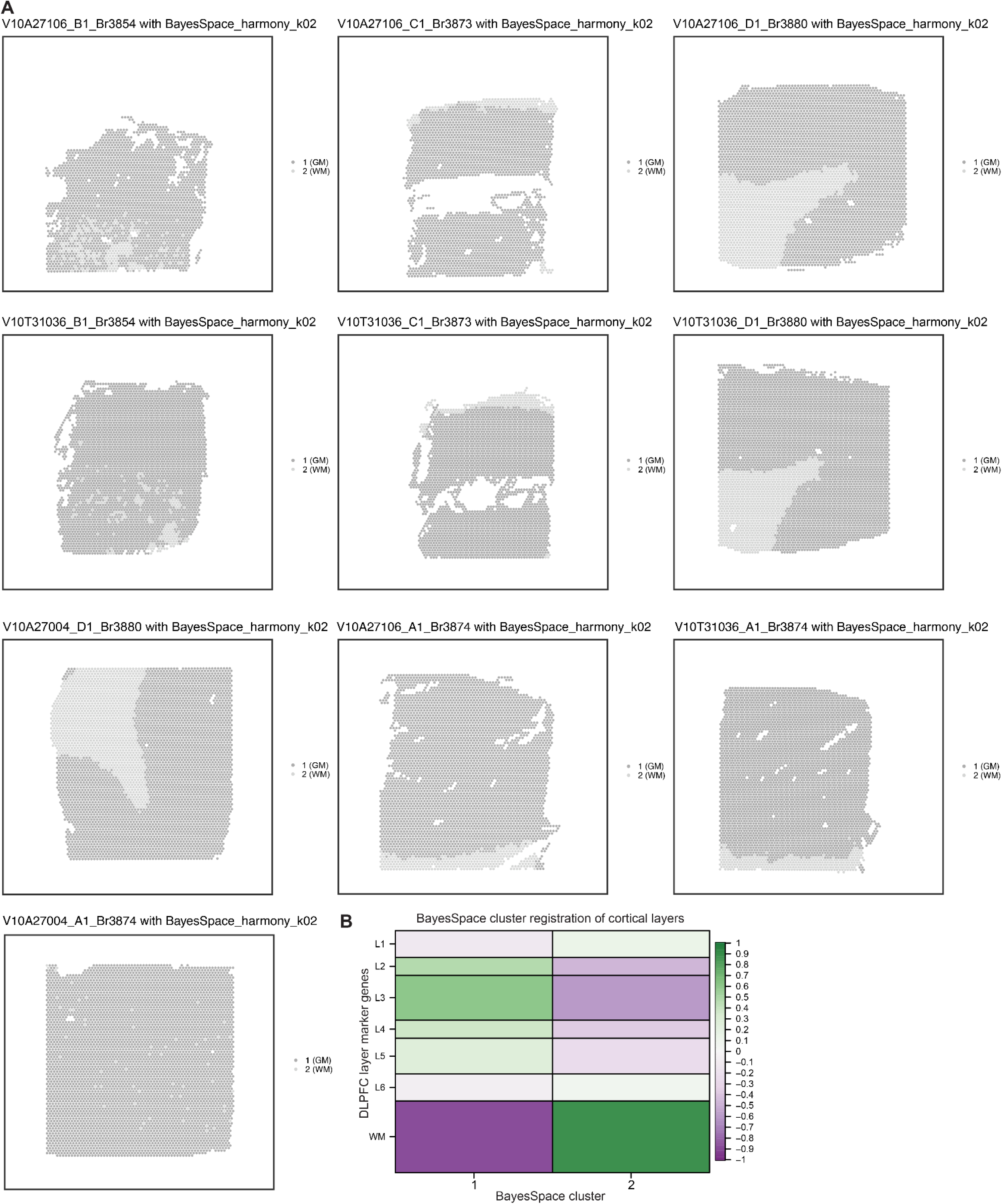
Unsupervised clustering with *BayesSpace* separates white and gray matter clusters in all tissue sections. (**A**) Spotplots of 10 tissue sections depicting the cluster labels for gray matter (1, GM, dark gray) and white matter (2, WM, light gray) identified in a data-driven manner with the *BayesSpace* clustering method at *k*=2. (**B**) Spatial registration heatmap illustrating the correlation between enrichment *t*-statistics calculated on *BayeSpace* clusters at *k*=2 on x-axis and those from manually annotated layers from Maynard, Collado-Torres, et al. 2021 on the y-axis. *BayesSpace* cluster 1 is associated with the gray matter based on enrichment of genes associated with cortical layers 2-5 in DLPFC.

**Figure S12.**
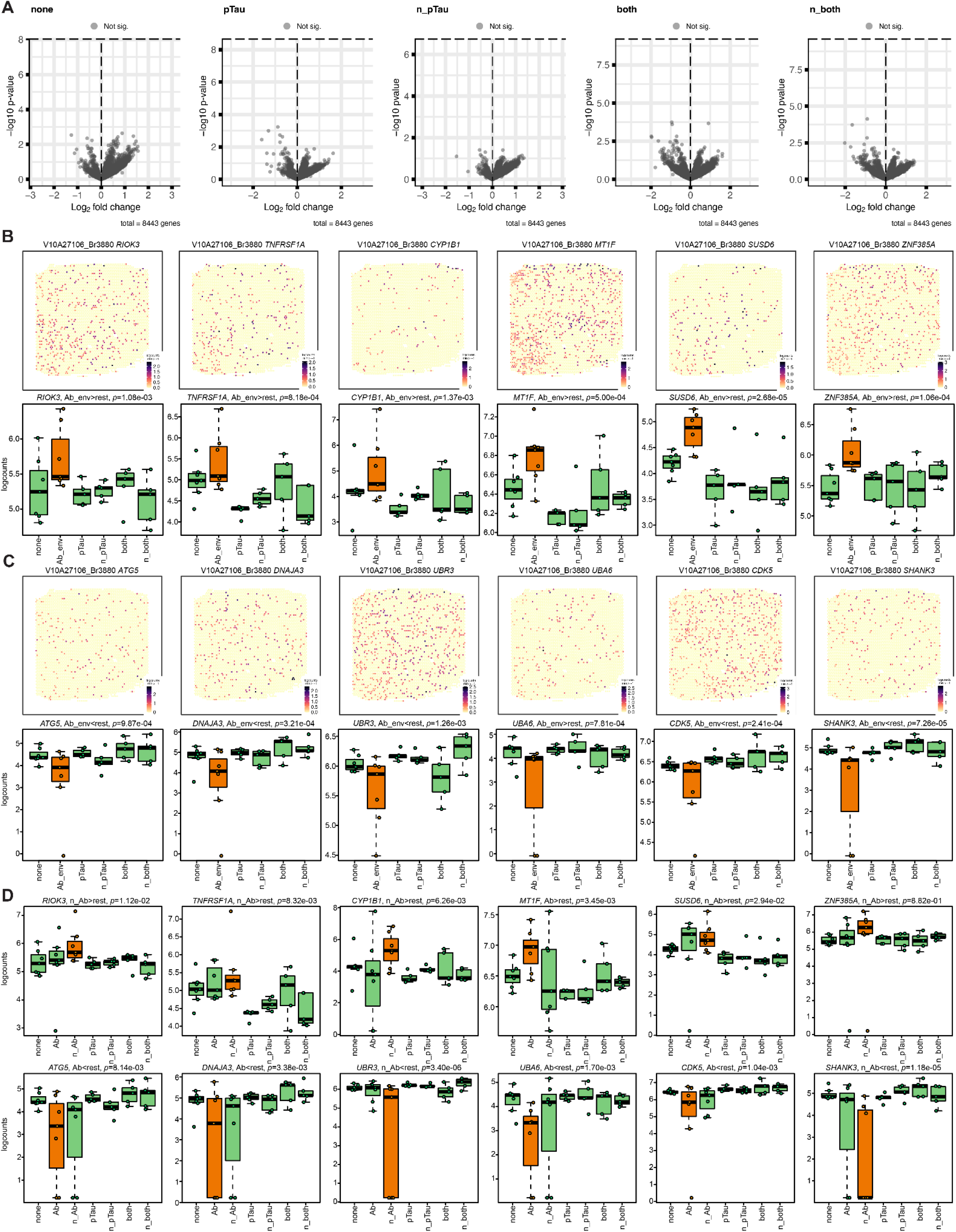
Inspecting biological functions and pathways of AD-associated microenvironments. (**A**) Volcano plots depicting DEGs in 5 other AD-related pathological categories based on the enrichment model. Each dot represents a gene, plotted with its log2 fold change (x-axis) and -log10 *p*-value (y-axis), thus comparing the effect size (fold change) against the statistical evidence for differential expression (*p*-value). The dashed line represents a *p*-value threshold matching FDR<0.1, with DEGs below the line considered not significant (gray, Not sig.) No DEGs (FDR<0.1) were identified for these 5 pathological categories. For brevity, ‘next’ was shortened to n. (**B**) Spotplots (top) and boxplots (bottom) of 6 DEGs enriched in the extended Aβ-associated microenvironment (Ab_env). Aβ spots and Aβ-adjacent spots (next_Aβ) were collapsed into a single pathological category to represent the extended Aβ-associated microenvironment (Ab_env), which was then subjected to differential expression testing based on the enrichment model. Spotplots are from Br3880 and color scale indicates spot-level gene expression in logcounts. Boxplots display gene expression levels (logcounts) for all samples and highlight differential expression between one pathological category (orange) versus all other categories (green). For brevity, ‘next’ was shortened to n. (**C**) Spotplots (top) and boxplots (bottom) of 6 DEGs depleted in the extended Aβ-associated microenvironment (Ab_env). Spotplots are from Br3880 and color scale indicates spot-level gene expression in logcounts. Boxplots display gene expression levels (logcounts) for all samples and highlight differential expression between one pathological category (orange) versus all other categories (green). (**D**) Boxplots of the 12 DEGs associated with the extended Aβ-associated microenvironment (Ab_env) listed in **Figure S12B-C** to indicate their differential gene expression levels across the previously classified 7 pathological categories where Ab_env was spilt into the Aβ and next_Aβ categories. Boxplots display gene expression levels (logcounts) for all samples and highlight differential expression between one pathological category (orange) versus all other categories (green). For brevity, ‘next’ was shortened to n.

**Figure S13.**
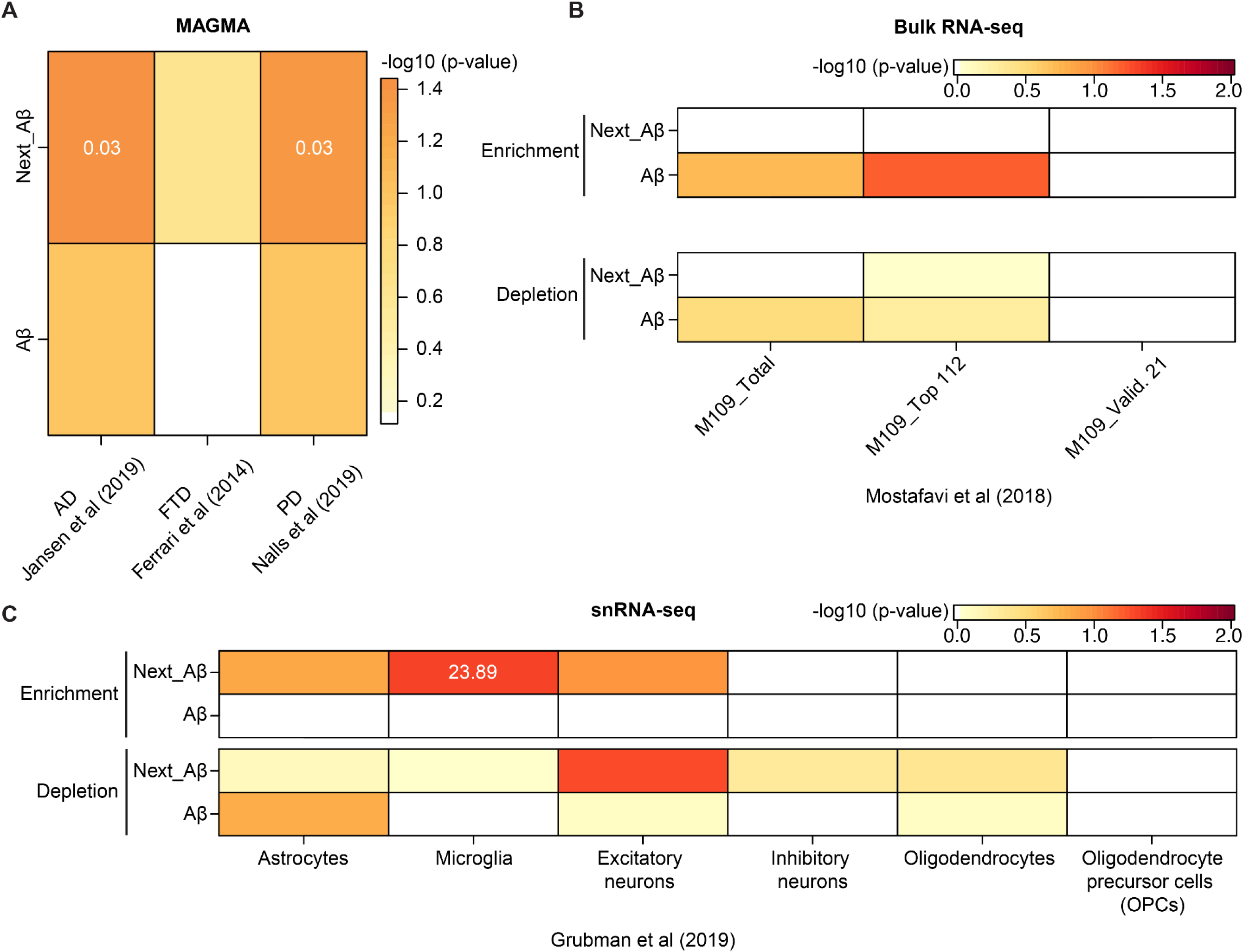
Enrichment of neurodegeneration- and dementia-related gene sets in Aβ-associated microenvironments. (**A**) MAGMA associations of GWAS datasets for Alzheimer’s disease (AD, Jansen et al. 2019), frontotemporal dementia (FTD, Ferrari et al. 2014), and Parkinson’s disease (PD, Nalls et al. 2019) with Aβ-associated microenvironments profiled in human ITC. The results were presented in a heatmap, with the color scale indicating −log10 (*p*-value) significance of the association test. The number in each box indicates the effect size (beta) for significant associations (*p*<0.05). (**B**) Spatial gene set enrichment analysis with a bulk RNA-seq dataset of dorsolateral prefrontal cortex (DLPFC) collected from the ROSMAP cohort (Mostafavi et al. 2018). M109_Total refers to a gene module of 390 co-expressed genes strongly associated with cognitive decline and Aβ pathology in AD. M109_Top 112 is a subset of M109 containing top 112 interesting genes selected by an initial Bayesian network, while M109_Valid 21 is a gene set containing 21 validated genes. These gene sets were compared and tested for their associations with Aβ-associated microenvironments profiled for enriched (top) and depleted (bottom) gene sets in the human ITC. The results are summarized in a heatmap, with the color scale indicating −log10 (*p*-value). (**C**) Spatial gene set enrichment analysis with a snRNA-seq dataset for AD (Grubman et al. 2019). Grubman et al. identified AD-associated DEGs between case and control across 6 different cell types in the human entorhinal cortex, which were found to overlap with those identified in human DLPFC (Mathys et al. 2019). These cell type-specific gene sets were compared and tested for their associations with the Aβ-associated microenvironment in the human ITC, separated by enriched (top) and depleted (bottom) gene sets. The results are summarized in a heatmap, with the color scale indicating −log10 (*p*-value) and the numbers in the boxes indicating odd ratios for the enrichment.

**Figure S14.**
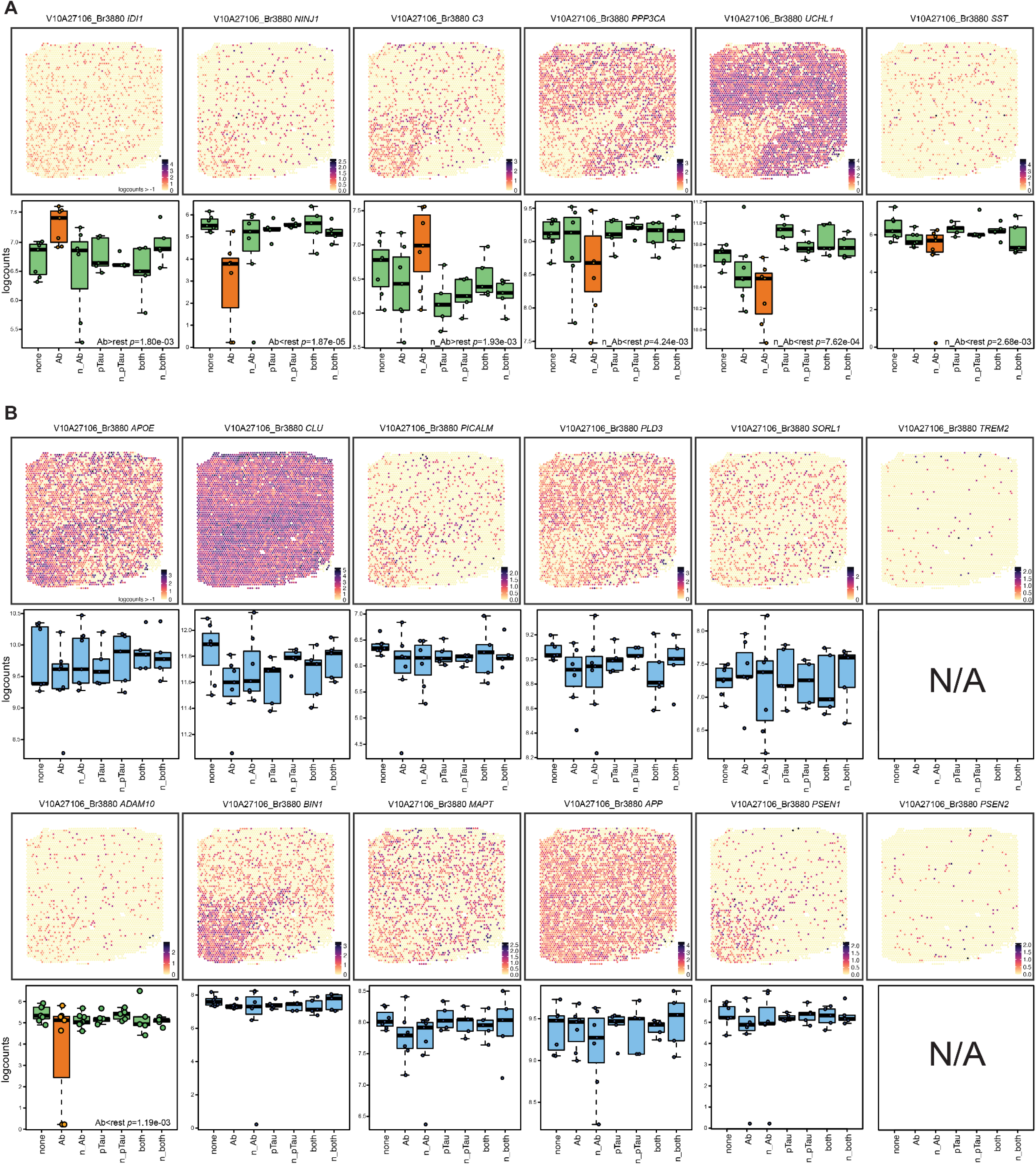
Spatial expression of Aβ-associated genes and AD risk genes. (**A**) Spotplots (top) and boxplots (bottom) of 6 genes differentially expressed in Aβ and Aβ-adjacent (next_Aβ;n_Ab) pathological categories. Spotplots are from Br3880 and color scale indicates spot-level gene expression in logcounts. Boxplots display gene expression levels (logcounts) for all samples and highlight differential expression between one pathological category (orange) versus all other categories (green). For brevity, ‘next’ was shortened to n. (**B**) Spotplots (top) and boxplots (bottom) of selected AD-associated risk genes identified from the literature. Except for *ADAM10* (orange-green color scheme), none of these risk genes were significantly enriched or depleted across the 7 spatial categories of AD-related neuropathology (blue color scheme). *TREM2* and *PSEN2* had low tissue-wide gene expression and were thus filtered out prior to performing differential expression analysis (N/A: not available). For brevity, ‘next’ was shortened to n. For spotplots, color scale indicates spot-level gene expression in logcounts.

**Figure S15.**
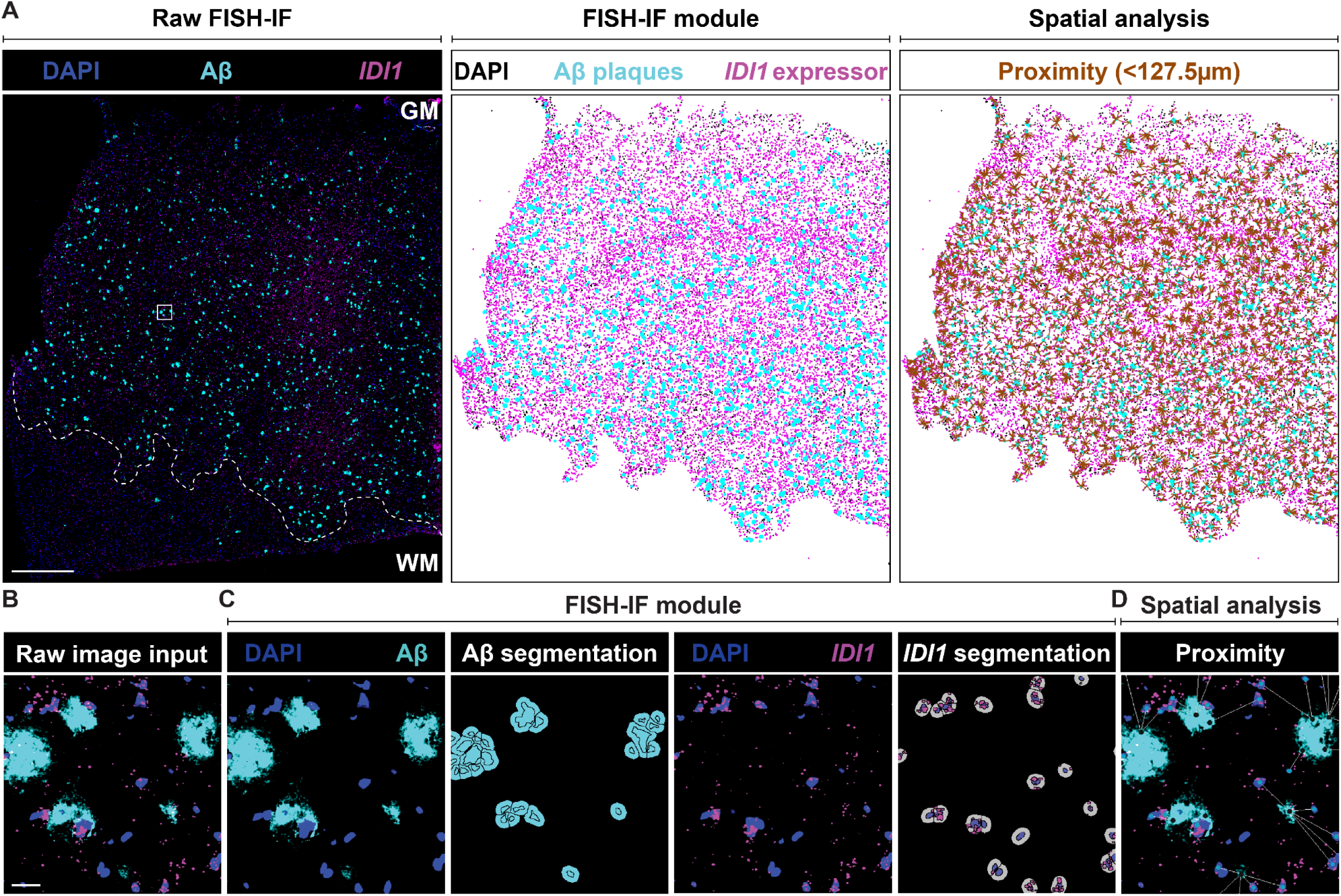
RNAscope FISH-IF image segmentation and quantification for spatial proximity analysis using HALO software. (**A**) Tissue-wide image segmentation and spatial analysis of Aβ and cells expressing GOIs (GOI expressors) using gene *IDI1* and Br3854 as a representative example. RNAscope FISH-IF staining image (left) showing the distribution of Aβ (cyan) and GOI expressors (*IDI1I* expressors; magenta) throughout the entire tissue section. Based on the high density of Aβ plaques, the gray matter (GM) was manually annotated (white dashed line) and selectively incorporated into downstream image analyses. The raw staining image was computationally processed using the FISH-IF module in the HALO image analysis software to segment Aβ plaques and *IDI1* expressors (middle). Subsequent spatial analysis measured the proximity between Aβ and *IDI1* expressors with proximity lines (brown) drawn between the segmented objects distanced less than 127.5µm, a predetermined radius based on the Visium spot grid that defined the Aβ-associated microenvironment. Scale bar, 1mm. (**B**) Higher-magnification raw input image of Aβ and *IDI1* expressors. Scale bar, 25µm. (**C**) Higher-magnification demonstration of FISH-IF module segmentation of Aβ plaques and *IDI1* expressors, independently. Due to the varied pixel intensities within Aβ plaques, a single Aβ plaque (cyan) was divided into multiple fragments upon segmentation (Aβ segmentation). The *IDI1* transcript signals (magenta) were segmented as individual puncta (*IDI1* segmentation), which were then counted to measure gene expression within single cells. *IDI1* expressors were identified based on the presence of the segmented *IDI1* signals in nuclei (blue) and estimated cytoplasm (gray). The puncta outside of the nucleus and cytoplasm was not able to be quantified with this module. (**D**) Higher-magnification view of spatial analysis to determine the proximity between the centroids of the segmented fragments of Aβ plaques and neighboring *IDI1* expressors, with proximity lines (white) drawn between objects at a maximum of 127.5µm apart.

**Figure S16.**
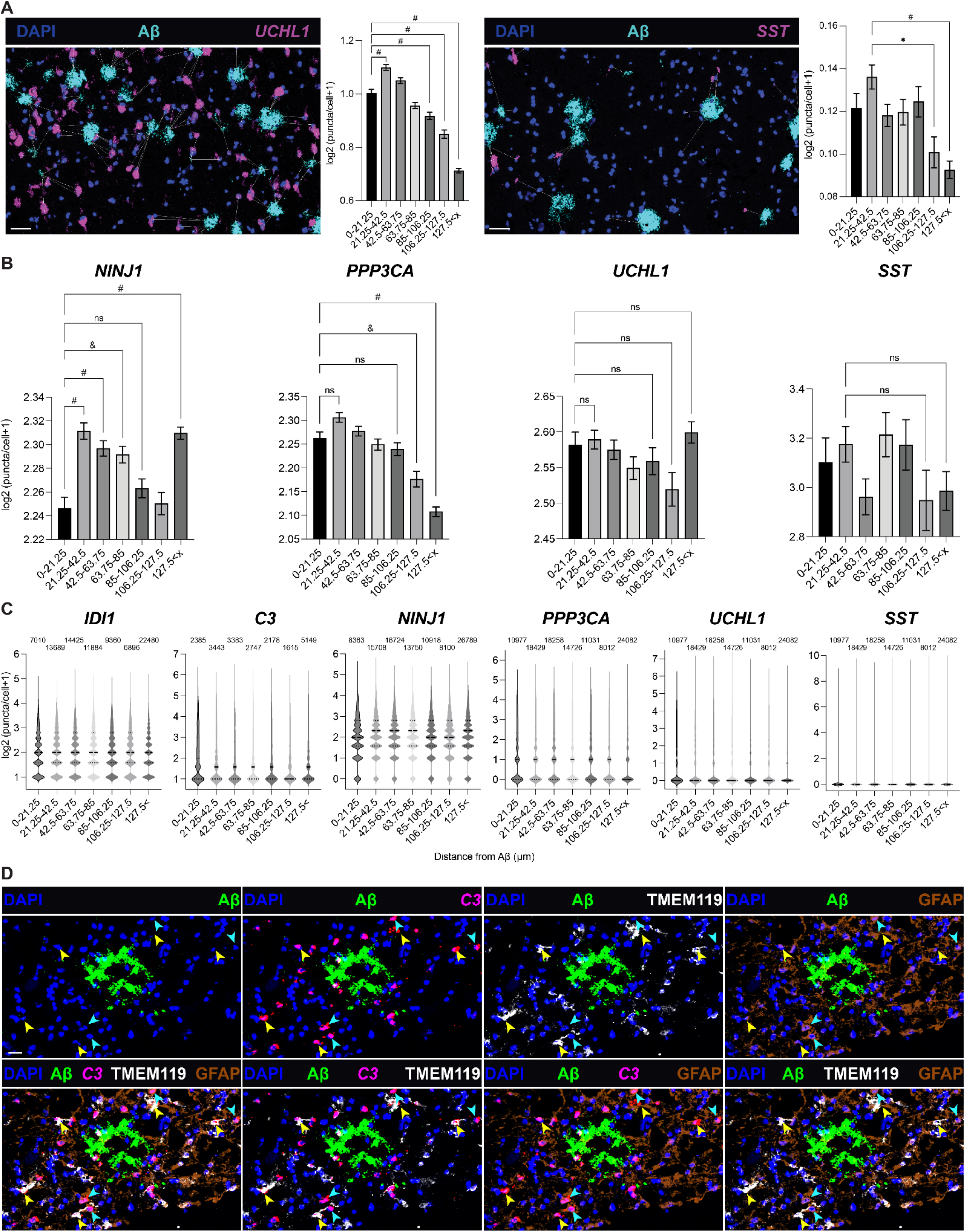
Validation of quantification strategies of Aβ-associated transcriptional signatures at cellular resolution. (**A**) RNA-protein co-detection of Aβ and two GOIs, *UCHL1* (left) and *SST* (right) showing the spatial expression of the GOIs (magenta) relative to nearby Aβ. Proximity lines indicate the distance between Aβ and nearby GOI expressors (max: 127.5µm). Scale bar, 50µm. Quantification of each GOI shows gene expression changes across the 7 bins with log2 (X+1) transformation where X indicates the puncta counts. A total of cells, including GOI expressors and non-expressors, were counted to normalize gene expression for each bin. The interval for each bin was labeled on the x-axis. Data are mean ± SEM (Kruskal-Wallis test, \**p*<0.05 and ^#^*p*<0.0001). (**B**) Bar plots illustrate gene expression changes for 4 GOIs, *NINJ1*, *PPP3CA*, *UCHL1*, and *SST* across 7 distance bins when normalization was performed only with GOI expressors, as in the box plots for *IDI1* and *C3* in Figure 3D. Gene expression levels were quantified with log2 (X+1) transformation where X indicates the puncta counts. The distance for each bin was specified on the x-axis. Data are mean ± SEM (Kruskal-Wallis test, ns=not significant, \**p*<0.05, ^&^*p*<0.005, and ^#^*p*<0.0001). (**C**) Violin plots showing the discrete distribution of GOI expressors with log2 (X+1) transformation where X indicates the puncta counts. The number labeled in each column indicates the total number of cells counted across the 7 bins and included for the quantification plots in Figure 3D and **Figure S16A**. For *IDI1* and *C3* (enriched genes), only the expressors were counted, while for *NINJ1*, *PPP3CA*, *UCHL1*, and *SST* (depleted genes), both expressors and non-expressors were counted to address potential depletion due to non-expressing cells which may artificially dilute gene expression by expressors. *PPP3CA*, *UCHL1*, and *SST* showed the same number of counted cells in every interval as their gene expression was stained and measured within the same tissue samples. (**D**) RNA-protein co-detection of Aβ (green), *C3* transcripts (magenta), TMEM119 (white), and GFAP (brown) to show enrichment of *C3* in TMEM119^+^ microglia (yellow arrowheads) and GFAP^+^ astrocytes (cyan arrowheads) in the vicinity of Aβ plaques. Scale bar, 20μm.

