## Supplementary figures and images for "Influence of Alzheimer’s disease related neuropathology on local microenvironment gene expression in the human inferior temporal cortex"

### FigureS1_TB&TO_300ppi.png

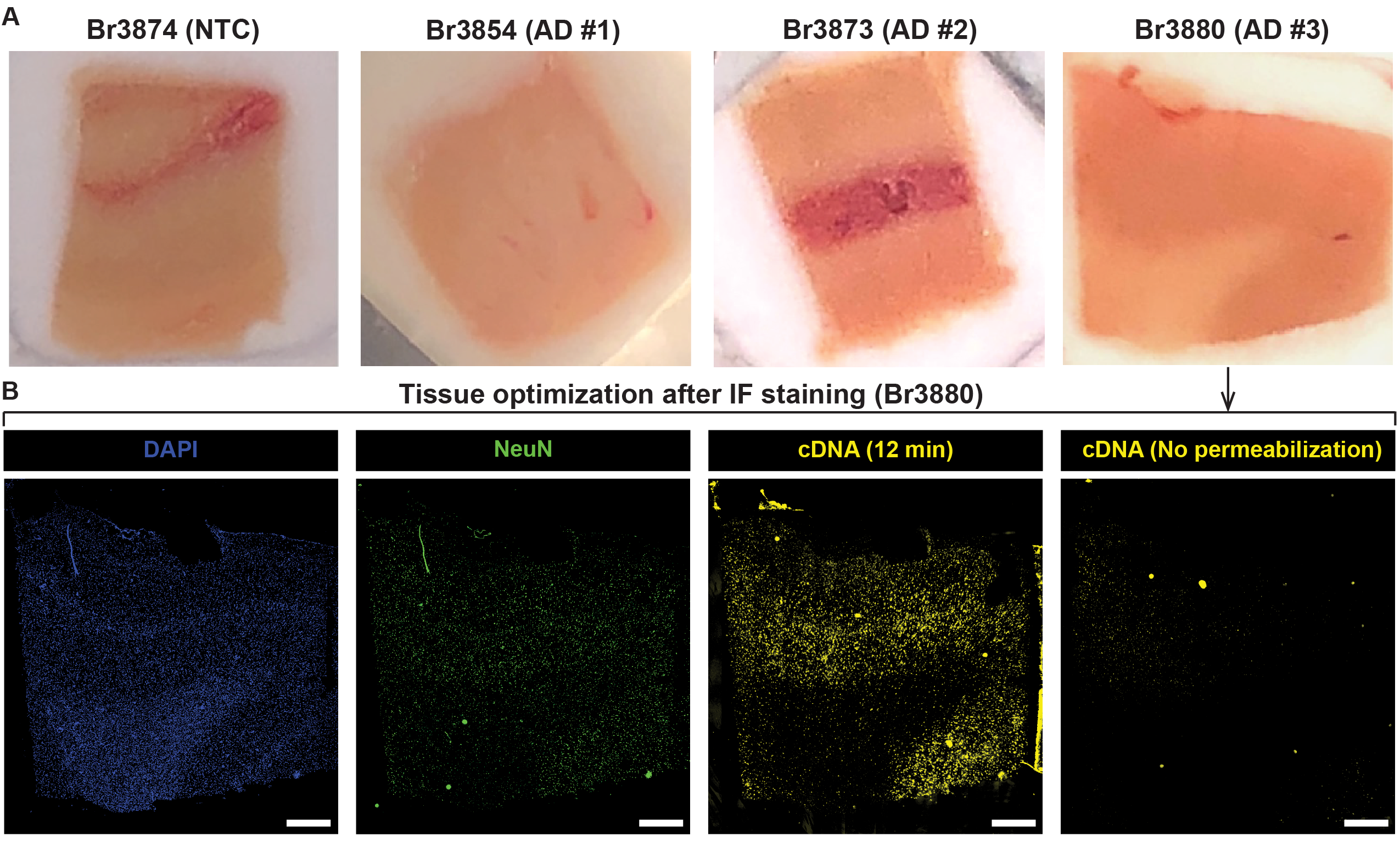

### FigureS2_HimagThresh_300ppi.png

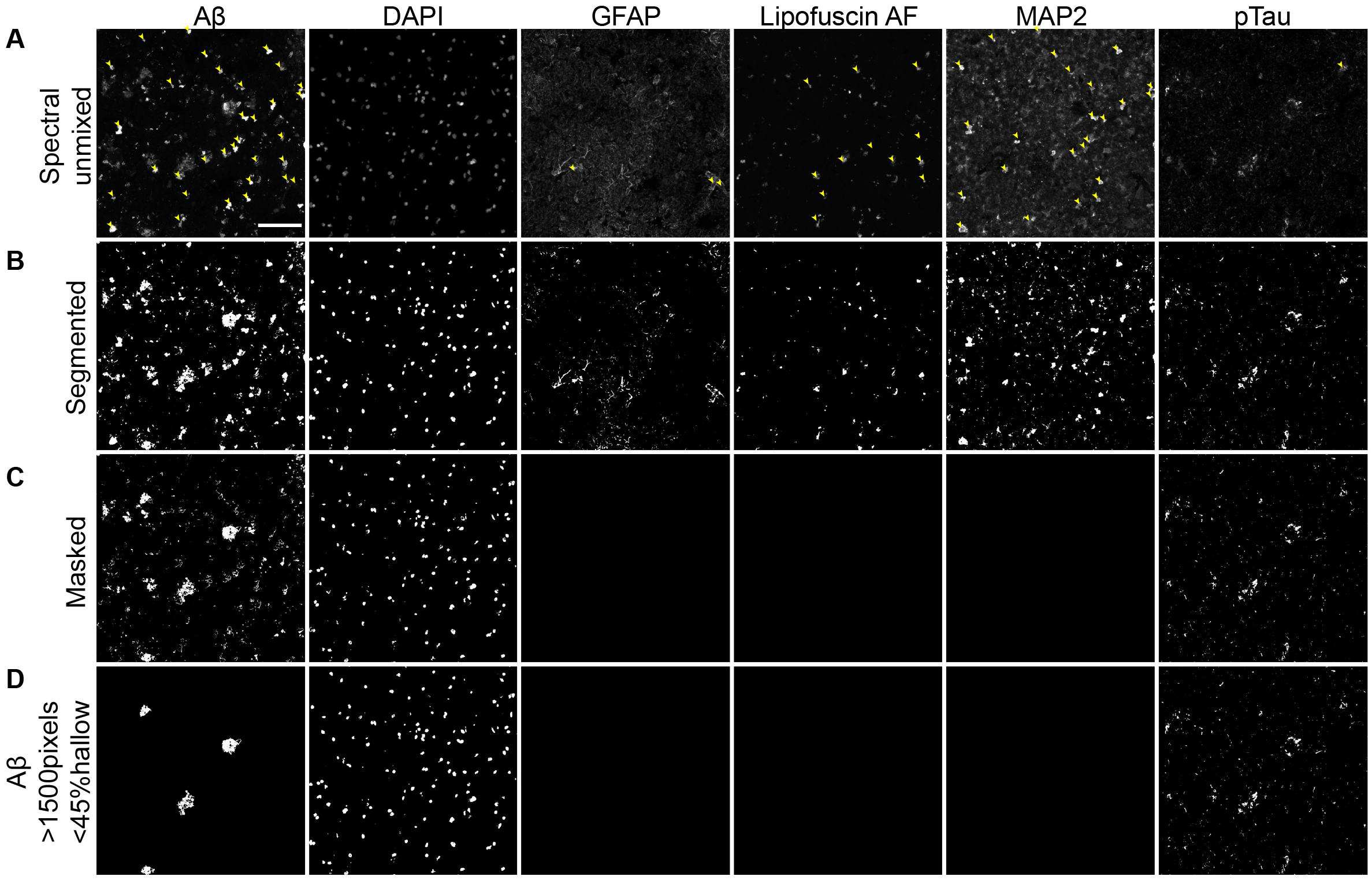

### FigureS3_CtrlBr3874_300ppi.png

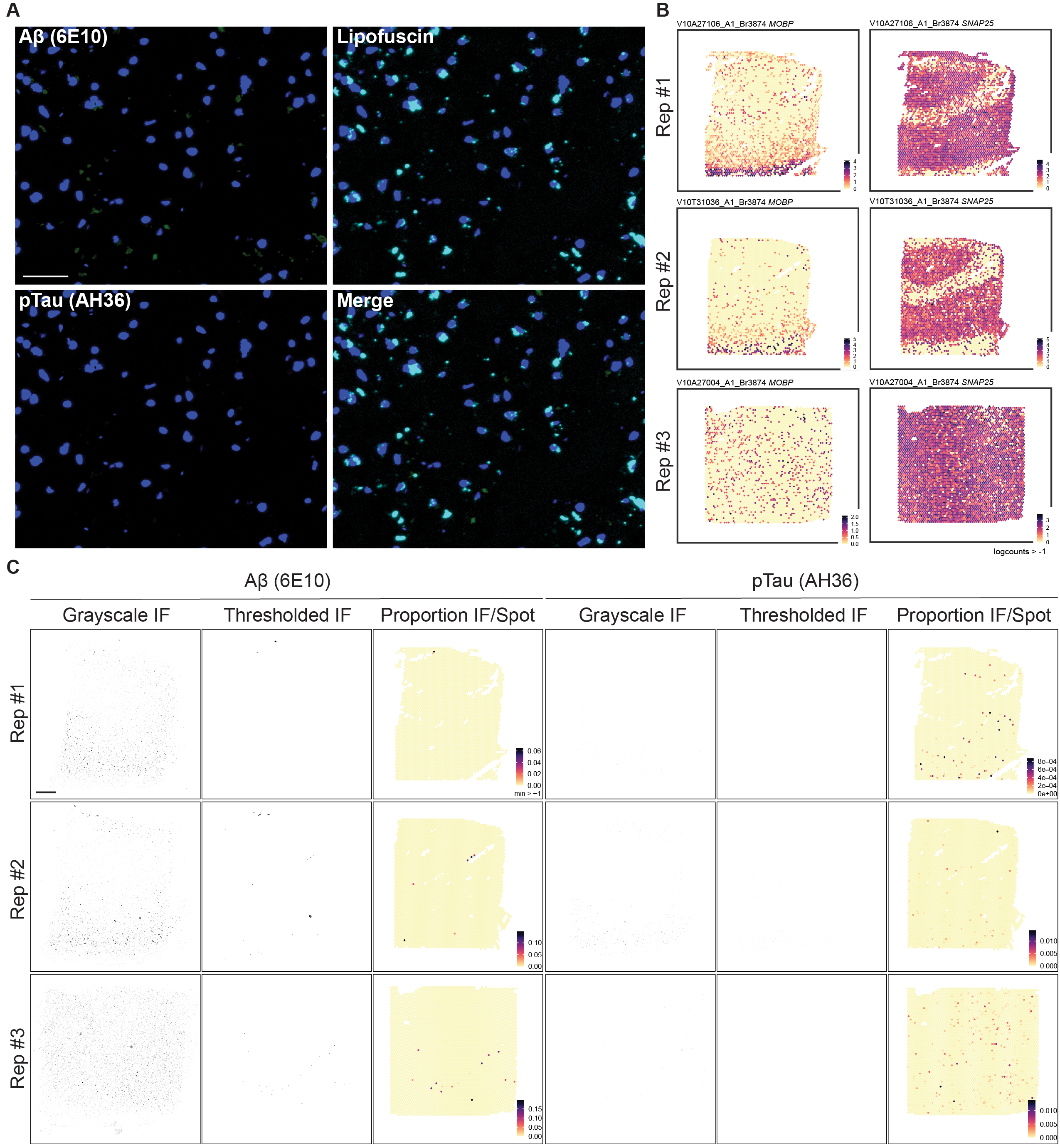

### FigureS4_MOBP_300ppi.png

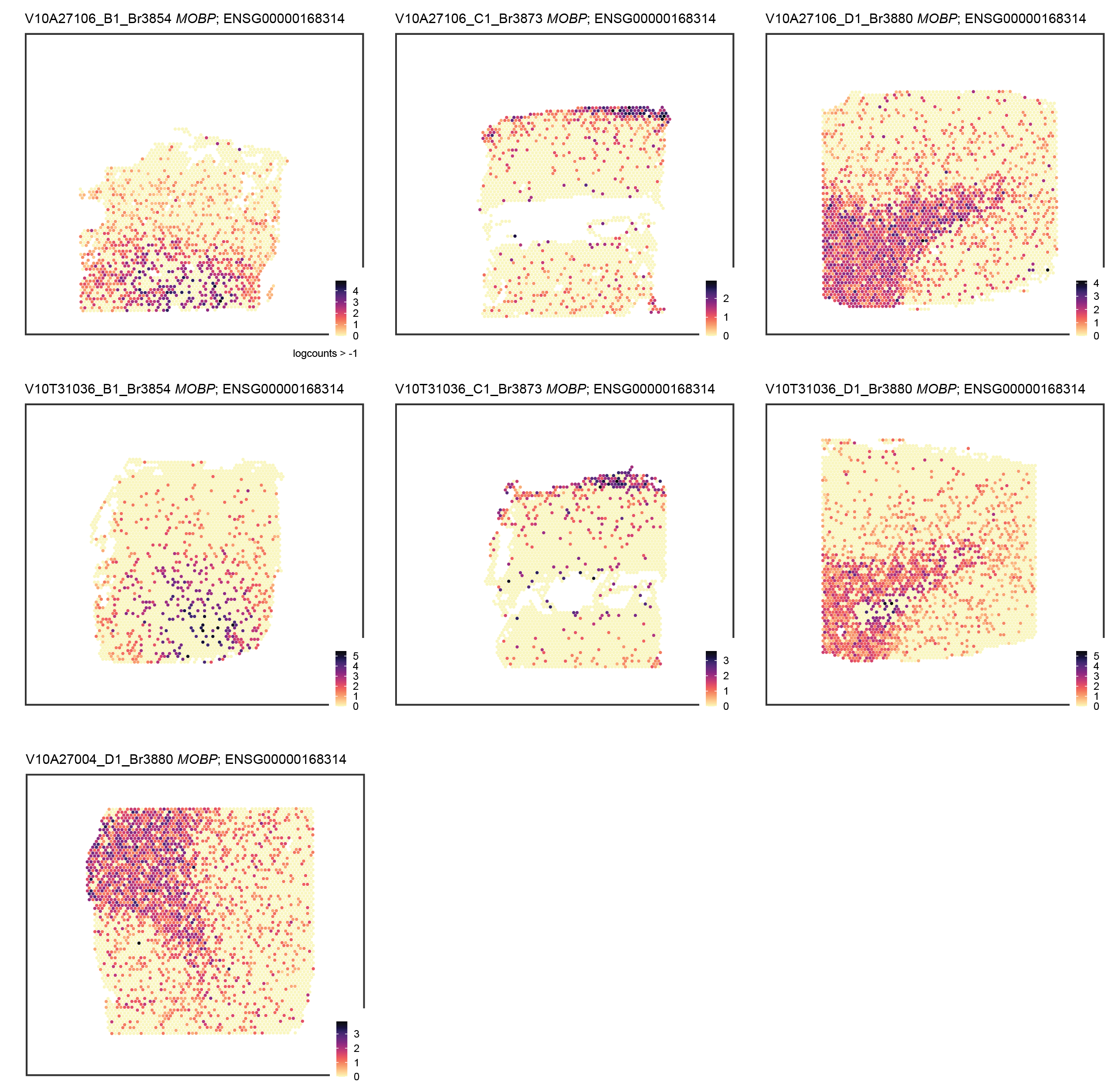

### FigureS5_SNAP25_300ppi.png

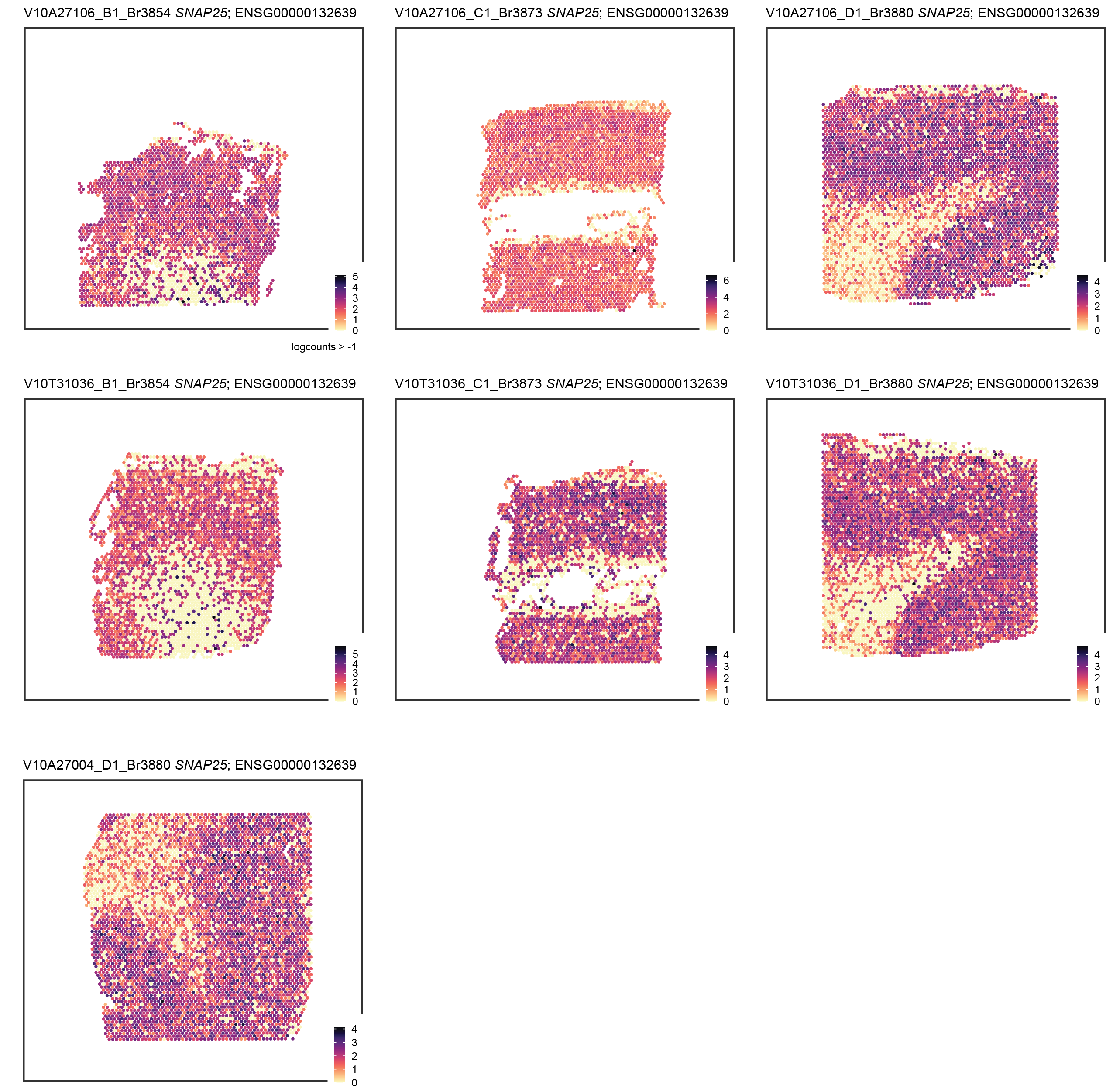

### FigureS6_TissueLevThreshAD_300ppi.png

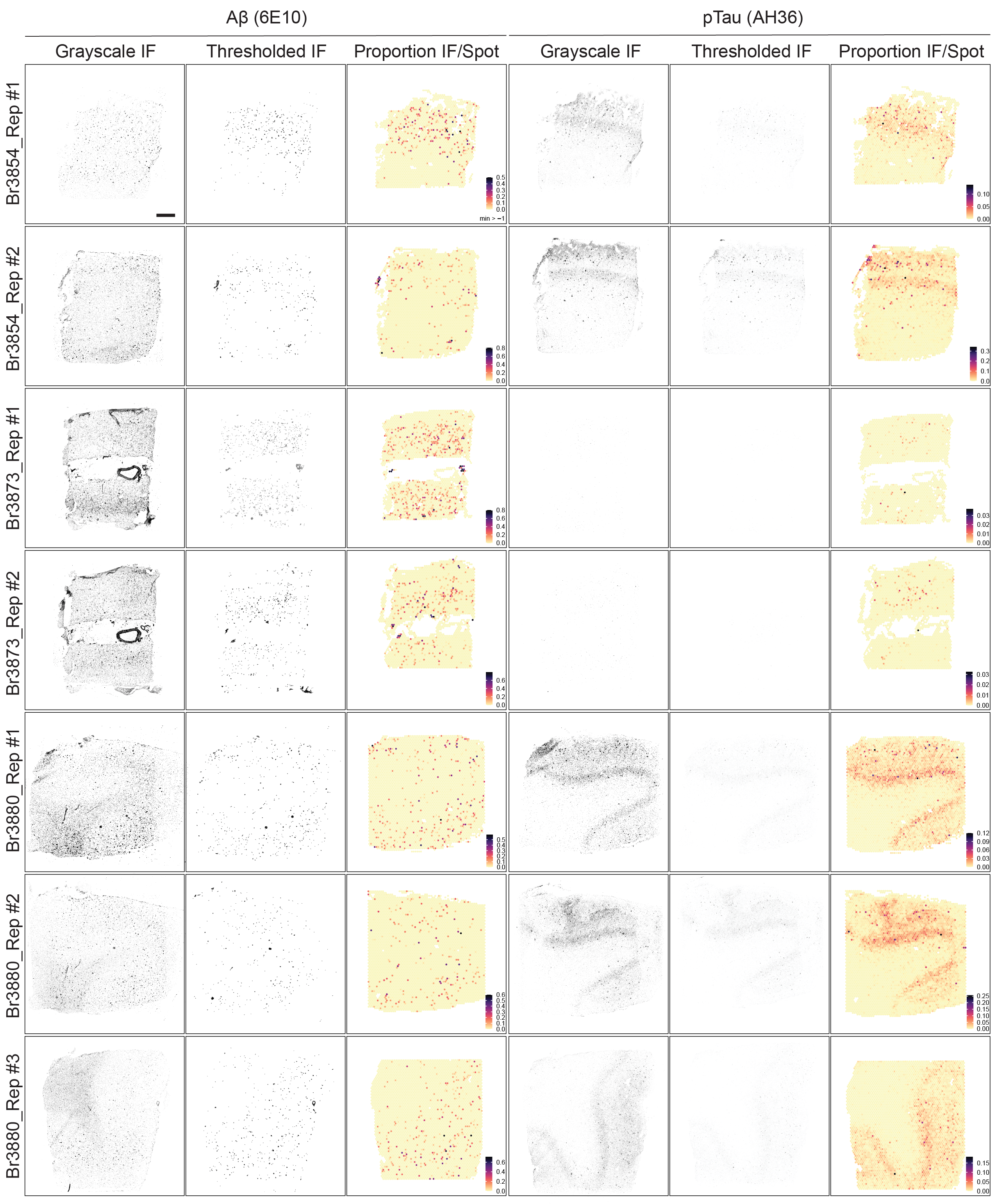

### FigureS7_HIBurdenSpots_300ppi.png

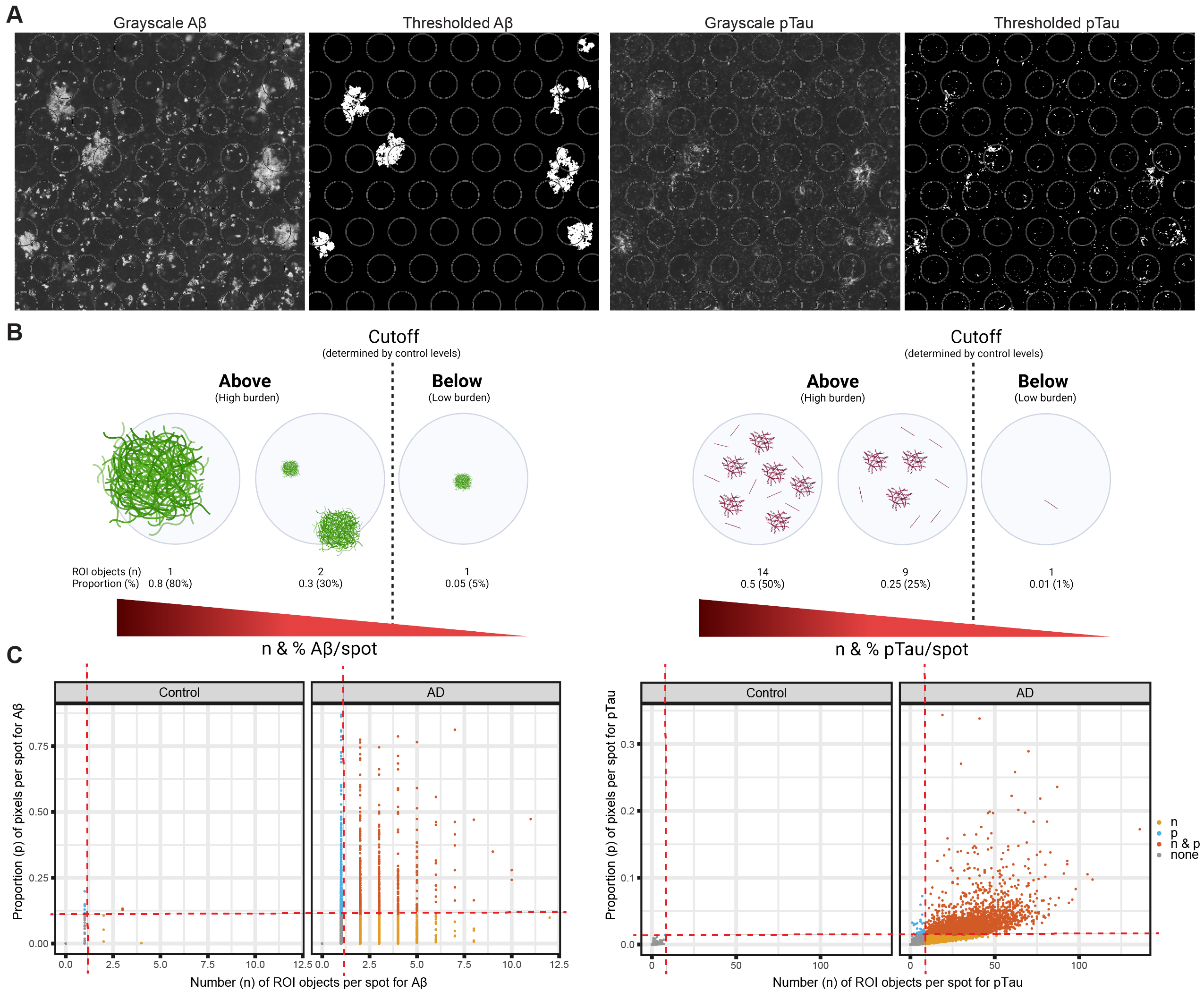

### FigureS8_SptLevAnnot_300ppi.png

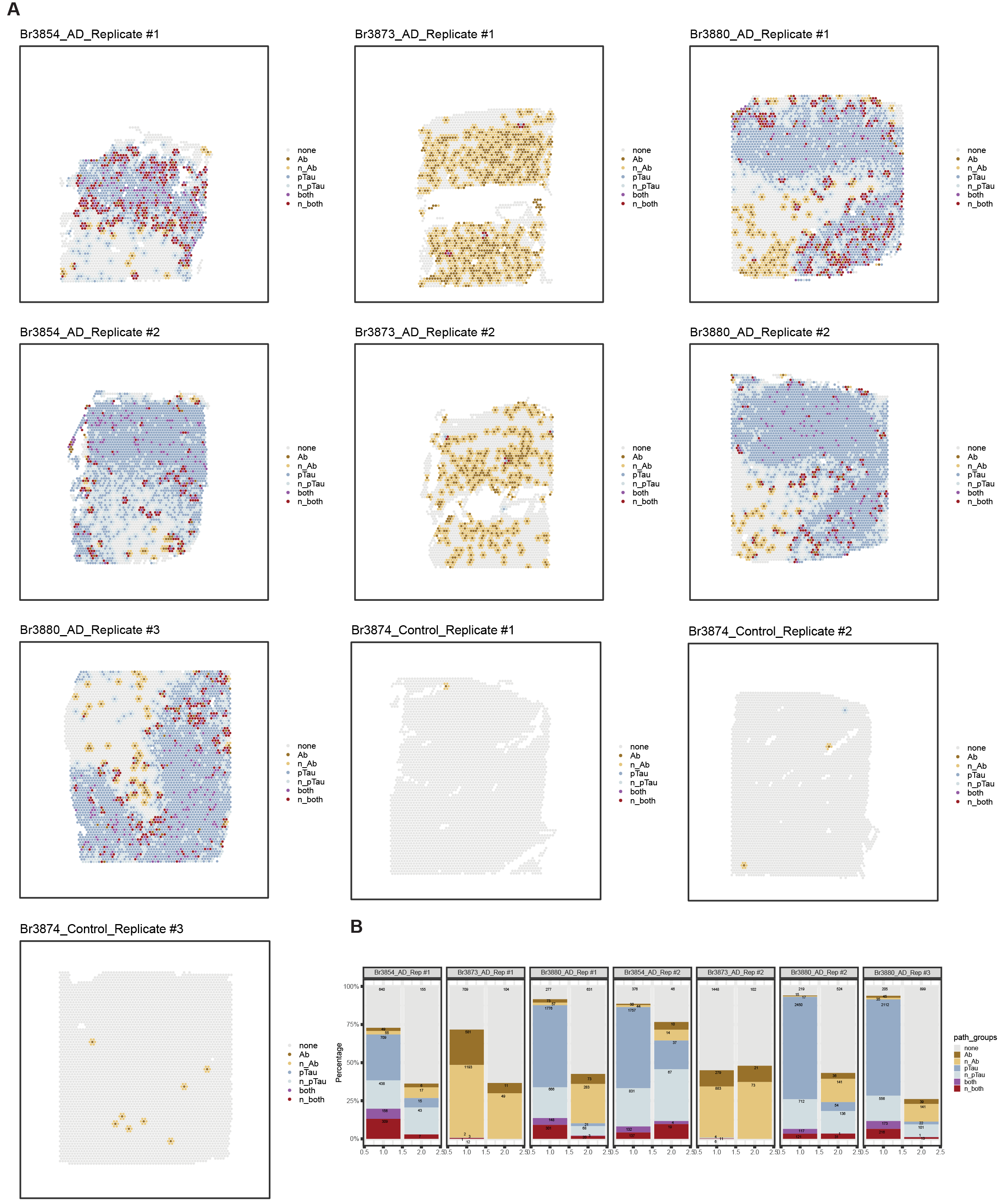

### FigureS9_SpotLevQC_300ppi.png

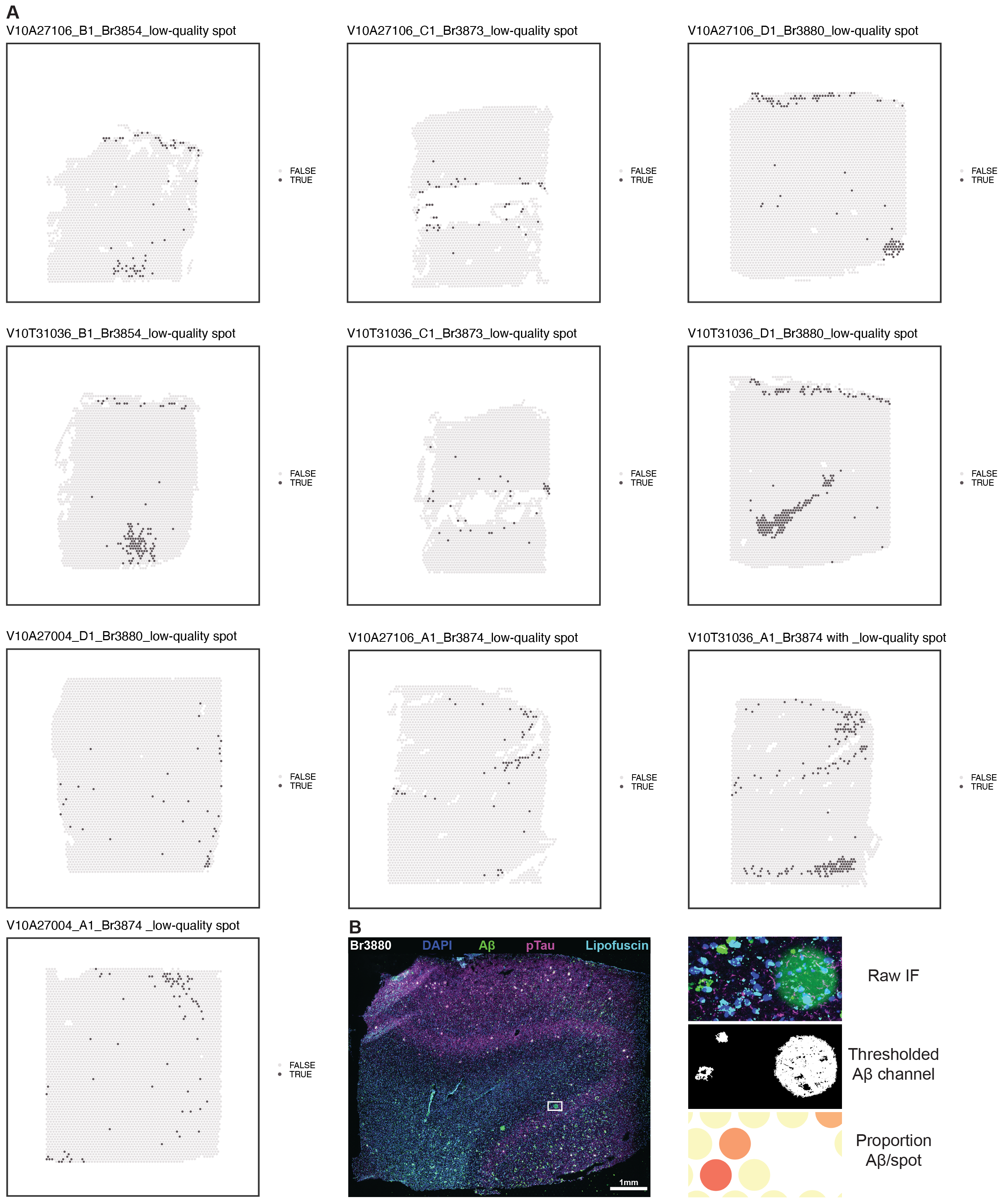

### FigureS10_Harmony_300ppi.png

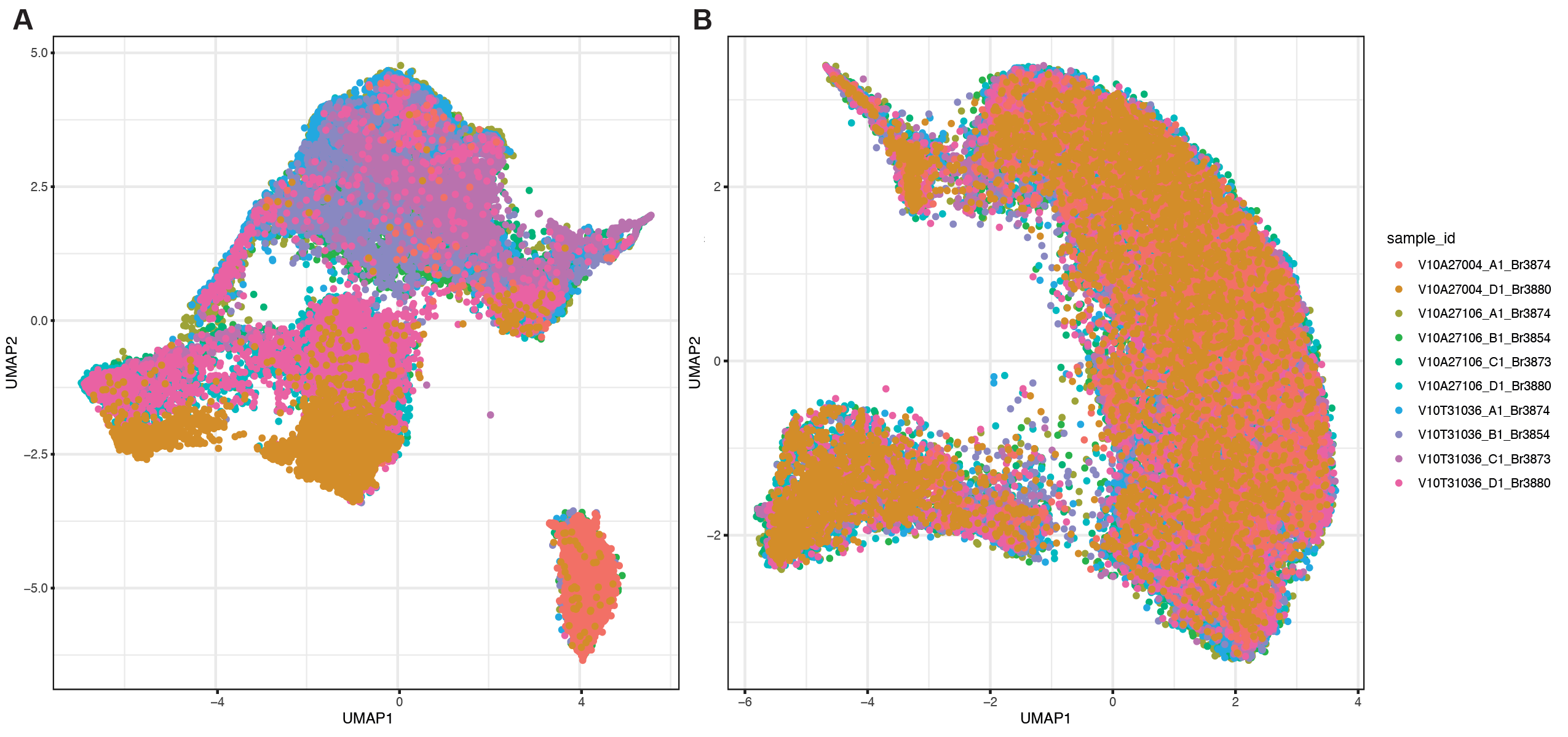

### FigureS11_300ppi.png

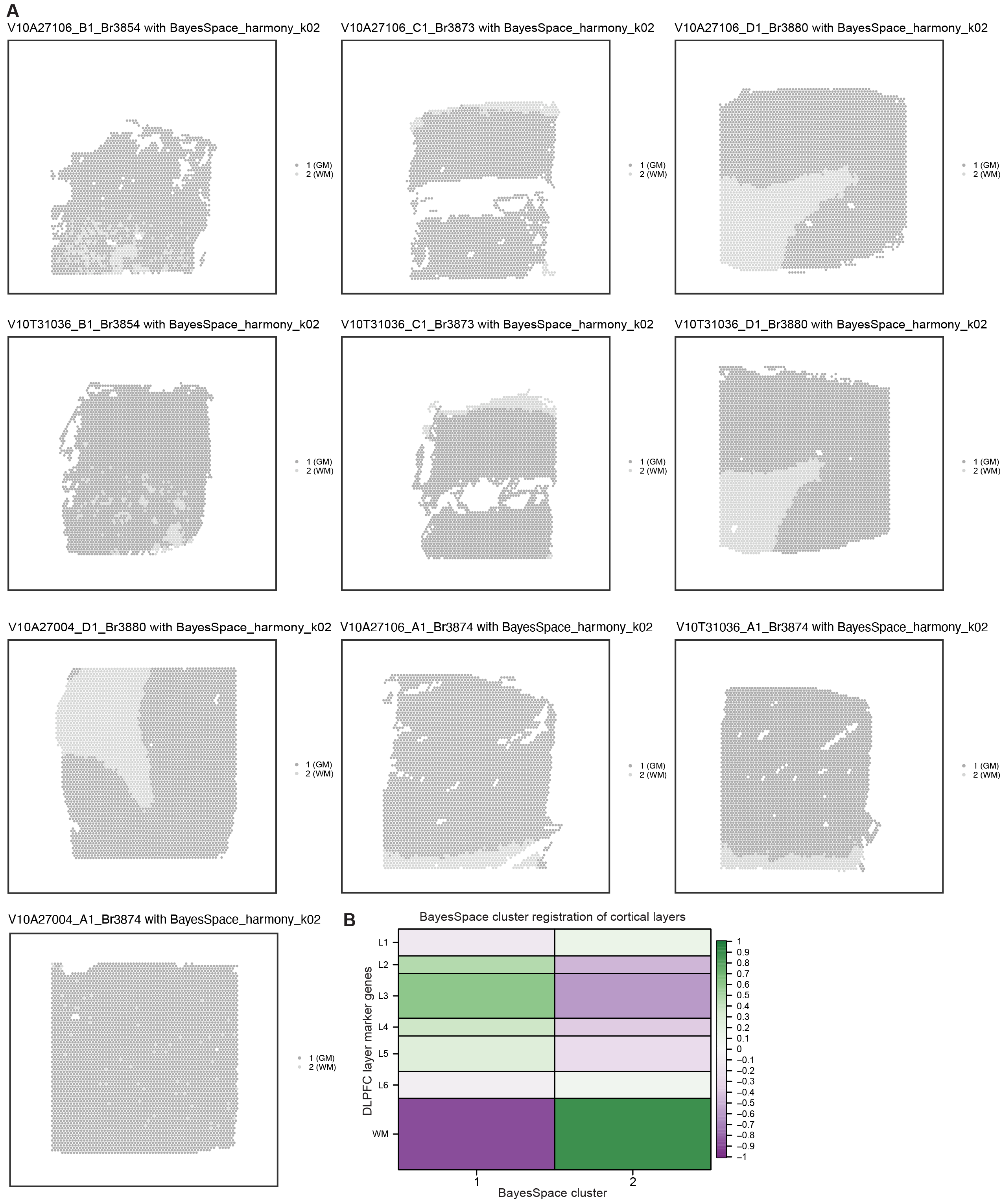

### FigureS12_GOKEGG_Ab_env_300ppi.png

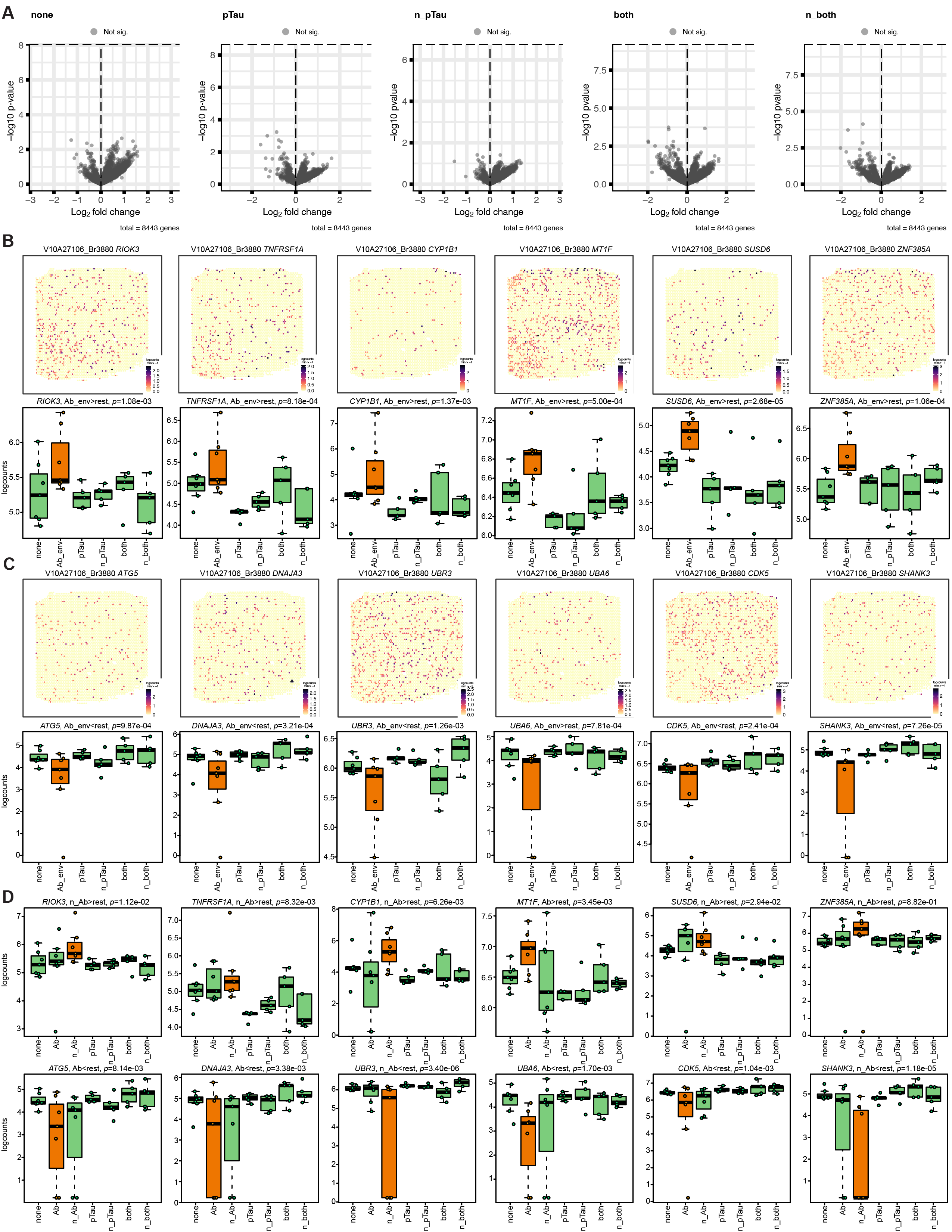

### FigureS13_GSEA_300ppi.png

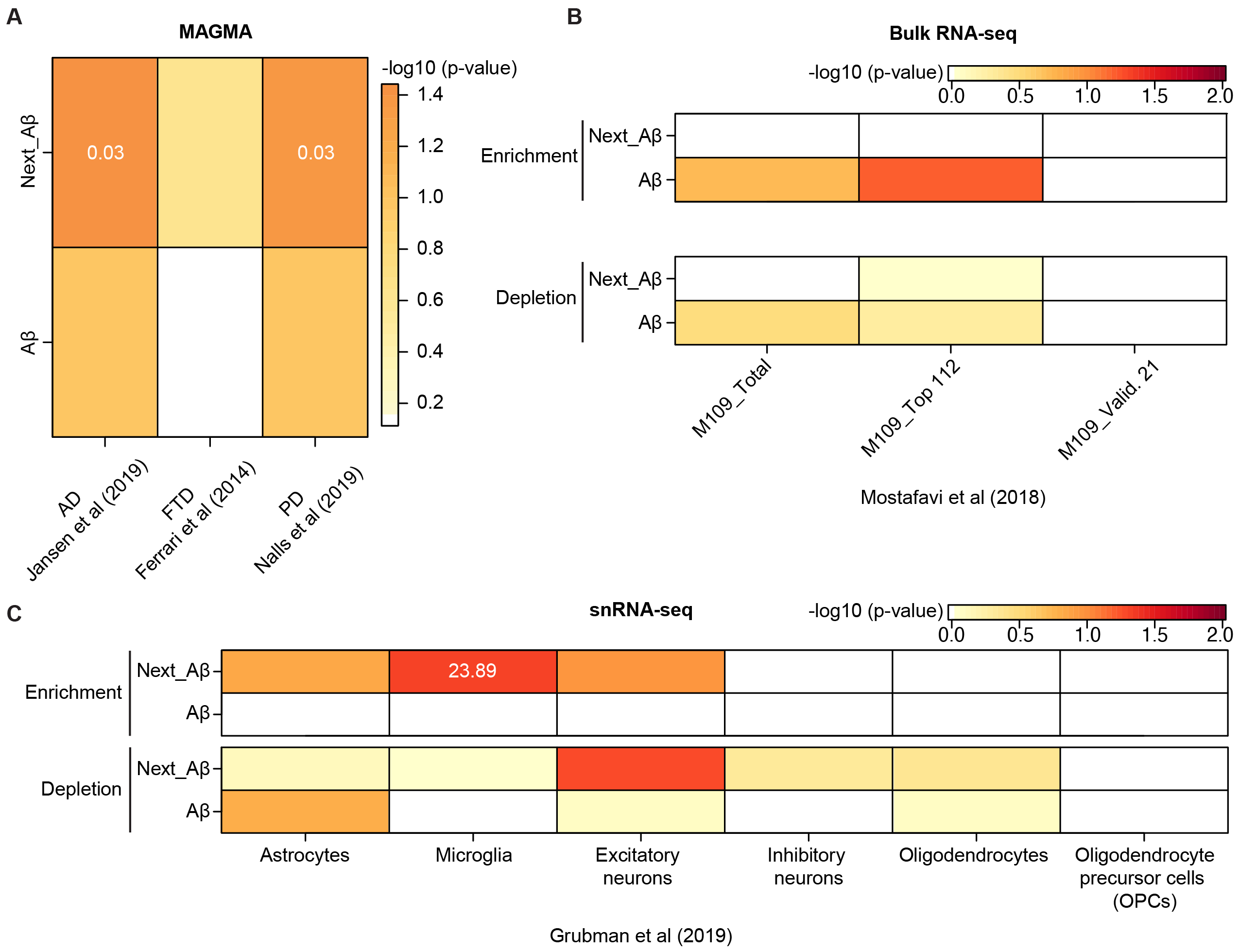

### FigureS14_SpEnrichment_300ppi.png

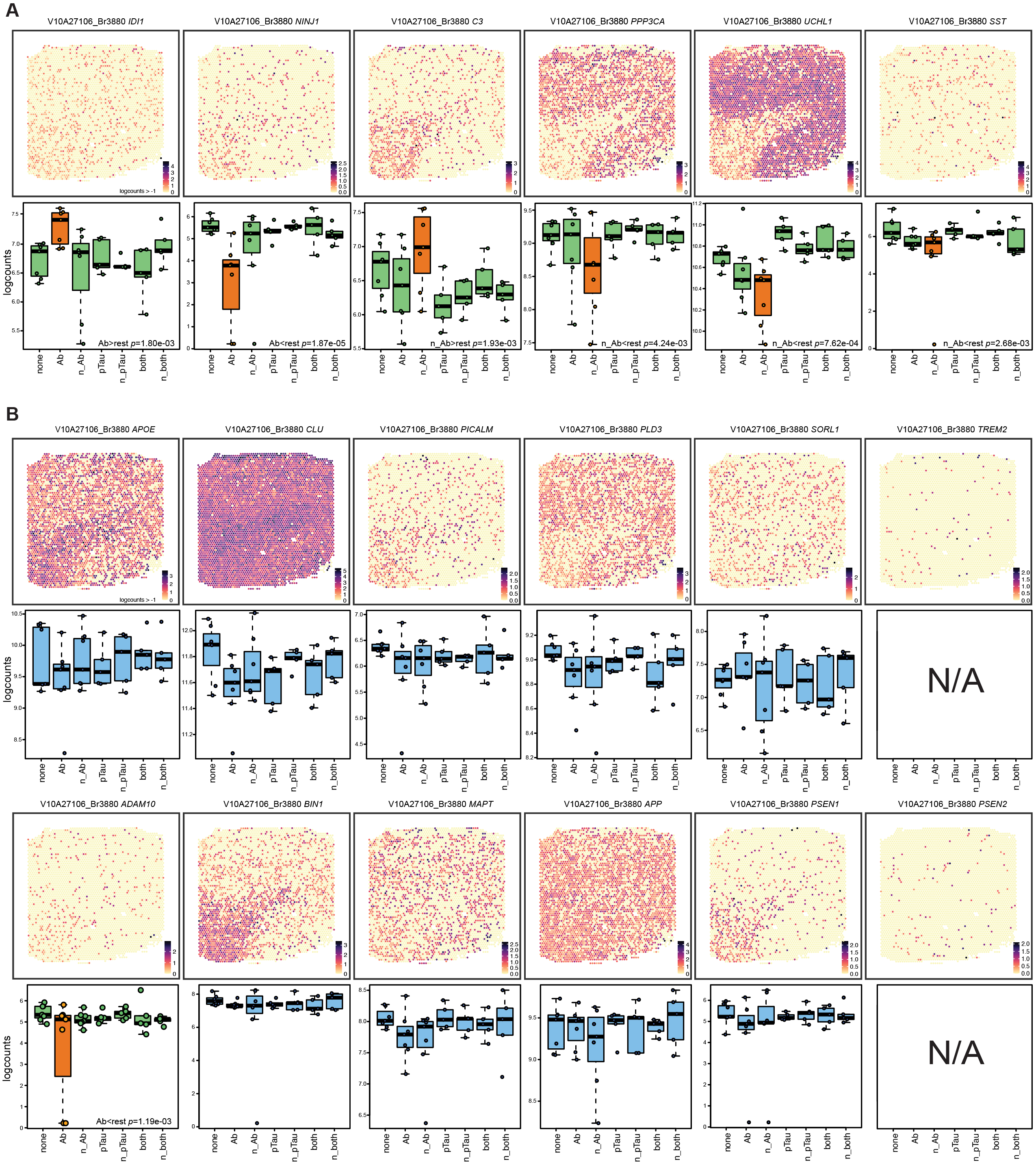

### FigureS15_RFISH-IF_300ppi.png

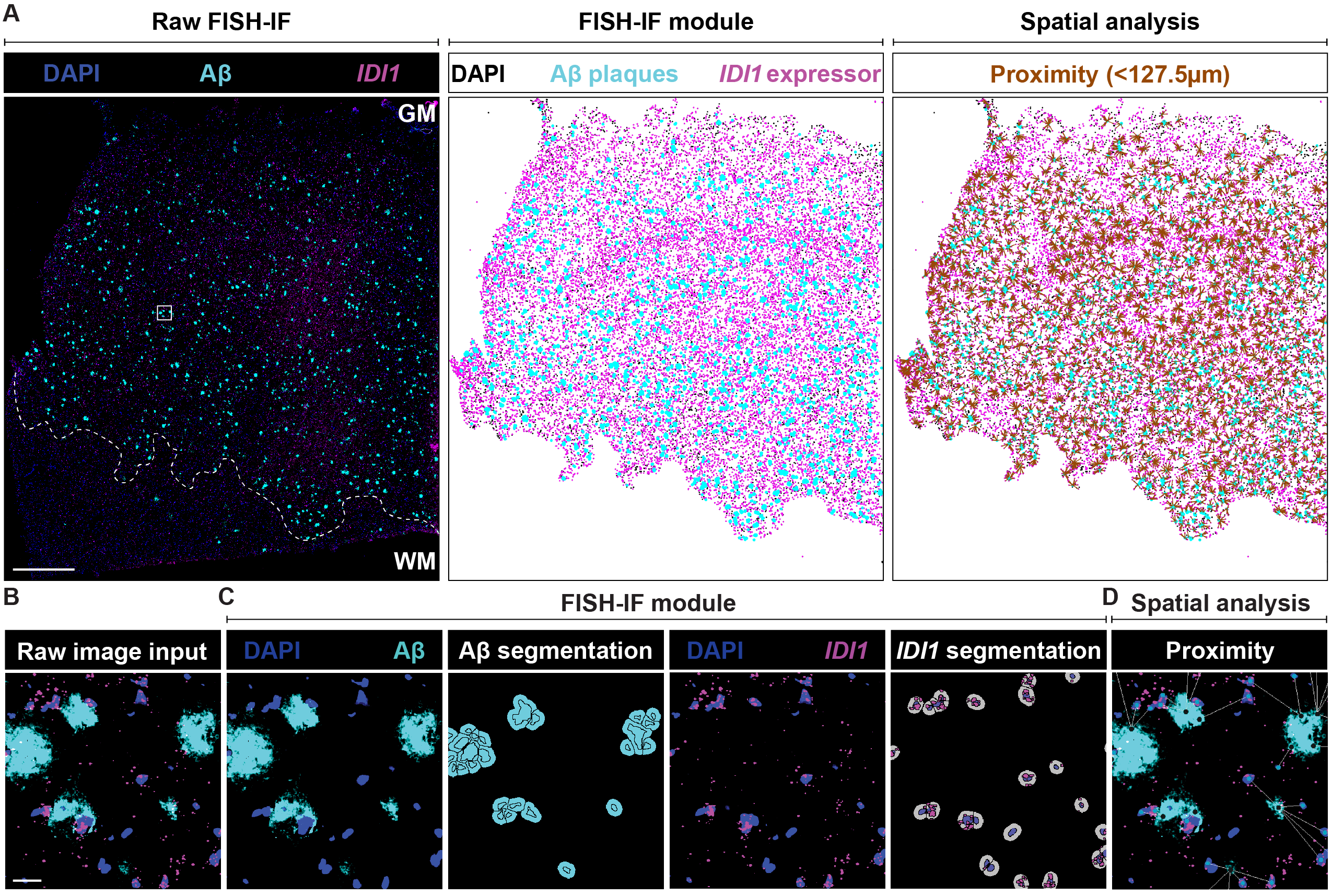

### FigureS16_RFISH-IF_2_300ppi.png

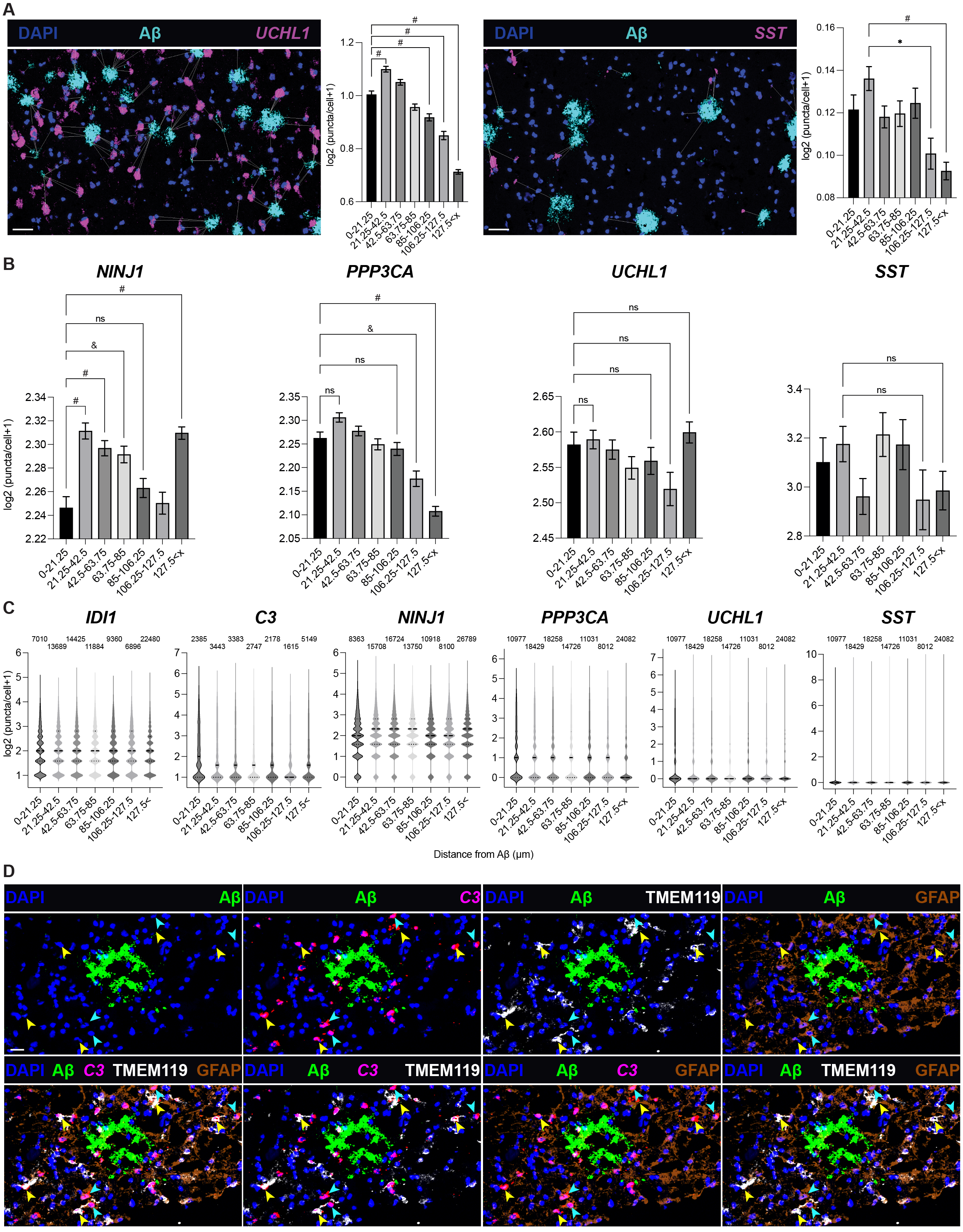
